# Genomic diversification of dehydrin gene family in vascular plants: three distinctive orthologue groups and a novel KS-dehydrin conserved protein motif

**DOI:** 10.1101/2021.04.21.440853

**Authors:** Alejandra E. Melgar, Alicia M. Zelada

## Abstract

Dehydrins (DHNs) are a family of plant proteins that play important roles on abiotic stress tolerance and seed development. They are classified into five structural subgroups: K-, SK-, YK-, YSK-, and KS-DHNs, according to the presence of conserved motifs named K-, Y- and S-segments.We carried out a comparative structural and phylogenetic analysis of these proteins, focusing on the less-studied KS-type DHNs. A search for conserved motifs in DHNs from 56 plant genomes revealed that KS-DHNs possess a unique and highly conserved N-terminal, 15-residue amino acid motif not previously described. This novel motif, that we named H-segment, is present in DHNs of angiosperms, gymnosperms and lycophytes, suggesting that HKS-DHNs were present in the first vascular plants. Phylogenetic and microsynteny analyses indicate that the five structural subgroups of angiosperm DHNs can be assigned to three groups of orthologue genes, characterized by the presence of the H-, F- or Y-segments. Importantly, the hydrophilin character of DHNs correlate with the phylogenetic origin of the DHNs rather than to the traditional structural subgroups. We propose that angiosperm DHNs can be ultimately subdivided into three orthologous groups, a phylogenetic framework that should help future studies on the evolution and function of this protein family.

## Introduction

Plants have to deal with different environmental stresses that can negatively affect their growth and development. Loss of intracelullar water in response to abiotic stresses like drought, salinity and low temperature results in the accummulation of Late Embryogenesis Abundant (LEA) proteins in different vegetative tissues. These proteins, which belong to several different families, were first identified in cotton seeds as proteins upregulated during a programmed maturation drying event during seed development^1,2^. LEA proteins belong to a large group of proteins known as “hydrophilins” characterized by glycine-rich, highly hydrofilic disordered amino acid sequences^3^. Based on sequence similarity, LEAs are classified into 7 families distinguished by the presence of different conserved motifs^4,5^.

Dehydrins (DHNs) constitute a biochemically and evolutionarily distinct group of LEAs with a highly modular structure consisting of a combination of different conserved motifs, variable in number and type, interspersed within weakly conserved amino acid segments. The presence of at least one conserved lysine rich-motif, named the K-segment, is usually used as a *sine qua non* condition to define a protein as a dehydrin^6^. Two other conserved motifs have been described, the Y- and S-segments, that in conjuction with the K-segment are the basis for the general classification of DHNs into 5 structural subgroups: KnS, SKn, YnK, YnSKn and Kn-DHNs, where n refers to the number of repetitions of a given motif^7^.

The Y-segment, whose conserved consensus sequence is [V/T]D[E/Q]YGNP, is usually located at the N-terminus of the protein in one or several tandem copies, while the S-segment, a tract of Ser combined with Asp and Glu residues, is always found in one copy per protein^7^. Recently, Strimbeck^8^ described a new conserved motif present in a subgroup of SK-DHNs that consists of a 11-residue amino acid consensus sequence (DRGLFDFLGKK), named the F-segment. These conserved motifs are usually surrounded by less conserved sequences denoted Phi-segments, characterized by a higher proportion of Gly, Thr, and Glu residues.

Several studies have identified, classified and determined the role of DHNs in different plants. A positive relationship between the level of DHN transcripts and/or protein accumulation and plant stress tolerance has been reported^9–11^. Furthermore, it has been observed that DHN overexpression in transgenic lines increases resistance to unfavourable environmental conditions, such as cold, drought and salinity^12–14^. In vitro experimental evidence from biochemical assays and localization experiments suggests multiple roles for dehydrins, including membrane protection, cryoprotection of enzymes, interaction with DNA and protection from reactive oxygen species^15–17^.

In most of these studies, the biochemical and functional characteristics of these proteins were analysed within the framework of conserved structural domains, but a comparative analysis taking an evolutionary point of view has not been fully explored. The phylogenetic relationships of DHNs have been studied in many different plants, but most of these studies are limited to one genus or species^18–20^. Only recently, a comprehensive understanding of the evolutionary history of DHNs has been attempted. A phylogenetic and structural analysis of a large number of plant DHNs by Riley et al (2019) suggests that the ancestral DHN belonged to a Kn or SKn group, and that YSKn and YKn-DHNs first arose in angiosperms^21^. On the other hand, Artur (2019) showed that angiosperm DHNs with Y- and F-segments belong to two different orthologue groups that can be distinguished by synteny conservation across angiosperms^22^. The evolutionary origin of KS-DHNs is still elusive, since previous works have neglected this group.

Here, we present a thorough phylogenetic and structural analysis of DHNs obtained from a wide spectrum of plant genomes. Even though KS-DHNs have previously been described only in a handful of species, we show that this DHN group is actually present in all angiosperms as well as in gymnosperms and lycophytes, indicating its ancient origin in vascular plant evolution. We show that KS-dehydrin genes share a conserved synteny neighbourhood in angiosperm genomes and possess a conserved N-terminal domain, that we named H-segment, and propose that all angiosperm DHNs belong to one of three orthologue groups, the H, F and Y groups. We also carried out a comparative analysis of the different domain structures and biochemical characteristics inherent to the hydrophilin quality of DHNS to investigate how they correlate with their evolutionary origin.

## Methods

### Dehydrin protein sequences database construction

Initially, DHN proteins were obtained by searching plant genomes or transcriptomes with the Hidden Markov Model (HMM) profile asigned to the DHN protein family (PF0027), downloaded from the Pfam database (http://pfam.xfam.org/), using the HMMER 3.1 software (http://hmmer.org/). The HMM profile was used to search the Phytozome v13 database (https://phytozome-next.jgi.doe.gov/) which harbours 56 genomes from species spanning the whole viridiplantae clade, including one rodophyte, nine chlorophytes, two briophytes (*Ceratodon purpureus* and *Physcomitrella patens*), the lychophyte *Selaginella moellendorffii*, the angiosperm basal species *Amborella trichopoda* and *Nymphaea colorata* and a subset of 9 monocots and 28 eudicots representing different families. To include gymnosperm species in our search, we employed the Gymno PLAZA 1.0 database^23^ and the ConGeniE database (http://congenie.org/) which contain the transcriptomes of *Ginkgo biloba, Picea abies* and *Picea glauca*.

To identify the conserved motif structures of DHN proteins, we used the MEME software (http://meme-suite.org/)^24^. Since we noticed that KS-type DHNs were underrepresented in this preliminary DHN database, we searched the National Center for Biotechnology Information (NCBI) database to retrieve homologues of *Arabidopsis thaliana* HIRD11 using the Blastp algorithm, and the new KS-DHN sequences were used to construct a HMM profile specific for this DHN group. In parallel, HMM profiles were also constructed for F- and Y-DHNs. Finally, the three HMM profiles were used to reanalise the databases. All DHN sequences identified in this work are shown in Supplementary Table S1.

### MEME searching conserved motif in DHNs database

The conserved motif structures of DHN proteins were identified using MEME software to find recurrent ungapped motifs assuming that each sequence may contain any number of non-overlapping motifs. The results presented correspond to an analysis made with the following parameters: number of motifs = 8, motif width = 6 to 20, and number of sites for each motif = 2 to 600 (Supplementary Figure S1). The E-values of the different motifs predicted by MEME for our DHN database were compared to E-values calculated from the same sequenced randomly shuffled using the same MEME run parameters to confirm the significance of the discovered motifs.

### Multiple sequence alignments and phylogenetic tree construction

In order to establish orthology/paralogy relationships among the sequences, phylogenetic relationships within each DHN family were estimated. The DHN protein sequences were aligned using Clustal Omega^25^ or T-coffee^26^ with default parameters, and multiple sequence alignments (MSA) were visualized using Jalview^27^. The phylogenetic tree was constructed using an MSA that included only angiosperm DHNs, in order to prevent very divergent sequences from reducing the quality of the alignment. Phylogenetic trees were estimated by the Maximum Likelihood (ML) method as implemented in the NGPhylogeny website (https://ngphylogeny.fr/)^28^ using FastTree^29^ with the LG amino acid substitution model^30^ and the GAMMA model with invariant sites for rate heterogeneity. A total of one thousand bootstrap samplings were run. The resulting tree was visualized using iTOL^31^.

### Microsynteny analysis

For microsynteny analysis of selected DHN genes, the corresponding proteins were identified in the NCBI database by pairwise BLASTP searches. Annotations with 100% identity were selected and the genomic context analysed using the NCBI Genome Data Viewer (GDV). Protein sequences of ten to twenty genes flanking both sides of DHN genes were compared between species, using the loci of *A. trichopoda* DHN genes as references. Reciprocal BLASTP analysis were used to confirm homology, with sequences that matched with an E-value of <10−5 being considered homologous to each other.

### Estimation of physicochemical properties of DHNs protein

The theoretical physicochemical properties of DHNs such as grand average hydropathicity index (GRAVY), molecular weight (MW), isoelectric point (pI) and glycine percentage were calculated with the ProtParam tool of Expasy (https://web.expasy.org/protparam/). The GRAVY index indicates the hydrophobicity of the protein and was calculated as the sum of the hydropathy values (Kyte and Doolittle parameters) of all amino acids divided by the sequence length. Proteins with positive GRAVY scores are hydrophobic whereas proteins with negative GRAVY scores are hydrophilic. The fold index of proteins was estimated using the FoldIndex© software (https://fold.weizmann.ac.il/fldbin/findex).

## Results

### Unbiased genome-wide identification of dehydrins in Viridiplantae genomes

As a first step to understand the evolutionay history of KS-DHNs and their relationship to the other structural subgroups (YnSKn-, YnKn-SKn- and Kn-DHNs), we performed a genome-wide sequence homology search to identify the complete repertoires of DHNs across 56 genomes of species belonging to the Viridiplantae clade, including representative members of chlorophytes (green algae) and streptophytes (see Materials and Methods). The initial screening was made using a Hidden Markov Model (HMM) profile defined for dehydrin family proteins (Pfam2057) obtained from the Pfam 33.1 database^32^. Surprisingly, when we analised the sequences retrieved, we noticed that well known KS-DHNs, such as the HIRD11 dehydrin from the dicot *Arabidopsis thaliana* (At1g54410)^33^ and the ZmDHN13 from the monocot *Zea mays*^13^ were not detected by the algorithm. That prompted us to hypothesize that a Pfam00257-based HMM is not sensitive enough to recognize KS-DHNs as members of the dehydrin family. To overcome this limitation we built three different HMM profiles: one (KS-HMM) using KS-DHN sequences from angiosperm genomes identified by Blastp searches using *A. thaliana* HIRD11, and two other profiles (F-HMM and Y-HMM) based on angiosperm proteins belonging to the F- and Y-DHNs orthologous groups recently described^22^.

After searching with Pfam2057 and the three DHN group-specific HMM profiles, we recovered a total of 305 non-redundant DHN sequences from genomes of representative species of briophytes (4), lycophytes (1), gymnosperms (3) and angiosperms (36) (Supplementary Table 1). No sequences were retrieved from the 9 chlorophyta green algaes genomes analysed, neither from the genome of the streptophyte alga *Chara brunii*, that belongs to a sister group to embryophytes^34^, confirming that the DHNs family emerged in land plants^35^.

Remarkably, the KS-HMM profile displayed an increased sensitivity in recognising DHN homologues, since it was able to identify 92.2% of DHN proteins, while the Pfam00257-based HMM identified 81.7% and the other two HMM profiles only 83% of DHN sequences (Fig. 1). Among a total of 62 DHNs exhibiting the KS-architecture, only 17 could be retrieved using the Pfam00257 profile, confirming its poor performance in recognizing KS-DHNs. While F-HMM has a better perfomance and could recognize 26 KS-DHNs, only the KS-HMM profile was able to retrieve all KS-DHNs. Indeed, 32 KS-DHNs could only be retrieved using the KS-HMM profile. Conversely, the KS-HMM profile failed to recognize 22 DHN sequences that were identified by the other HMM profiles. No DHNs were retrieved solely by Pfam00257, indicating that group specific-HMM profiles are necessary and sufficient for a thorough search of DHN proteins in angiosperm genomes.

**Figure 1.**
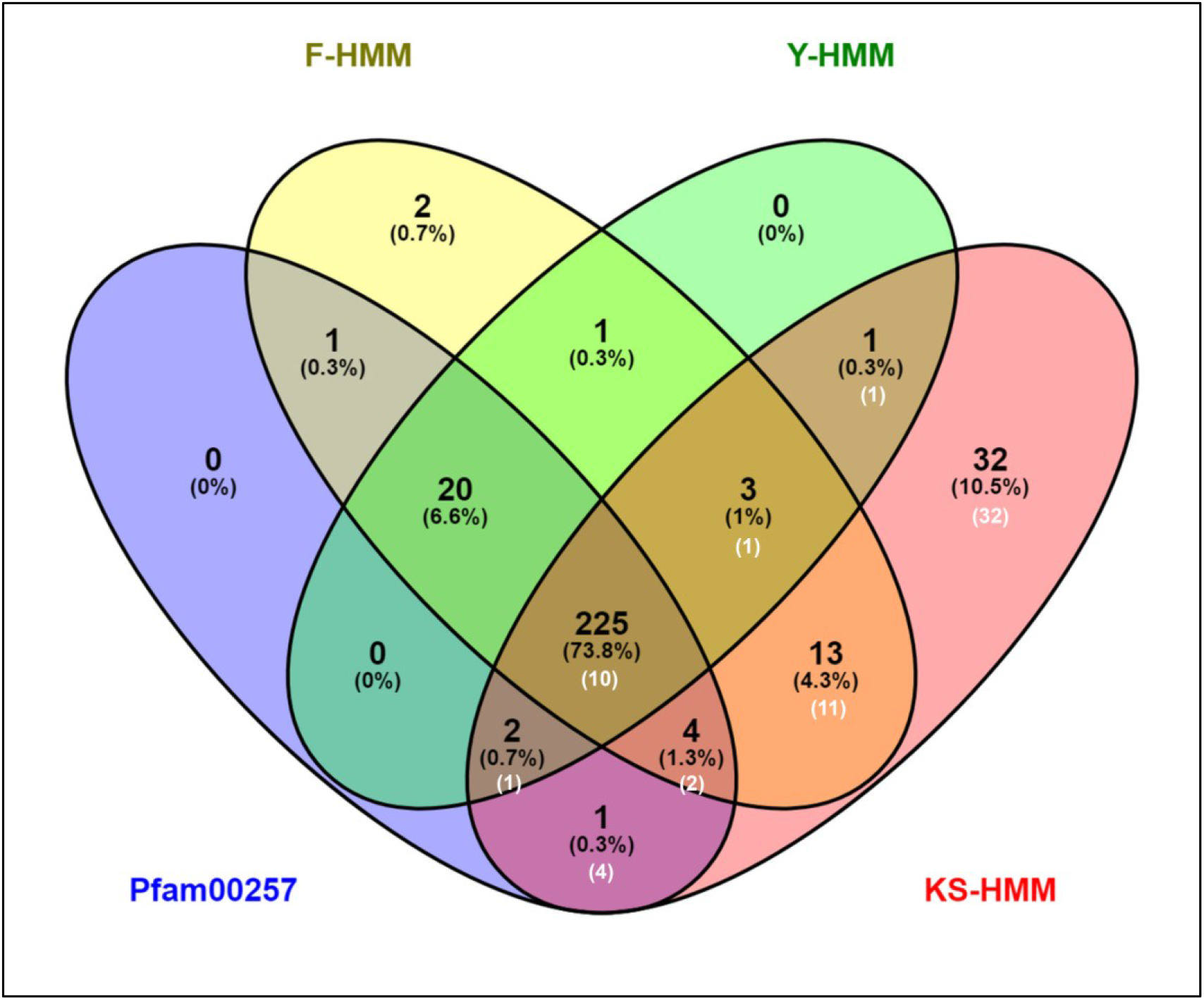
DHNs identified in Viridiplantae genomes by different HMM profiles. Sequences retrieved from species genomes using F-HMM, Y-HMM, KS-HMM profiles constructed in this study and PF00257 the Pfam profile for dehydrin family proteins are displayed as a Venn diagram. White numbers indicate the number of KS-DHNs present in each subset. Note that most KS-DHNs are not recovered using Pfam00257.

### Analysis of conserved protein motif and classification of the dehydrin database

To classify the DHNs of our unbiased database into the structural subgroups, we used the MEME program to check for the presence of known dehydrin motifs (K-Y-F- and S-segments) and to discover putative novel motifs (Supplementary Figure S1). The LOGO representations of the conserved motifs detected and the distribution of DHNs sequences in the different structural subgroups are shown in Figure 2.

**Figure 2.**
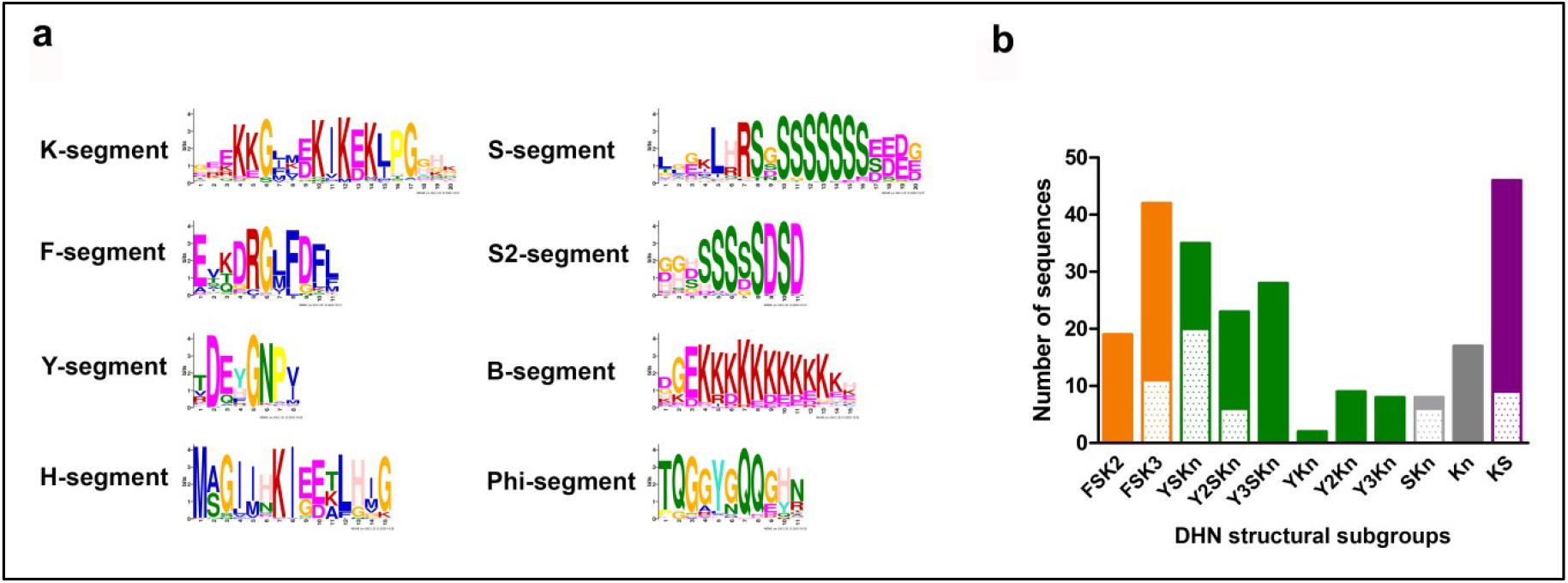
Identification of conserved protein motifs and structural classification of DHNs. (**a**) LOGO representation of the different conserved motifs detected by MEME in the set of DHNs of the unbiased database. (**b**) Number of members of each angiosperm DHN structural subgroup identified in the database. We distinguished FSK2 and FSK3 structural subgroups in accordance to Strimbeck (Strimbeck, 2017). All classified DHNs are listed in Supplementary Table 1. The dotted pattern indicates monocots, while the filled pattern indicates eudicots.

We confirmed the presence of the K-segment in 302 of the 307 dehydrins identified by homology searches based on HMM profiles. The MEME program failed to recognize a sequence similar to K-segment in a few proteins, all of which from non-angiosperm species. However, these proteins all possess a degenerate, less conserved K-segment, as well as other DHN motifs, indicating that they are bona fide DHNs. This is the case, for instance, for DHNs from the lycophyte *S. moellendorffii* and the gymnosperm Ginkgo biloba (see Fig. 3).

**Figure 3.**
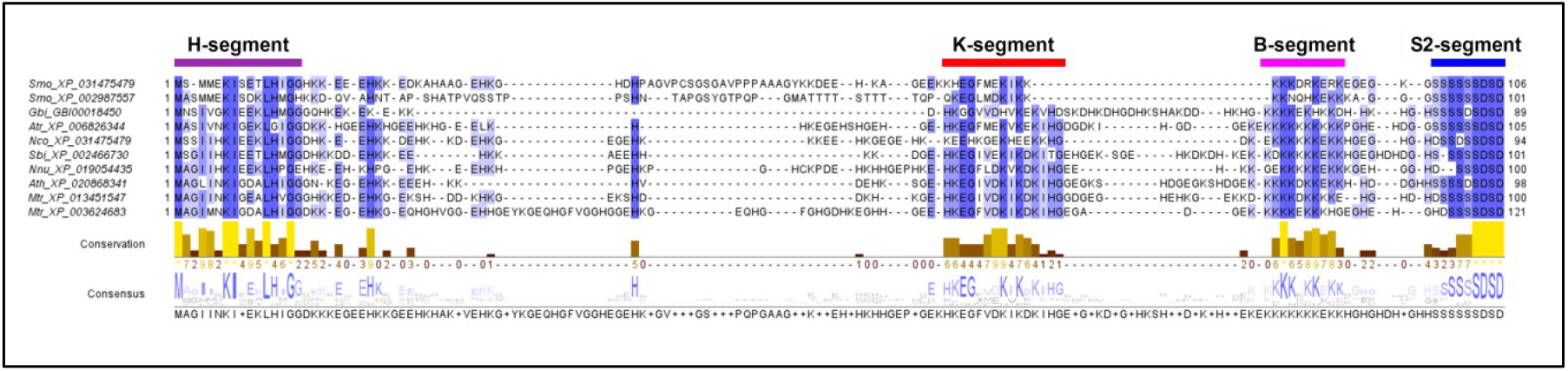
Conservation of KS-DHNs in vascular plants. Multiple sequence alignment of KS-DHNs (HKS-DHNs) of representative angiosperms (*A. trichopoda, S. bicolor, N. nucifera, A. thaliana, M. truncatula)* and non-angiosperms (the lycophyte *S. moellendorffii* and the gymnosperm *G. biloba*) performed with T-Coffee and visualised with Jalview. The H-, K-, B and S-segments are indicated. Note that the general structure of the proteins is conserved in all vascular plant groups.

We identified 75 DHNs bearing a unique F-segment located in the N-terminal region of the proteins, that we classified within the FSKn-DHN structural subgroup. The F-segment predicted with our protein database is similar to the one described by Strimbeck (2017) (Fig. 2A). Importantly, even a search with a specific F-HMM profile failed to identify FSKn-DHNs in the genomes of four bryophyte and one lycophyte species, but we did find them in the three gymnosperms included in this study, *Picea abies, Picea glauca* and *Ginkgo biloba*, confirming that this subtype of DHN probably arose in seed plants. We observed an expansion of the FSKn-DHN gene family in the Pinaceae clade, in accordance with previous observations^36^, but that was not a general feature of gymnosperm species. Only three DNHs were identified in the gymnosperm *Ginkgo biloba*, two of them with a F-segment (FSK2 and FK2) and a third harbouring a novel conserved motif (see below). In angiosperms, the FSKn-DHN subgroup is mostly comprised of FSK2 and FSK3 proteins (Fig. 2B) but, interestingly, only FSK3-DHNs are found in monocots and in the early divergent eudicot *Nelumbo nucifera*, as well as in the basal angiosperms *A. trichopoda* and N. colorata (Supplementary Figure S2), suggesting that FSK2-DHNs might have arisen from an ancestral FSK3-DHNs.

We found a total of 101 DHNs containing one to three copies of the Y-segment per sequence at a N-terminal position, all of them in angiosperms. The mayority of the proteins belong to the YnSKn subgroup, while sequences lacking the S-segment (YnKn) only represent 15%. In monocots we found YSKn and Y2SKn-DHNs only, while Y3SKn- and YnKn-DHNs seem to be restricted to dicots. A motif that resembles a previously sequence defined as the Phi-segment is present only in YSK3- and Y2SK3-DHNs of monocots, as determined by MEME analysis and multiple sequence alignments (Supplementary Figure S4 to Figure S6).

As already mentioned, we identified a total of 62 KS-DHNs in plant genomes, all of which share a novel N-terminal motif (H-segment, see below). Interestingly, the KS-HMM profile allowed us to identify KS-DHNs in the genome of the lycophyte *S. mollendorfii* and the gymnosperm *Ginkgo biloba*, but no KS-DHNs were identified in the genomes of the conifers *P. abies* and *P. glauca*. As can be seen in Figure 3, these proteins present a typical arrangement of KS motifs, with a K-segment followed by a lysine-rich stretch (B-segment) and a S-segment characteristic of this structural subgroup (S2-segment). This is the first time that KS-DHNs are identified in non-angiosperm species and indicates that this kind of dehydrin arose early in land plant evolution.

In angiosperms, all species analysed possessed one or two KS-DHNs genes with the exception of *Glycine max*, with four genes, and two Malpighiales species, *Salix purpurea* and *Populus trichocarpa*, with six and three KS-DHNs, respectively. These Malpigiales proteins are unique between KS-DHNs, since they contain multiple K-segment repeats interspersed with glycine-rich sequences (Phi-segment) and the S2-segment is absent (Supplementary Figure S8).

Concerning the Kn- and SKn-DHN structural subgroups, their representation in vascular plants was minor and, ultimately, they are phylogenetically related to other DHN structural subgroups, as will be discussed later (see below). In contrast, most of the eighteen non-vascular DHNs that we identified in the genomes of the mosses *P. patens* (six proteins), *Ceratodon purpureus* (six), *Sphagnum fallax* (three) and the liverwort *Marchantia polymorpha* (three) belong to the Kn-structural subgroup. The exception is an atypical DHN containing a series of repetitive motifs resembling the Y-segment in the N-terminus that is present in *P. patens* (PpDHNA)^37^ and *C. purpureus* (Supplementary Figure S10). A phylogenetic analysis indicates the presence of five DHN orthologue groups in *P. patens* and *C. purpureus* (Supplementary Figure S9), which reflects the phylogenetic proximity of the Funariidae and Dicranidae clades^38^. The DHNs from the more distantly related *S. fallax* (Sphagnophytina) did not cluster with the DHNs of the other mosses, but multiple sequence alignments and reciprocal Blastp analyses suggest that two of the *S. fallax* DHNs (Sphfalx0010s0103.1 and Sphfalx0064s0013.1) are related to groups III and V of *P. patens* and *C. purpureus* (Supplementary Figure S10). The three DHN proteins of the liverwort *M. polymorpha* do not display any obvious homology to moss DHN groups outside the K-segment.

### The H-segment is a novel conserved motif present in all KS-dehydrins

Our MEME analysis identified a highly conserved motif, not previously described, at the N-terminal region of all angiosperm KS-DHNs analysed. This 15-residue segment is characterized by a combination of hydrophobic amino acids Ile and Leu with amphipathic amino acids Lys and Glu, framed by two Gly at positions 3 and 15 conserved in 91% and 87% KS-DHNs (Fig. 2). In addition, the high percentage of conservation of the Lys (97%) and Ile (97%) located in the central positions 7 and 8 strongly suggest an important function in KS-DHNs. Ile residues at positions 4 and 5 are less conserved, and are often replaced by other hydrofobic amino acids like Phe, Val or Met. KS-DHNs are characterized by sequences enriched in His amino acids, as reflected in the name HIRD11 for the *A. thaliana* KS-DHN, which stands for <u>Hi</u>stidine-<u>R</u>ich <u>D</u>omain 11 kDa protein^39^. Two His residues are found in positions 6 and 13 in 56% and 77% of KS-DHNs, respectively. Since this novel motif seems to be a signature of KS-DHNs, we propose to name it the H-segment, reflecting the particular feature of these kind of proteins, even when histidines are not the most prevalent aminoacids in the motif.

A structural prediction of representative angiosperms KS-DHNs, obtained by Phyre2^40^, indicates with high confidence the presence of a helical α-helix spanning the H-motif in all proteins analysed (Fig. 4). This helical wheel projection is conformed by the alternation of hydrophobic and hydrophilic amino acids and is surrounded by highly conserved Gly amino acids that could function as a helix breaker due to their high conformational flexibilty, which makes it entropically expensive to adopt the relatively constrained α-helical structure. A very similar structure is predicted for the K-segment, suggesting that the H-segment could also have amphiphilic membrane or protein binding properties as described for the K-segment^15,41^.

**Figure 4.**
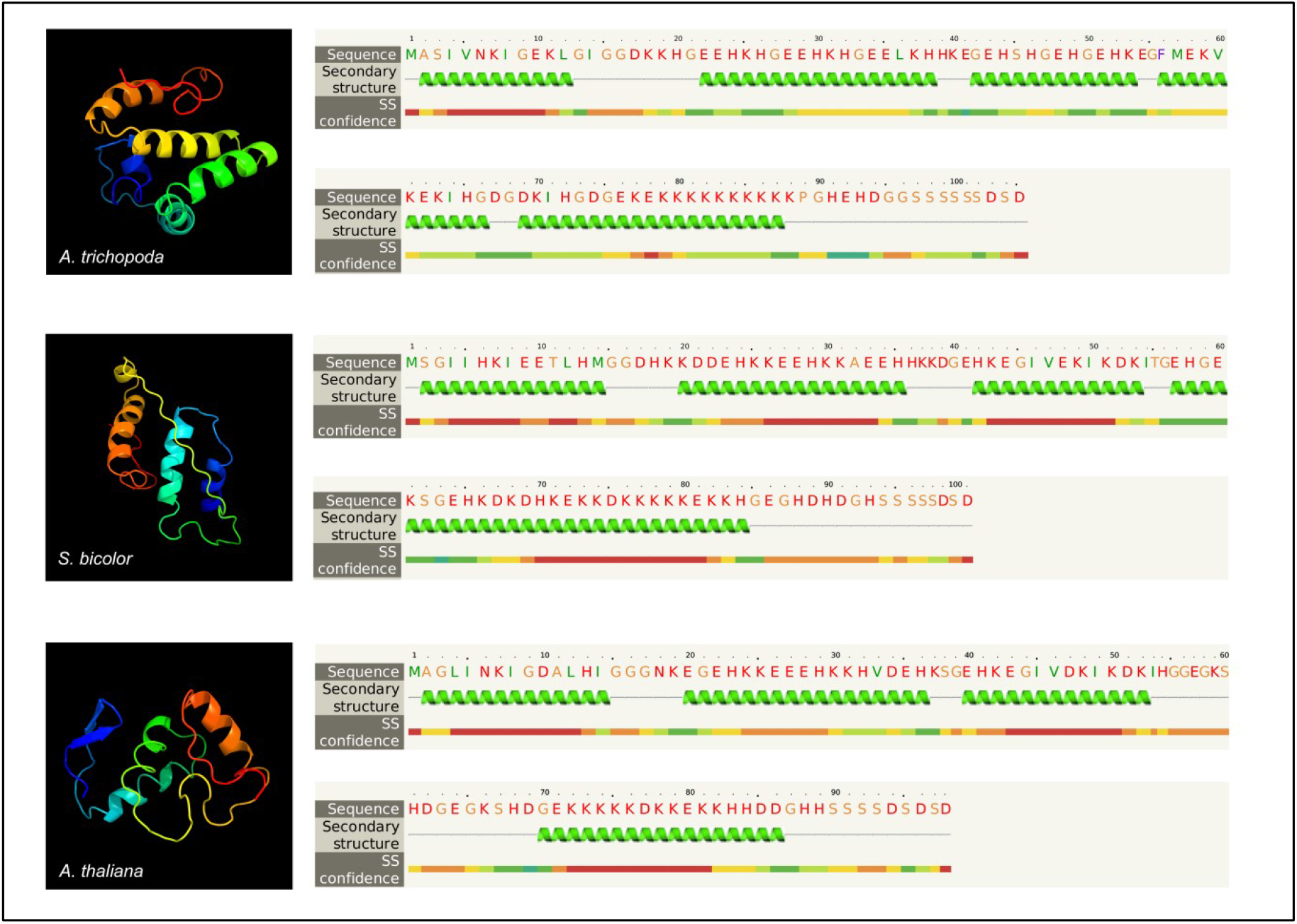
Predicted H-DHN structures. The secondary structure of H-DHN proteins from *A. trichopoda, S. bicolor* and *A. thaliana* (HIRD11) were predicted by the Phyre2 program. Note that a helical α-helix is predicted to be present near the N-terminus of the proteins, spanning the H-segment. Red colour indicates high confidence in the prediction.

The K- and S-segments of KS-DHNs present some particular characteristics compared to FSK and YSK-DHNs. The prevalence of amino acids at positions 6, 16 and 17 differs in the K-segments of KS-DHNs (Fig. 2 and Supplementary Figure S7). Thus, position 6 is occupied by an Asp in all KS-DHNs instead of the Lys that it is typically present in DHNs, with the exception of three FSK-dehydrins of the gymnosperm *P. abies*. Concerning position 16, KS-DHNs usually have an Ile instead of a Leu. It is notable that in the species with more KS-DHN genes, proteins with one or the other amino acid in this position can be found (see Supplementary Table 1; *Solanum tuberosum*; *Populus trichocharpa, Phaseolus vulgaris*). Even though Ile and Leu amino acids are generally considered conservative, there is evidence that these amino acids are not always interchangeable, affecting the affinity and specificity of protein-protein and protein-membrane interactions^42^, which might potentially lead to functional diversification of the KS-DHNs by modulating K-segment behaviour. In contrast to other DHNs, KS-dehydrins do not show a clear prevalence of Pro at position 17; instead, His is the most frequent amino acid at this position, while Pro is only found in the DHNs of Rutaceae and in a subgroup of Malpighiales species, and Thr is prevalent in monocot KS-DHNs at this position. The capacity to tolerate different kinds of amino acids at that position could indicate that it is not essential for K-segment functionality. In spite of these differences, the predicted α-helix structure delimited by conserved Gly amino acids of the K-segment is conserved in KS-DHNs (Fig. 4).

As for S-segments, which are characterised by a stretch of Ser residues, there are differences in the length of the Ser-amino acid stretch and neighbouring amino acids between KS-DHNs and other DHNs. We observed that the core of 6 to 9 Ser residues usually ended with negatively-charged Asp or Glu amino acids in all structural subgroups of DHNs. On the other hand, the triad Leu-His-Arg that precedes the Ser stretch, which is highly conserved in all FSK-DHNs and in the mayority of YSK-DHNs, is not found in KS-DHNs. Figure 2A shows the S-segment consensus for FSK and YSK-DHNs (segment S1) and the one found in KS-DHNs (segment S2, see also the alignments in Supplementary Figure S2 and Figure S7). The S-segment of all types of DHNs has been shown to be a hotspot for phosphorylation by kinases^13,43,44^, and the differences between the S1- and S2-segment could result in different kinase specificities. For instance, the triad Leu-His-Arg constitutes part of the recognition sequence for SnRK2 kinases^45^, which have been recently demonstrated to phosphorylate *A. thaliana* dehydrins ERD4 and ERD10 in response to osmotic stress^46^.

A 11-residue Lys-rich motif has been consistently detected in all KS-DHNs as well as in all FSK2 and the majority of FSK3-DHNs (Supplementary Figure S1 to Figure S3; Fig. 5). The whole motif comprises 9-11 Lys residues preceded by Gly and Asp amino acids in positions 2 and 3 (Fig. 2). In KS- and FSK-DHNs, the Lys-rich motif is located between the S-segment and the K-segment while, at the same position, YK-and YSK-DHNs usually have a RRKK or RRKKK sequence framed by Gly residues, a motif that resembles monopartite nuclear localization signals^47,48^. It has been demonstrated that monopartite NLS require specific residues flanking the core basic cluster for their complete activity, and in particular the inclusion of Asp- and Glu-aminoacid seems to be detrimental for its activity^48^. Some Lys-rich motifs could constitute a NLS sequence, but the presence of Asp or Glu amino acids at positions 2 and 8 in KS- and FSK-DHNs suggests that the conservation of the Lys-rich motif could fulfill a distinct funtion. In conclusion, the KS-DHNs can be better described as having a H-K-S structural organization, with H being a newly described segment, exclusively present in this group of DHNs.

**Figure 5.**
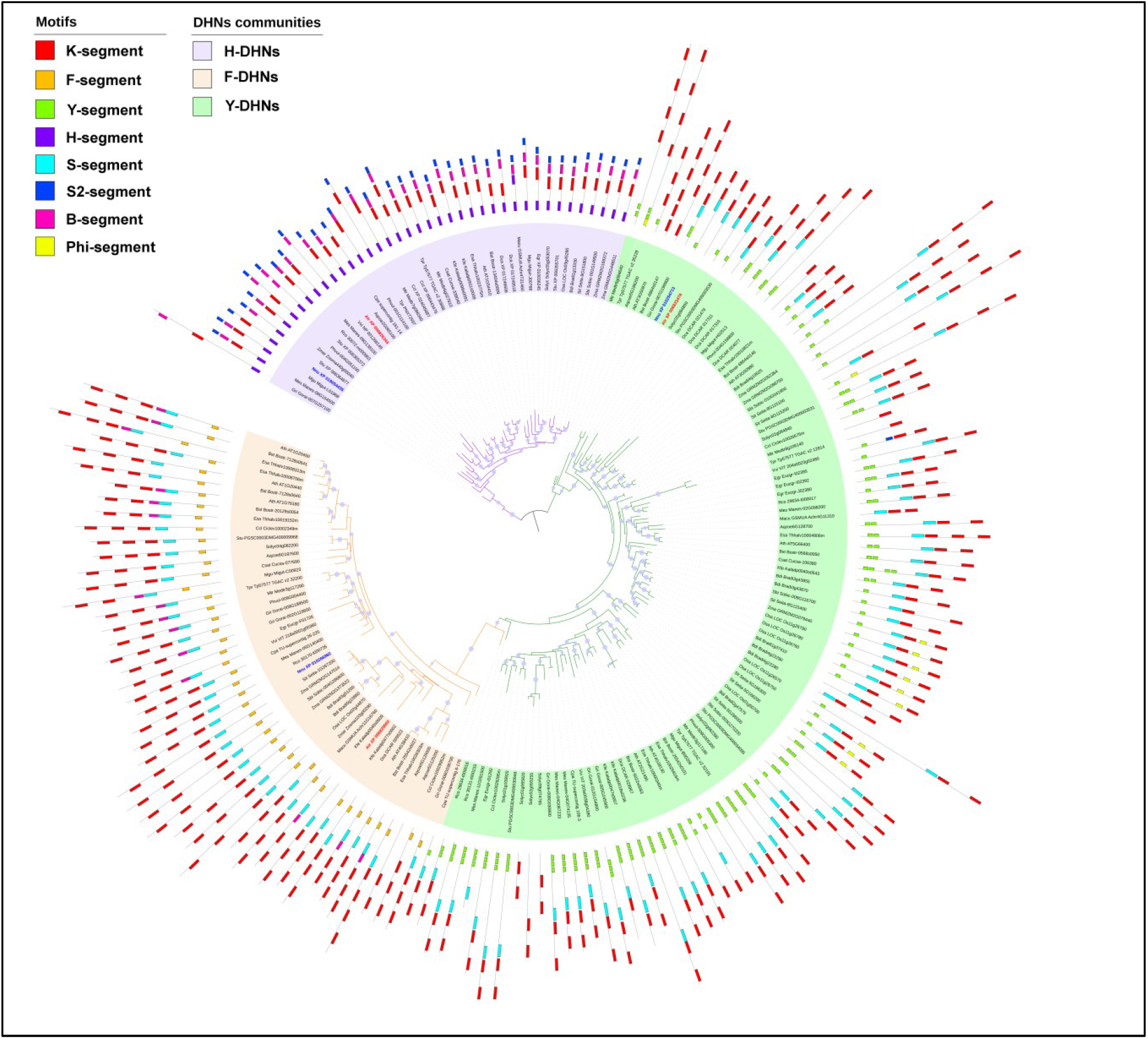
Phylogenetic tree of DHN proteins. Amino acid sequences of DHN proteins from angiosperms were aligned and an approximately maximum-likelihood reconstruction of the phylogenetic relationships was generated using FastTree 2 (Price et al. 2010). Nodes with bootstrap support over 95% are indicated by a violet dot. The motifs of each protein are indicated by coloured boxes, as indicated. Note that all H-DHNs are grouped together in a branch with high support, and most F- and Y-DHNs are also grouped together, forming three groups. DHN proteins from the basal angiosperm A. trichopoda (red) and the basal eudicot N. nucifera (blue) are indicated to show that they possess one DHN in each group.

### Phylogenetic analysis reveals three basic groups of DHNs in angiosperms

Having identified the motifs that characterize the KS-structural subgroup, we sought to infer the phylogenetic relationships between the angiosperm DHN protein sequences that we identified. We decided to use only angiosperm DHNs to build a phylogenetic tree due to the sparse taxonomic sampling of other land plant DHNs. We used the approximately-maximum-likelihood principle as implemented in FastTree 2^49^ and estimated statistical robustness with the bootstrap method.

The resulting tree is roughly organised in three branches or groups (Fig. 5). All KS-DHNs, characterised by the presence of the H-segment (H-DHNs), are grouped together in a branch with high bootstrap support (96%). The other DHNs are separated into two branches, one harbouring most DHNs that contain the F-segment (FSKn), while the other contains DHNs carrying the Y-segment (YnSKn, YnKn). Interestingly, the few DHNs that contain only the K-segment or a combination of K- and S-segments are placed within the F or Y branches, indicating that these DHNs actually belong to either of these groups. Overall, the phylogenetic tree suggests that each angiosperm DHN belongs to one of three phylogenetic families, basically distinguished by the presence of the H-, F- or Y-segments. This conclusion is reinforced by the observation of the DHNs of plants at key phylogenetic positions. Thus, the basal angiosperm *Amborella trichopoda* possesses three DHNs, each one belonging to the H, F or Y groups (Fig. 5). Similarly, the three DHNs from the basal dicot *Nelumbo nucifera* are also each one placed into the H, F and Y groups. Overall, the phylogenetic results suggest that these three groups of DHNs were present since the begining of angiosperm evolution.

### H-DHNs belong to a separate synteny community in angiosperms

Even though the phylogenetic tree described above separates angiosperm DHNs into three groups, the large number of different motifs and their divergent arrangement in DHNs makes the sequences difficult to align and reduces the certainty of the phylogenetic reconstruction. Since synteny analyses of orthologue genes can give important hints about the evolution of genomes and gene families^50^, we performed an analysis of the genomic neighbourhood (microsynteny) of DHN genes in order to reinforce our phylogenetic results.

Recently, Artur et al. (2018) analysed DHN genes plant genomes and identified two main synteny blocks (or communities) among angiosperms, corresponding to DHNs containing the F (community 1) and Y (community 2) motifs^22^. Their analysis, however, did not include DHNs of the KS group, presumably due to the difficulty of retrieving these sequences using the Pfam motif PF00257. To verify whether DHNs containing the H-segment would also be part of a syntenic community, we compared 40 genes surrounding the unique H-DHN of the basal angiosperm, *Amborella trichopoda* to the genomic neighbourhoods of H-DHN loci of the waterlily *Nymphaea colorata* ^51^, the basal eudicot sacred locus, *Nelumbo nucifera* ^52^, the legume *Medicago truncatula* ^53^, the model plant *A. thaliana* (HIRD11)^54^ and the monocot grass *Sorghum bicolor* ^55^. All of these species possess only one H-dehydrin paralogue except for *M. truncatula*, which has two (Supplementary Table 1). We chose *A. trichopoda* as the basis for synteny comparison since this flowering plant belongs to a sister lineage to all other angiosperms (Amborellales), did not undergo the whole genome duplications that affected other lineages and its genome exhibits conserved synteny with other angiosperms, features that facilitate the study of gene family evolution in plants^56,57^. As shown in Figure 6, 17 genes that surround the H-DHN of *A. trichopoda* (LOC18421535) are also present around the H-DHN gene of *N. colorata*, which belongs to a group, the Nymphaeales, that is a sister lineage to all angiosperms except for Amborellales^58^. A smaller number of conserved genes are present around the H-DHN genes from the eudicots *N. nucifera, A. thaliana* (*HIRD11*) and *M. truncatula* and the monocot *S. bicolor* (Fig. 6). The microsynteny of H-DHN genes of other angiosperms is likewise conserved (not shown). H-DHNs possess two exons, with the whole coding region contained within the first exon and the second exon being no-coding, while F- and Y-DHNs usually have two coding exons (not shown). The conserved exon-intron structure also points to a common origin of H-DHNs. In conclusion, the microsynteny of H-DHN genes is conserved in angiosperms, indicating their true orthologous status and common evolutionary origin.

**Figure 6.**
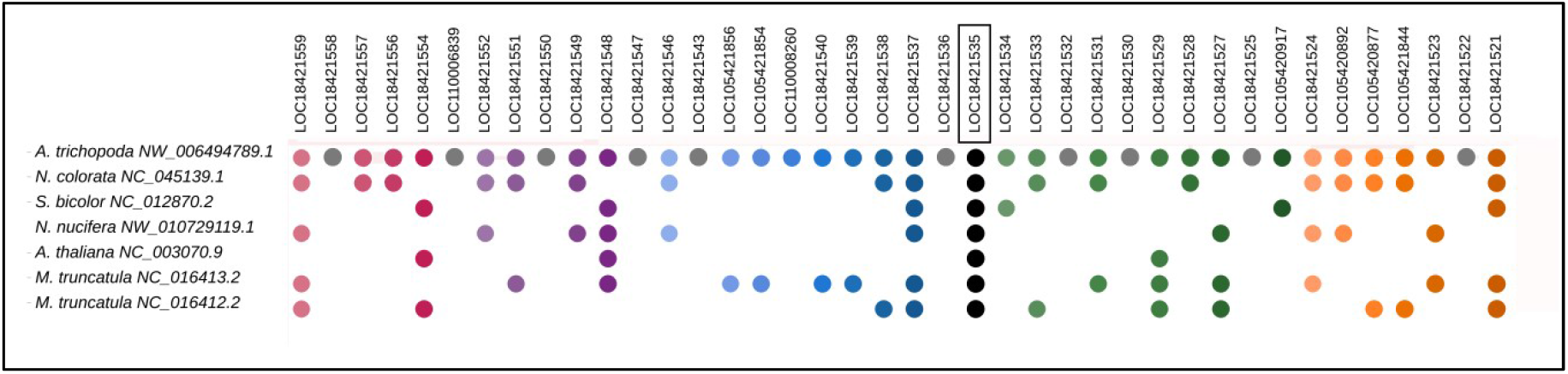
Microsynteny analysis of angiosperm H-DHNs genes. The genomic neighbourhood of the H-DHN gene of *A. trichopoda* (LOC18421535) is compared to that of other angiosperms. H-DHN genes are indicated as black dots, and a colour code indicates homologous genes present in the other species. Grey dots indicate genes only present in *A. trichopoda*. Some intervening genes in species other than *A. trichopoda* are not shown for clarity.

Along with one H-DHN gene, the genome of *A. trichopoda* contains two other DHN genes (LOC18424350 and LOC18426770). As mentioned above, in our phylogenetic tree, LOC18424350 is grouped together with F-DHNs and LOC18424350 with Y-DHNs (Fig. 5). Curiously, the F and Y motifs of these proteins are quite degenerated and are not readily recognised by the MEME program. A comparison of the genomic neighbourhoods of LOC18424350 and LOC18426770 of *A. trichopoda* with F- and Y-DHNs of *N. colorata* and *N. nucifera*, which likewise have only three DHN genes, reveals microsynteny conservation around these genes (Supplementary Figure S11), confirming that LOC18424350 and LOC18426770 belong to the F- and Y-DHN synthenic communities, respectively.

In summary, it is apparent that DHN genes of angiosperms can be generally divided into three syntenic communities, each one characterised, among other features, by the presence of the H, F or the Y motif. We propose that these orthologous groups be called F-dehydrins (community 1), Y-dehydrins (community 2) and H-dehydrins (community 3). The presence of only three dehydrin genes in the basal angiosperm *Amborella*, as well as in the early diverging Nymphaeales and the basal eudicot *N. nucifera*, suggests that the genomes of the first flowering plants had one H-, F- and Y-dehydrin gene each. Subsequent whole genome duplications in eudicots and monocots greatly increased the repertoire of these genes, specially those encoding F- and Y-dehydrins.

### Each DHN orthologous group presents distinctive hydrophylin biochemical properties

To analyse if the existence of three DHNs orthologous groups could result in proteins with distinctive characteristics, we compared the biochemical and biophysical properties of angiosperm DHNs from the H-, F-and Y-orthologous groups. Specifically, we determined general biochemical features such as molecular weight (MW) and isoelectric point (pI), as well as parameters related to the hydrophilin character of the proteins (Supplementary Table 1). We observed that each DHN orthologous group has a different MW distribution, with a characteristic statistical median (Fig.7A). H-DHNs are the smallest DHNs with the narrowest range of MW (10-16 kDa), reflecting that the number of residues and domain structure of the members of this DHN group are relatively constant. F-DHNs also have a compact MW distribution that ranges from 18 kDa to 35 kDa. Y-DHNs, on the other hand, present a main subgroup of proteins ranging from 10 kDa to 25 kDa and a number of DHNs with MW over 30 kDa that belong exclusively to monocot species. The high MW of this latter subgroup is not due to an increased number of conserved Y or K domains, but to the presence of long Gly-rich regions separating these domains (Supplementary Figure S4 to Figure S6). As for the isoelectric point, most H-DHNs present acidic pI values, with neutral and basic isoforms being found in some species (Fig 7B). F-DHNs have a very homogeneous acidic pI profile, with a unimodal distribution between 5 and 6. In contrast, Y-DHNs display a bimodal distribution consisting of two main subgroups of DHNs with basic and acid pI values, and a smaller subgroup with pI values close to neutrality. Interestingly, we were able to determine that almost all plant species have at least one basic and one acidic Y-DHN isoform, suggesting that a functional specialization of both types of proteins may have occurred during evolution. In monocots, the number of basic Y-DHNs is always greater than the acidic ones, and the opposite occurs in dicots. It should be noted that the early-diverging angiosperms *A. thrichopoda* and *N. colorata* encode a single basic Y-DHN in their genomes, which may represent the original pI character of these proteins in angiosperms (Supplementary Table 1).

**Figure 7.**
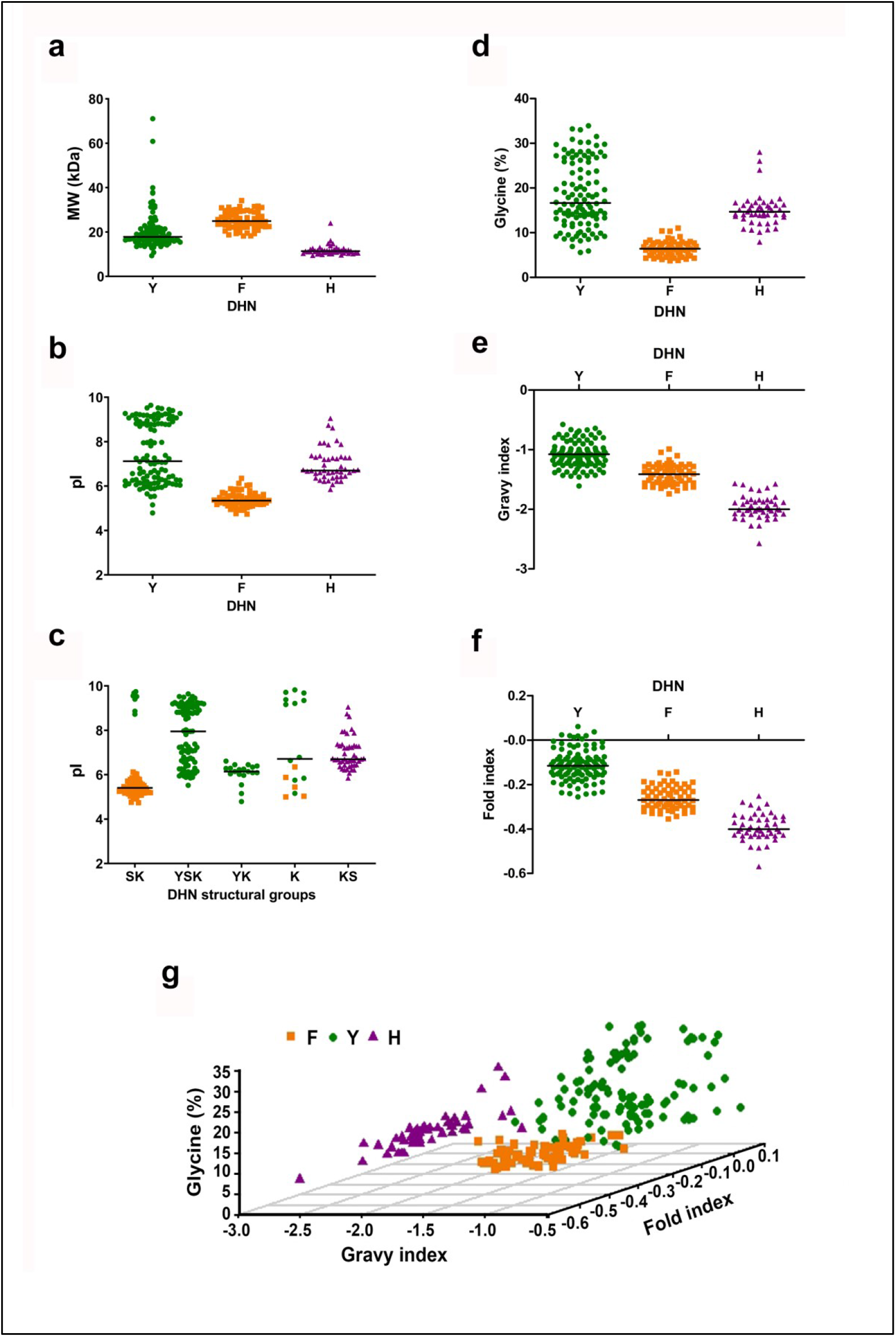
Distribution of biochemical and biophysical properties of angiosperm DHNs. Scatter plots show the distribution of molecular weight (**a**), isoelectric point (**b** and **c**), glycine content (**d**), GRAVY index (**e**), Fold Index (**f**) or glycine content, GRAVY and Fold index simultaneously (**g**) in orthologous or structural subgroups of DHNs. Members of the three orthologous groups of DHNs are colour-coded: Y-(green), F-(orange) and H-DHNs (violet).

When analysing the pI distribution in the five traditional DHN structural subgroups, it can be noted that the bimodal character observed in SK- and K-type DHNs strongly correlates with their evolutionary origin (Fig. 7C). For example, five of the K-type DHNs that display acidic pI, which corresponds to the pI of F-DHNs, belong to this orthologous group. Similarly, all members of the SK- and K-DHNs structural subgroups with high pI values belong to the Y-DHN orthologous group (Fig. 7B-7C). This suggests that DHN orthologous groups are better indicators of the pI character of DHNs compared to the traditional structural classification.

As for glycine content, both F- and H-DHNs present a compact and homogeneous distribution of percentage of glycine residues (Fig 7D). The DHNs with the lowest percentage of glycine (around 10%) are the F-DHNs. Remarkably, DHNs of the FSK3 structural subgroup are characterized by the presence of many proline stretches, which might play an equivalent role to that of glycine in terms of the disruption of the protein structure (Supplementary Figure S2 and Figure S3). Notably, a larger dispersion in the percentage of glycine residues is observed in Y-DHNs (range: 5% to 35%).

All DHNs display a negative GRAVY index, showing the characteristic hydrophilicity of this type of proteins (Fig. 7E). The scores for F-DHNs are distributed in the range of -1 to -1.7, overlaping with the other two groups. The Y-DHNs include the least hydrophilic proteins, with scores in the range of -0.5 to -1.5. H-DHNs, in contrast, are the most hydrophilic proteins, with GRAVY indexes ranging from -2.8 to -1.3. The atypical H-DHNs from the Malpighiales species *S. purpurea* and *P. trichocarpa* are the least hydrophilic dehydrins in this group, with a GRAVY index around -1.3. We also evaluated the Fold Index of DHNs using the FoldIndex algorithm, which estimates the mean net charge and hydrophobicity of a given protein sequence to predict if it is intrinsically unfolded^59^. The fold index of DHNs shows a similar distribution to that of the GRAVY index, with H-DHNs being the most intrinsically unfolded, while F- and Y-DHNs have a less unfolded character (Fig.7F).

In summary, in general terms, the biochemical and biophysical characteristics of DHNs correlate well with the three orthologous groups (Fig. 7G). Since these features are likely related to the function of DHNs, this suggests that functional studies of these proteins should take into consideration the phylogenetic framework proposed here.

## Discussion

Dehydrins are characterised by a great diversity of structural domains, arranged in various ways, which constitute the basis for the current classification into six structural subgroups, namely Kn-, SKn-, YnKn-, YnSKn-, KS-DHNs and the recently proposed FSKn-DHNs. However, the underlying evolutionary relationships between these DHNs in angiosperms and other plant groups have been unclear. In this work, we present a phylogenetic framework for DHNs that sheds light on the relationships between these proteins, specially in angiosperms. The main points of our work are: i) searches of DHN in plant genomic databases need to be done with a combination of HHM profiles to retrieve all types of DHN proteins; ii) KS-DHNs possess a new, conserved structural domain present at the N-terminus, which we named the H-domain; iii) phylogenetic and synteny analyses show that all angiosperm DHNs can be subdivided into three DHN orthologous groups, distinguished by the presence of the H-, F- or Y-domains, and iv) the psychochemical characteristics that are typical for DHNs correlate with each orthologous group, indicating that the evolutionary origin of DHNs should be taken into consideration when studying their function.

The reconstruction of the evolutionary history of DHNs is a complex task, due to the modular nature of these proteins, which are characterized by the presence of various small conserved segments surrounded by less conserved sequences of various lengths. Thus, the coupling of phylogenetic reconstruction with microsynteny analyses was crucial for the determination of the evolutionary relationships between DHNs. Angiosperm DHNs can be divided into three orthologous groups, H-DHNs, F-DHNs and Y-DHNs, which can, in most cases, be readly recognised by the presence of the H-, F- or Y-segments. All angiosperms analysed by us possess at least one DHN member of each homologous group, including the basal angiosperms *A. trichopoda* and *N. colorata*, indicating that the first angiosperms had genes enconding the three types of DHNs. Synteny analyses could not be extended to non-angiosperm species due to the fast rate of synteny loss that is typical for plants^50,60^.

Our analysis indicated that H-, F- and Y-DHNs are clearly distinguished from each other in features that characterize hydrophilins and intrinsically-disordered proteins (IDPs). Since all dehydrins of K and SK-structural subgroups actually belong to the F- and Y- syntenic groups, we were able to show that the classification based on the structural subgroups ends up putting in the same category DHNs with very different physicochemical properties. It has been observed that the cryoprotective capacity of DHNs depends on the size (hydrodynamic radius) and the intrisic disorder, highlighting the importance of the composition and size of the Phi segments, which are generally less conserved than the structural motifs^61^. It has been also demonstrated that it is the size and sequence composition of DHNs that is the most important for preventing aggregation, while for freeze damage it is the sequence composition that is most significant^62^. Thus, it seems that the simple presence of K and S segments would not be necessarily good predictors of the functional characteristics of DHNs.

It should be noted that the diversity of DHNs is not encompassed by the HMM model that is usually employed to search for DHN genes in the scientific literature, namely Pfam00257. Indeed, we show here that most H-DHNs (which belong to the KS-DHN structural subgroup) are not recognized by this model, which might be the reason that genomic-wide analyses of DHNs usually failed to retrieve many members of this orthologous group^18,20,22^. In view of this, we propose that studies aimed at identifying DHNs should use HMM profiles based on H-, F- and Y-DHNs separately in order to pinpoint all members of this protein family.

Importantly, we describe that KS-DHNs possess a new motif that we named the H-segment, due to the presence of two conserved His residues. This segment is always located at the N-terminus of the proteins and is predicted to have an α-helical structure. Thus, KS-DHNs can be better described as bearing a H-K-S organization of motifs. As mentioned above, phylogenetic and synteny analyses indicate that angiosperm DHNs are all evolutionarily related. The presence of DHNs with a distinct H-K-S organization in the lycophyte *S. moellendorffii* and the gymnosperm *Ginkgo biloba* strongly suggests that H-DHNs appeared in the early evolution of vascular plants. Although some KS-DHNs have been described before in the scientific literature, our work is the first, to our knowledge, to provide a thorough description of this group of DHNs.

The best studied member of the H-DHN group is the HIRD11 protein from *A. thaliana*. AtHIRD11 is expressed ubiquitously, with somewhat higher levels in flowers^54^. Functional studies showed that HIRD11 binds to metal ions and can protect proteins from heavy metal damage^54,63^ and can also reduce free radical generation^64^. Interestingly, both the binding to metals and the inhibition properties of HIRD11 depend on His residues, which are present in the H-segment. Importantly, Yokoyama et al (2020) have recently showed that both the K- and the H-segments (which the authors called K and NK1, respectively) of AtHIRD11 can protect proteins from freezing damage with similar efficiencies. Structurally, the presence of the K- and H-segments were needed for AtHIRD11 to transition from a disordered to an ordered state^65^. Overall, the functional results by Yokoyama et al (2020) show that the H-segment is an important component of H-DHNs, as implied by its high degree of phylogenetic conservation, and suggests that K- and H-segments might play overlapping roles in the activity of H-DHNs.

In conclusion, we consider that the classification of angiosperm DHNs into three homologous groups, as proposed here, better reflects the diversity of DHNs and should complement the traditional classification into six structural subgroups in the study of the function of these proteins.

## Acknowledgements

This work was supported by grant from the National Agency for the Promotion of Science and Technology of Argentina (ANPCyT, PICT 2015-3527) and National Scientific and Technical Research Council (CONICET, PIP 11220150100584). AMZ is a career research scientist from CONICET and AEM is a CONICET fellow.

## Author contributions

AMZ planned and designed the research. AEM and AMZ performed the research and analysed the data. AMZ wrote the article with contribution of AEM. Both authors read and approved the manuscript.

## Supplemental material

**Table S1:** List of all DHNs analysed in this study. It includes sequence name, taxonomic data, accession number, synteny/homologous group (H-, F- or Y-DHN), segmental structure and physicochemical characteristics.

**Fig. S1:** MEME analysis of unbiased DHN datbasey.

**Fig. S2:** Multiple sequence alignment of FSK2 dehydrins. Protein sequences of FSK2 from eudicot species were aligned with Clustal Omega and visualized with Jalview. Structural segments are indicated and a consensus sequence is shown below the alignment.

**Fig. S3:** Multiple sequences alignment of FSK3 dehydrins. Protein sequences of FSK3-DHNs from angiosperms were aligned with Clustal Omega and visualized with Jalview. Structural segments are indicated and a consensus sequence is shown below the alignment. Note that there is a lysine-rich region adjacent to the S-segment but it is not as conserved as the B-segment found in FSK2-DHNs (compare to Fig. S2).

**Fig. S4:** Multiple sequences alignment of YSKn dehydrins. Protein sequences of YSKn-DHNs from angiosperms were aligned with T-Coffee and visualized with Jalview. Structural segments are indicated and a consensus sequence is shown below the alignment.

**Fig. S5:** Multiple sequence alignment of dehydrins Y2SKn. Protein sequences of Y2SKn-DHNs from angiosperms were aligned with T-Coffee and visualized with Jalview. Structural segments are indicated and a consensus sequence is shown below the alignment. Y2SK2 and Y2SK3-DHNs are present in eudicots and the grass *Brachypodium distachyon*, while other Poaceae only have Y2SK2-DHNs.

**Fig. S6:** Multiple sequences alignment of Y3SKn dehydrins. Protein sequences of Y3SKn-DHNs from angiosperms were aligned with T-Coffee and visualized with Jalview. Structural segments are indicated and a consensus sequence is shown below the alignment.

**Fig. S7:** Multiple sequence alignment of HSK-dehydrins. Protein sequences of H-DHNs from vascular plants were aligned with Clustal Omega and visualized with Jalview. Structural segments are indicated and a consensus sequence is shown below the alignment. DHNs come from angiosperms except for proteins from *Selaginella moellendorffii* (Smo) and *Ginkgo biloba* (Gbi).

**Fig. S8:** Multiple sequence alignment of atypical H-DHNs from Malpighiales. (A) Alignment of HKS-DHNs from *P. trichocarpa* and *S. purpurea* and a HS-DHN from *P. trichocarpa*. (B) Atypical H-DHNs with multiple K segments interspersed with Phi-segments. Segments are indicated by a colour code: H (purple), K (red), S2 (blue) and Phi (green). Sequences were aligned with Clustal Omega and visualized with Jalview.

**Fig. S9:** Evolutionary relationships of bryophyte dehydrins. (A) Maximum-likelihood phylogenetic tree constructed with PhyML 3.0. Branches with bootstrap values over 90 are indicated with a circle. Note that DHN sequences from *P. patens* and *C. purpureus* form five homologous groups, while *S. fallax* DHNs are not grouped with the other sequences. (B) DHN sequences and homologous groups of *P. patens* and *C. purpureus*.

**Fig. S10:** Multiple sequence aligment of bryophyte DHNs. Segments are indicated by a colour code: K (red), Y (green) and S (blue). Note that Group I has a Y8K structure; the Y-segments with an asterisk (*) have a sequence identical to the Y-segments of angiosperms (DEYGNP), while the others have a modified Y-segment (DNYGN/QP). Group II has a KS-structure, Group III and V have a K2-structure and Group IV a K-structure. Sequences were aligned with T-Coffee and visualized with Jalview.

**Fig. S11:** Scatter plots of physicochemical features of angiosperm DHN-structural groups: Glycine content, GRAVY index and Fold index. Homologous groups are colour-coded: H-DHNs (purple), F-DHNs (orange) and Y-DHNs (green).

