## Supplemental Data for "Genomic diversification of dehydrin gene family in vascular plants: three distinctive orthologue groups and a novel KS-dehydrin conserved protein motif"

| Clade | Family | Specie | Synteny cluster | DHN structure | Protein accession | Lenght (aa) | MW | PI | Fold Index | GRAVY | Gly |
|---|---|---|---|---|---|---|---|---|---|---|---|
| Bryophytes | Ditrichaceae | Ceratodon purpureus | | | Cpu_CepurGG1.10G131500 | 661 | 69808 | 5,84 | -0,139 | -1,112 | 19,4 |
| Bryophytes | Ditrichaceae | Ceratodon purpureus | | | Cpu_CepurGG1.1G002400 | 139 | 14589 | 8,83 | -0,149 | -1,214 | 21,6 |
| Bryophytes | Ditrichaceae | Ceratodon purpureus | | | Cpu_CepurGG1.1G151900 | 223 | 23627 | 4,77 | -0,203 | -1,221 | 16,6 |
| Bryophytes | Ditrichaceae | Ceratodon purpureus | | | Cpu_CepurGG1.4G092400 | 195 | 20762 | 5,33 | -1,850 | -1,212 | 13,3 |
| Bryophytes | Ditrichaceae | Ceratodon purpureus | | | Cpu_CepurGG1.1G098000 | 291 | 30112 | 6,76 | -0,081 | 1,010 | 15,1 |
| Bryophytes | Ditrichaceae | Ceratodon purpureus | | | Cpu_CepurGG1.4G196600 | 242 | 26237 | 4,59 | -0,255 | -1,256 | 9,5 |
| Bryophytes | Marchantiaceae | Marchantia polymorpha | | | Mpo_Mapoly0053s0068 | 373 | 37921 | 6,55 | -0,096 | -1,04 | 22,0 |
| Bryophytes | Marchantiaceae | Marchantia polymorpha | | | Mpo_Mapoly0056s0073 | 505 | 52748 | 5,10 | -0,149 | -1,050 | 14,9 |
| Bryophytes | Marchantiaceae | Marchantia polymorpha | | | Mpo_Mapoly0114s0014 | 131 | 13811 | 8,11 | -0,038 | -0,877 | 7,6 |
| Bryophytes | Funariaceae | Physcomitrella patents | | | Ppa_PP00211G00460 | 554 | 59191 | 5,61 | -0,174 | -1,231 | 19,0 |
| Bryophytes | Funariaceae | Physcomitrella patents | | | Ppa_PP00052G01250 | 130 | 13772 | 8,03 | -0,142 | -1,215 | 16,9 |
| Bryophytes | Funariaceae | Physcomitrella patents | | | Ppa_PP00201G00250 | 184 | 19516 | 5,96 | -0,142 | -0,905 | 13,0 |
| Bryophytes | Funariaceae | Physcomitrella patents | | | Ppa_PP00421G00070 | 233 | 24905 | 4,86 | -0,210 | -1,225 | 16,7 |
| Bryophytes | Funariaceae | Physcomitrella patents | | | Ppa_PP00025G00800 | 238 | 24680 | 6,61 | -0,056 | -0,921 | 14,7 |
| Bryophytes | Funariaceae | Physcomitrella patents | | | Ppa_PP00442G00100 | 290 | 31371 | 4,90 | -0,215 | -1,24 | 10,3 |
| Bryophytes | Sphagnaceae | Sphagnum fallax | | | Sfa_Sphfalx0005s0334.1 | 357 | 37474 | 4,68 | -0,197 | -1,19 | 12,6 |
| Bryophytes | Sphagnaceae | Sphagnum fallax | | | Sfa_Sphfalx0010s0103.1 | 231 | 23736 | 4,17 | -0,215 | -1,197 | 16,5 |
| Bryophytes | Sphagnaceae | Sphagnum fallax | | | Sfa_Sphfalx0064s0013.1 | 220 | 23063 | 4,00 | -0,298 | -1,319 | 11,8 |
| Lycophytes | Selaginellaceae | Selaginella moellendorffii | H | HKS | Smo_437484 / XP_031475479 | 106 | 11527 | 7,88 | -0,252 | -1,565 | 12,3 |
| Lycophytes | Selaginellaceae | Selaginella moellendorffii | H | HS | Smo_446994 / XP_002987557 | 101 | 10661 | 9,70 | -0,194 | -1,215 | 7,9 |
| Gymnosperms | Ginkgoaceae | Ginkgo biloba | F | FSK2 | Gbi_GBI00024041 | 181 | 19874 | 6,74 | -0,243 | -1,513 | 11,6 |
| Gymnosperms | Ginkgoaceae | Ginkgo biloba | F | FK2 | Gbi_GBI00029052 | 163 | 17666 | 6,61 | -0,284 | -1,680 | 16,6 |
| Gymnosperms | Ginkgoaceae | Ginkgo biloba | H | HKS | Gbi_GBI00018450 | 89 | 10153 | 9,18 | -0,449 | -2,088 | 10,1 |
| Gymnosperms | Pinaceae | Picea abies | F | FK | Pab_MA_120345g0010 | 171 | 18498 | 6,72 | -0,286 | -1,686 | 18,1 |
| Gymnosperms | Pinaceae | Picea abies | F | FSK3 | Pab_MA_130855g0010 | 209 | 23137 | 5,98 | -0,331 | -1,726 | 12,9 |
| Gymnosperms | Pinaceae | Picea abies | F | FK | Pab_MA_151075g0010 | 171 | 18493 | 6,76 | -0,259 | -1,6 | 17,0 |
| Gymnosperms | Pinaceae | Picea abies | F | FSK | Pab_MA_187114g0010 | 154 | 16675 | 6,25 | -0,202 | -1,371 | 12,3 |
| Gymnosperms | Pinaceae | Picea abies | F | F | Pab_MA_19049g0010 | 75 | 8303 | 4,97 | -0,201 | -1,085 | 9,3 |
| Gymnosperms | Pinaceae | Picea abies | F | FK | Pab_MA_205576g0010 | 138 | 15498 | 8,33 | -0,223 | -1,433 | 9,4 |
| Gymnosperms | Pinaceae | Picea abies | F | F | Pab_MA_448618g0010 | 69 | 7665 | 5,50 | -0,274 | -1,386 | 11,6 |
| Gymnosperms | Pinaceae | Picea abies | F | FSK3 | Pab_MA_474985g0010 | 201 | 22043 | 6,51 | -0,287 | -1,626 | 13,4 |
| Gymnosperms | Pinaceae | Picea abies | F | FK2 | Pab_MA_95995g0010 | 84 | 9222 | 9,23 | -0,287 | -1,554 | 10,7 |
| Gymnosperms | Pinaceae | Picea abies | | | Pab_MA_104214g0010 | 155 | 16696 | 6,59 | -0,207 | -1,408 | 12,9 |
| Gymnosperms | Pinaceae | Picea abies | | | Pab_MA_10430236g0010 | 128 | 13641 | 9,19 | -0,110 | -1,06 | 9,4 |
| Gymnosperms | Pinaceae | Picea abies | | | Pab_MA_107783g0020 | 182 | 19573 | 9,79 | -0,139 | -1,052 | 9,3 |
| Gymnosperms | Pinaceae | Picea abies | | | Pab_MA_144878g0010 | 86 | 9103 | 9,90 | -0,146 | -1,063 | 9,3 |
| Gymnosperms | Pinaceae | Picea abies | | | Pab_MA_17462g0010 | 86 | 9075 | 9,90 | -0,161 | -1,114 | 11,6 |
| Gymnosperms | Pinaceae | Picea abies | | | Pab_MA_2408574g0010 | 90 | 9575 | 7,11 | -0,249 | -1,584 | 16,7 |
| Gymnosperms | Pinaceae | Picea abies | | | Pab_MA_43017g0010 | 148 | 15876 | 9,87 | -0,135 | -1,063 | 8,8 |
| Gymnosperms | Pinaceae | Picea abies | | | Pab_MA_45025g0010 | 188 | 20137 | 9,87 | -0,144 | -1,074 | 9,0 |
| Gymnosperms | Pinaceae | Picea abies | | | Pab_MA_4978040g0010 | 164 | 17614 | 9,78 | 0,133 | -1,074 | 9,1 |
| Gymnosperms | Pinaceae | Picea abies | | | Pab_MA_59157g0010 | 53 | 6145 | 6,53 | -0,118 | -1,162 | 7,5 |
| Gymnosperms | Pinaceae | Picea abies | | | Pab_MA_65580g0010 | 250 | 27478 | 6,86 | -0,274 | -1,627 | 12,8 |
| Gymnosperms | Pinaceae | Picea abies | | | Pab_MA_816406g0020 | 77 | 8602 | 5,75 | -0,183 | -1,330 | 9,1 |
| Angiosperms-Basal | Amborellaceae | Amborella trichopoda | F* | SK3 | Atr_evm_27.TU.AmTr_v1.0_scaffold00001.295 | 212 | 23366 | 5,79 | -0,258 | -1,446 | 10,4 |
| Angiosperms-Basal | Amborellaceae | Amborella trichopoda | H | HKS | Atr_evm_27.model.AmTr_v1.0_scaffold00004.97 / XP_006826344.1 | 105 | 11644 | 7,30 | -0,343 | -1,888 | 17,1 |
| Angiosperms-Basal | Amborellaceae | Amborella trichopoda | Y* | SK2 | Atr_evm_27.TU.AmTr_v1.0_scaffold00082.27 | 148 | 15668 | 8,72 | -0,169 | -1,282 | 21,6 |
| Angiosperms-Basal | Nymphaeaceae | Nymphea colorata | F | FSK3 | Nco_Nycol.J01522 | 218 | 24307 | 5,2 | -0,214 | -1,274 | 8,7 |
| Angiosperms-Basal | Nymphaeaceae | Nymphea colorata | F | SK3 | Nco_Nycol.I01776 | 243 | 26810 | 5,71 | -0,221 | -1,232 | 7,8 |
| Angiosperms-Basal | Nymphaeaceae | Nymphea colorata | H | HKS | Nco_Nycol.B01145/ XP_031475479 | 94 | 10972 | 6,96 | -0,551 | -2,528 | 11,7 |
| Angiosperms-Basal | Nymphaeaceae | Nymphea colorata | Y | YSK2 | Nco_Nycol.B00068 | 251 | 26583 | 9,51 | -0,056 | -0,794 | 16,7 |
| Angiosperms-Monocots | Musaceae | Musa acuminata | F | FSK3 | Macu_GSMUA_Achr11G16760_001 | 220 | 24866 | 5,08 | -0,279 | -1,419 | 7,3 |
| Angiosperms-Monocots | Musaceae | Musa acuminata | H | HKS | Macu_GSMUA_Achr4T21460_001 | 92 | 10561 | 7,28 | -0,391 | -2,045 | 12,0 |
| Angiosperms-Monocots | Musaceae | Musa acuminata | Y* | SK2 | Macu_GSMUA_Achr4G11310_001 | 96 | 10195 | 9,74 | -0,254 | -1,367 | 19,8 |
| Angiosperms-Monocots | Bromeliaceae | Anana comusus | F | FSK3 | Aco_Aco011968 | 262 | 28992 | 4,92 | -0,233 | -1,287 | 10,3 |
| Angiosperms-Monocots | Bromeliaceae | Anana comusus | H | HS | Aco_OAY73247.1 | 91 | 10255 | 8,66 | -0,410 | -2,034 | 13,2 |
| Angiosperms-Monocots | Bromeliaceae | Anana comusus | Y | YSK2YSK2 | Aco_Aco016515 | 341 | 34665 | 9,44 | -0,860 | -0,86 | 22,6 |
| Angiosperms-Monocots | Bromeliaceae | Anana comusus | Y | YSK2 | Aco_Aco016518 | 89 | 9311 | 9,52 | -1,374 | -1,374 | 21,3 |
| Angiosperms-Monocots | Bromeliaceae | Anana comusus | Y* | SK2 | Aco_Aco016516 | 163 | 16700 | 9,55 | -0,229 | -1,118 | 20,9 |
| Angiosperms-Monocots | Bromeliaceae | Anana comusus | Y* | K2 | Aco_Aco021122 | 221 | 23375 | 10,02 | 0,002 | -0,57 | 15,4 |
| Angiosperms-Monocots | Bromeliaceae | Anana comusus | Y* | SK | Aco_Aco021124 | 129 | 13717 | 9,95 | -0,169 | -1,077 | 10,1 |
| Angiosperms-Monocots | Zosteraceae | Zostera marina | F | FSK3 | Zmar_Zosma103g00290 | 198 | 22009 | 5,37 | -0,223 | -1,338 | 7,1 |

| | | | | | | | | | | | |
|---|---|---|---|---|---|---|---|---|---|---|---|
| Angiosperms-Monocots | Zosteraceae | Zostera marina | H | HKS | Zmar_Zosma440g00040 | 92 | 10333 | 5,85 | -0,451 | -2,027 | 14,1 |
| Angiosperms-Monocots | Poaceae | Brachypodium distachyon | F | FSK3 | Bdi_Bradi3g51200 | 254 | 27504 | 5,24 | -0,144 | -1,043 | 7,9 |
| Angiosperms-Monocots | Poaceae | Brachypodium distachyon | F | FSK3 | Bdi_Bradi5g10860 | 252 | 27506 | 5,24 | -0,185 | -1,172 | 8,3 |
| Angiosperms-Monocots | Poaceae | Brachypodium distachyon | H | HKS | Bdi_Bradi1g13330 | 100 | 11313 | 7,23 | -0,424 | -2,152 | 14,0 |
| Angiosperms-Monocots | Poaceae | Brachypodium distachyon | Y | YSK2 | Bdi_Bradi1g37410 | 163 | 16348 | 9,13 | -0,107 | -1,068 | 25,8 |
| Angiosperms-Monocots | Poaceae | Brachypodium distachyon | Y | Y2SK2 | Bdi_Bradi2g47575 | 557 | 60848 | 6,03 | 0,018 | -0,665 | 9,9 |
| Angiosperms-Monocots | Poaceae | Brachypodium distachyon | Y | YSK2 | Bdi_Bradi3g43855 | 157 | 16125 | 9,07 | -0,133 | -1,148 | 25,5 |
| Angiosperms-Monocots | Poaceae | Brachypodium distachyon | Y | YSK2 | Bdi_Bradi3g43870 | 183 | 18456 | 9,25 | -0,121 | -1,101 | 26,8 |
| Angiosperms-Monocots | Poaceae | Brachypodium distachyon | Y | Y2SK3 | Bdi_Bradi4g19525 | 395 | 37817 | 9,03 | 0,024 | -0,669 | 28,1 |
| Angiosperms-Monocots | Poaceae | Brachypodium distachyon | Y* | SK2 | Bdi_Bradi4g22280 | 107 | 10993 | 9,40 | -0,165 | -1,194 | 19,6 |
| Angiosperms-Monocots | Poaceae | Brachypodium distachyon | Y* | SK2 | Bdi_Bradi4g22290 | 143 | 14469 | 8,86 | -0,096 | -1,045 | 22,4 |
| Angiosperms-Monocots | Poaceae | Oryza sativa | F | FSK3 | Osa_LOC_Os02g44870 | 290 | 30922 | 5,68 | -0,147 | -1,101 | 7,9 |
| Angiosperms-Monocots | Poaceae | Oryza sativa | H | HKS | Osa_LOC_Os03g45280 | 92 | 10363 | 6,70 | -0,404 | -2,015 | 15,2 |
| Angiosperms-Monocots | Poaceae | Oryza sativa | Y | Y2SK2 | Osa_LOC_Os01g50700 | 652 | 71043 | 7,07 | 0,061 | -0,577 | 9,7 |
| Angiosperms-Monocots | Poaceae | Oryza sativa | Y | YSK2 | Osa_LOC_Os11g26570 | 326 | 31251 | 8,95 | 0,020 | -0,687 | 26,7 |
| Angiosperms-Monocots | Poaceae | Oryza sativa | Y | YSK2 | Osa_LOC_Os11g26750 | 151 | 15551 | 9,13 | -0,165 | -1,248 | 18,5 |
| Angiosperms-Monocots | Poaceae | Oryza sativa | Y | YSK2 | Osa_LOC_Os11g26760 | 164 | 16724 | 9,25 | -0,108 | -1,049 | 26,2 |
| Angiosperms-Monocots | Poaceae | Oryza sativa | Y | YSK2 | Osa_LOC_Os11g26780 | 164 | 16537 | 9,27 | -0,102 | -1,033 | 26,8 |
| Angiosperms-Monocots | Poaceae | Oryza sativa | Y | YSK2 | Osa_LOC_Os11g26790 | 172 | 17325 | 9,19 | -0,109 | -1,076 | 27,3 |
| Angiosperms-Monocots | Poaceae | Sorghum bicolor | F | FSK3 | Sbi_Sobic.004G286600 | 283 | 31029 | 5,79 | -0,219 | -1,282 | 7,1 |
| Angiosperms-Monocots | Poaceae | Sorghum bicolor | H | HKS | Sbi_Sobic.001G149500 / XP_002466730.1 | 101 | 11606 | 6,73 | -0,449 | -2,168 | 10,9 |
| Angiosperms-Monocots | Poaceae | Sorghum bicolor | Y | Y2SK2 | Sbi_Sobic.003G270200 | 208 | 21903 | 7,95 | -0,075 | -1,006 | 13,9 |
| Angiosperms-Monocots | Poaceae | Sorghum bicolor | Y | YSK2 | Sbi_Sobic.009G116700 | 152 | 15399 | 8,81 | -0,122 | -1,132 | 27,6 |
| Angiosperms-Monocots | Poaceae | Sorghum bicolor | Y | YSK3 | Sbi_Sobic.010G041900 | 388 | 37488 | 8,50 | -0,022 | -0,836 | 30,9 |
| Angiosperms-Monocots | Poaceae | Zea mays | F | FSK3 | Zma_GRMZM2G147014 | 290 | 31440 | 6,05 | -0,187 | -1,250 | 7,6 |
| Angiosperms-Monocots | Poaceae | Zea mays | F | FSK3 | Zma_GRMZM2G373522 | 289 | 31466 | 5,51 | -0,233 | -1,300 | 6,9 |
| Angiosperms-Monocots | Poaceae | Zea mays | H | HKS | Zma_GRMZM2G169372 | 100 | 11312 | 6,59 | -0,431 | -2,075 | 14,0 |
| Angiosperms-Monocots | Poaceae | Zea mays | H | HKS | Zma_GRMZM2G448511 | 108 | 12199 | 6,22 | -0,482 | -2,158 | 13,9 |
| Angiosperms-Monocots | Poaceae | Zea mays | Y | YSK3 | Zma_GRMZM2G052364 | 325 | 31828 | 8,56 | 0,005 | -0,745 | 28,0 |
| Angiosperms-Monocots | Poaceae | Zea mays | Y | YSK2 | Zma_GRMZM2G079440 | 168 | 17075 | 8,78 | -0,124 | -1,144 | 27,4 |
| Angiosperms-Monocots | Poaceae | Zea mays | Y | YSK3 | Zma_GRMZM2G098750 | 326 | 31690 | 7,37 | -0,005 | -0,798 | 28,2 |
| Angiosperms-Monocots | Poaceae | Panicum hallii | F | FSK3 | Pha_Pahal.A02866 | 284 | 31103 | 5,86 | -0,193 | -1,234 | 7,7 |
| Angiosperms-Monocots | Poaceae | Panicum hallii | H | HKS | Pha_Pahal.I02369 | 104 | 11884 | 6,76 | -0,428 | -2,102 | 10,6 |
| Angiosperms-Monocots | Poaceae | Panicum hallii | Y | Y2SK2 | Pha_Pahal.E01980 | 216 | 22036 | 6,20 | -0,015 | -0,769 | 16,2 |
| Angiosperms-Monocots | Poaceae | Panicum hallii | Y | YSK2 | Pha_Pahal.J00522 | 166 | 16726 | 8,78 | -0,107 | -1,087 | 29,5 |
| Angiosperms-Monocots | Poaceae | Panicum hallii | Y | YSK3 | Pha_Pahal.J00526 | 343 | 33255 | 8,02 | -0,001 | -0,804 | 28,6 |
| Angiosperms-Monocots | Poaceae | Setaria italica | F | FSK3 | Sit_Seita.1G267200 | 290 | 31644 | 5,76 | -0,199 | -1,246 | 7,6 |
| Angiosperms-Monocots | Poaceae | Setaria italica | H | HKS | Sit_Seita.9G151800 | 102 | 11640 | 7,32 | -0,400 | -2,075 | 10,8 |
| Angiosperms-Monocots | Poaceae | Setaria italica | Y | YSK2 | Sit_Seita.5G166200 | 138 | 14126 | 9,19 | -0,145 | -1,178 | 21,0 |
| Angiosperms-Monocots | Poaceae | Setaria italica | Y | YSK2 | Sit_Seita.5G166300 | 138 | 14101 | 8,90 | -0,142 | -1,192 | 21,0 |
| Angiosperms-Monocots | Poaceae | Setaria italica | Y | Y2SK2 | Sit_Seita.5G286000 | 223 | 22836 | 6,41 | -0,033 | -0,843 | 17,9 |
| Angiosperms-Monocots | Poaceae | Setaria italica | Y | YSK3 | Sit_Seita.8G115200 | 347 | 33741 | 8,99 | -0,019 | -0,804 | 27,7 |
| Angiosperms-Monocots | Poaceae | Setaria italica | Y | YSK2 | Sit_Seita.8G115400 | 169 | 16914 | 8,81 | -0,095 | -1,050 | 30,2 |
| Angiosperms-Monocots | Poaceae | Setaria italica | Y* | SK2 | Sit_Seita.8G115100 | 192 | 19613 | 9,68 | -0,110 | -0,985 | 20,3 |
| Angiosperms-Early Eudicot | Nelumbonaceae | Nelumbo nucifera | F | FSK3 | Nnu_XP_010266960 | 236 | 27015 | 5,07 | -0,355 | -1,614 | 6,4 |
| Angiosperms-Early Eudicot | Nelumbonaceae | Nelumbo nucifera | H | HKS | Nnu_XP_019054435 | 100 | 11560 | 7,37 | -0,431 | -2,172 | 12,0 |
| Angiosperms-Early Eudicot | Nelumbonaceae | Nelumbo nucifera | Y | YSK2 | Nnu_XP_010254713 | 147 | 14855 | 9,52 | -0,175 | -1,235 | 27,2 |
| Angiosperms-Eudicots | Ranunculaceae | Aquilegia coerulea | F | FSK4 | Aqcoe5G125000 | 247 | 28012 | 5,19 | -0,303 | -1,551 | 6,9 |
| Angiosperms-Eudicots | Ranunculaceae | Aquilegia coerulea | F | FSK3 | Aqcoe6G197600 | 233 | 26392 | 5,17 | -0,234 | -1,285 | 4,3 |
| Angiosperms-Eudicots | Ranunculaceae | Aquilegia coerulea | F* | K2 | Aqcoe5G125200 | 213 | 23796 | 5,03 | -0,331 | -1,247 | 5,6 |
| Angiosperms-Eudicots | Ranunculaceae | Aquilegia coerulea | H | HKS | Aqcoe2G065100 | 93 | 10070 | 8,74 | -0,287 | -1,639 | 16,1 |
| Angiosperms-Eudicots | Ranunculaceae | Aquilegia coerulea | Y | YK3 | Aqcoe5G186200 | 152 | 16357 | 6,23 | -0,206 | -1,384 | 18,4 |
| Angiosperms-Eudicots | Ranunculaceae | Aquilegia coerulea | Y | Y2SK2 | Aqcoe6G128700 | 202 | 19775 | 8,76 | -0,075 | -0,99 | 29,7 |
| Angiosperms-Eudicots | Apiaceae | Daucus carota | F | FSK3 | Dca_DCAR_009522 | 213 | 23796 | 5,03 | -0,215 | -1,247 | 5,6 |
| Angiosperms-Eudicots | Apiaceae | Daucus carota | H | HKS | Dca_XP_017246606.1 | 90 | 9849 | 8,05 | -0,427 | -1,913 | 17,8 |
| Angiosperms-Eudicots | Apiaceae | Daucus carota | H | HKS | Dca_XP_017249516.1 | 103 | 11575 | 7,86 | -0,362 | -2,130 | 10,7 |
| Angiosperms-Eudicots | Apiaceae | Daucus carota | Y | Y2SK3 | Dca_DCAR_017310 | 199 | 20779 | 5,85 | -0,187 | -1,270 | 19,6 |
| Angiosperms-Eudicots | Apiaceae | Daucus carota | Y | Y2SK2 | Dca_DCAR_017311 | 158 | 16209 | 6,70 | -0,109 | -1,113 | 23,4 |
| Angiosperms-Eudicots | Apiaceae | Daucus carota | Y | Y3SK2 | Dca_DCAR_028967 | 261 | 25524 | 8,77 | 0,012 | -0,717 | 23,4 |
| Angiosperms-Eudicots | Apiaceae | Daucus carota | Y* | SK2 | Dca_DCAR_021479 | 117 | 11958 | 9,52 | -0,185 | -1,241 | 26,5 |
| Angiosperms-Eudicots | Apiaceae | Daucus carota | Y* | K | Dca_DCAR_024077 | 106 | 10891 | 5,84 | -0,026 | -0,711 | 17,0 |
| Angiosperms-Eudicots | Phrymaceae | Erythranthe guttata | F | FSK2 | Mgu_Migut.C00923 | 264 | 29646 | 4,91 | -0,152 | -0,989 | 4,9 |
| Angiosperms-Eudicots | Phrymaceae | Erythranthe guttata | H | HKS | Mgu_Migut.J00768 | 119 | 12995 | 6,21 | -0,346 | -1,789 | 17,6 |

| | | | | | | | | | | | |
|---|---|---|---|---|---|---|---|---|---|---|---|
| Angiosperms-Eudicots | Phrymaceae | Erythranthe guttata | H | HKS | Mgu_Migut.L01868 | 107 | 12109 | 6,36 | -0,433 | -1,997 | 14,0 |
| Angiosperms-Eudicots | Phrymaceae | Erythranthe guttata | Y | Y3SK2 | Mgu_Migut.B00209 | 203 | 21376 | 7,00 | -0,081 | -1,025 | 13,8 |
| Angiosperms-Eudicots | Phrymaceae | Erythranthe guttata | Y | Y2SK3 | Mgu_Migut.H02513 | 235 | 23959 | 6,79 | -0,111 | -1,113 | 26,8 |
| Angiosperms-Eudicots | Solanaceae | Solanum tuberosum | F | FSK3 | Stu_PGSC0003DMG400009968 | 209 | 23673 | 5,24 | -0,289 | -1,499 | 6,2 |
| Angiosperms-Eudicots | Solanaceae | Solanum tuberosum | H | HS | Stu_XP_006355701.1 | 91 | 10475 | 7,23 | -0,411 | -2,109 | 12,1 |
| Angiosperms-Eudicots | Solanaceae | Solanum tuberosum | H | HKS | Stu_XP_006364877.1 | 89 | 10207 | 8,62 | -0,485 | -2,276 | 10,1 |
| Angiosperms-Eudicots | Solanaceae | Solanum tuberosum | H | HS | Stu_XP_006365372.1 | 112 | 13497 | 9,54 | -0,739 | -2,852 | 5,4 |
| Angiosperms-Eudicots | Solanaceae | Solanum tuberosum | Y | YSK2 | Stu_PGSC0003DMG400003530 | 140 | 14534 | 7,07 | -0,151 | -1,268 | 27,1 |
| Angiosperms-Eudicots | Solanaceae | Solanum tuberosum | Y | Y2SK2 | Stu_PGSC0003DMG400003531 | 157 | 16659 | 7,23 | -0,134 | -1,214 | 19,7 |
| Angiosperms-Eudicots | Solanaceae | Solanum tuberosum | Y | Y3SK2 | Stu_PGSC0003DMG400030949 | 243 | 25121 | 7,38 | 0,020 | -0,715 | 15,6 |
| Angiosperms-Eudicots | Solanaceae | Solanum tuberosum | Y | YSK2 | Stu_PGSC0003DMG400034095 | 140 | 14938 | 6,59 | -0,101 | -1,085 | 14,3 |
| Angiosperms-Eudicots | Solanaceae | Solanum lycopersicum | F | FSK3 | Solyc04g082200 | 206 | 23111 | 5,12 | -0,292 | -1,488 | 6,8 |
| Angiosperms-Eudicots | Solanaceae | Solanum lycopersicum | H | HKS | Solyc_Solyc05g053070 | 89 | 10181 | 7,20 | -0,401 | -2,076 | 12,4 |
| Angiosperms-Eudicots | Solanaceae | Solanum lycopersicum | Y | Y3SK2 | Solyc01g109920 | 249 | 25363 | 7,09 | -0,022 | -0,838 | 17,3 |
| Angiosperms-Eudicots | Solanaceae | Solanum lycopersicum | Y | YSK2 | Solyc02g062390 | 133 | 14432 | 9,02 | -0,086 | -0,984 | 12,0 |
| Angiosperms-Eudicots | Solanaceae | Solanum lycopersicum | Y | Y2SK2 | Solyc02g084840 | 157 | 16660 | 7,23 | -0,149 | -1,261 | 19,7 |
| Angiosperms-Eudicots | Solanaceae | Solanum lycopersicum | Y | YSK2 | Solyc02g084850 | 130 | 13948 | 6,06 | -0,205 | -1,368 | 23,8 |
| Angiosperms-Eudicots | Solanaceae | Solanum lycopersicum | Y* | K2 | Solyc01g065820 | 78 | 8871 | 9,82 | -0,354 | -1,636 | 12,8 |
| Angiosperms-Eudicots | Solanaceae | Solanum lycopersicum | Y* | K2 | Solyc02g093255 | 174 | 19503 | 9,71 | -0,023 | -0,649 | 9,2 |
| Angiosperms-Eudicots | Solanaceae | Solanum lycopersicum | Y* | K2 | Solyc09g074765 | 102 | 11735 | 9,68 | -0,302 | -1,504 | 10,8 |
| Angiosperms-Eudicots | Crassulaceae | Kalanchoe fedtschenkoi | F | FSK2 | Kfe_Kaladp0040s0609 | 192 | 21369 | 4,93 | -0,247 | -1,292 | 6,8 |
| Angiosperms-Eudicots | Crassulaceae | Kalanchoe fedtschenkoi | F | FSK3 | Kfe_Kaladp0477s0002 | 195 | 21615 | 5,28 | -0,190 | -1,262 | 6,7 |
| Angiosperms-Eudicots | Crassulaceae | Kalanchoe fedtschenkoi | H | HKS | Kfe_Kaladp0098s0091 | 87 | 9643 | 7,20 | -0,396 | -1,834 | 13,8 |
| Angiosperms-Eudicots | Crassulaceae | Kalanchoe fedtschenkoi | H | HKS | Kfe_Kaladp0911s0009 | 107 | 11903 | 6,42 | -0,326 | -1,938 | 13,1 |
| Angiosperms-Eudicots | Crassulaceae | Kalanchoe fedtschenkoi | Y | Y3SK2 | Kfe_Kaladp0018s0236 | 153 | 15858 | 6,63 | -0,097 | -1,052 | 16,3 |
| Angiosperms-Eudicots | Crassulaceae | Kalanchoe fedtschenkoi | Y | Y2SK2 | Kfe_Kaladp0040s0543 | 142 | 15387 | 7,97 | -0,130 | -1,179 | 19,0 |
| Angiosperms-Eudicots | Crassulaceae | Kalanchoe fedtschenkoi | Y | Y3SK | Kfe_Kaladp0047s0007 | 178 | 17922 | 6,75 | 0,037 | -0,642 | 17,4 |
| Angiosperms-Eudicots | Myrtaceae | Eucalyptus grandis | F | FK2 | Egr_Eucgr.F01726 | 210 | 23781 | 5,00 | -0,212 | -1,220 | 6,2 |
| Angiosperms-Eudicots | Myrtaceae | Eucalyptus grandis | H | HKS | Egr_XP_010036245.1 | 104 | 11676 | 6,45 | -0,417 | -2,004 | 16,3 |
| Angiosperms-Eudicots | Myrtaceae | Eucalyptus grandis | Y | Y3S | Egr_Eucgr.I01292 | 101 | 10782 | 5,65 | -0,117 | -1,030 | 14,9 |
| Angiosperms-Eudicots | Myrtaceae | Eucalyptus grandis | Y | Y2SK2 | Egr_Eucgr.I02392 | 170 | 17647 | 7,81 | -0,155 | -1,264 | 24,1 |
| Angiosperms-Eudicots | Myrtaceae | Eucalyptus grandis | Y | YSK2 | Egr_Eucgr.I02395 | 209 | 20521 | 5,91 | -0,088 | -1,034 | 33,0 |
| Angiosperms-Eudicots | Myrtaceae | Eucalyptus grandis | Y | Y2SK2 | Egr_Eucgr.J02380 | 137 | 14774 | 8,81 | -0,184 | -1,329 | 14,6 |
| Angiosperms-Eudicots | Vitaceae | Vitis vinifera | F | FSK2 | Vvi_VIT_218s0001g00360 | 206 | 23483 | 5,18 | -0,305 | -1,514 | 5,3 |
| Angiosperms-Eudicots | Vitaceae | Vitis vinifera | H | HS | Vvi_NP_001268149.1 | 126 | 13554 | 6,59 | -0,331 | -1,775 | 27,8 |
| Angiosperms-Eudicots | Vitaceae | Vitis vinifera | Y | Y3SK2 | Vvi_VIT_203s0038g04390 | 191 | 20064 | 6,26 | -0,114 | -1,030 | 12,6 |
| Angiosperms-Eudicots | Vitaceae | Vitis vinifera | Y | YK2 | Vvi_VIT_204s0023g02480 | 130 | 13917 | 9,27 | -0,241 | -1,459 | 18,5 |
| Angiosperms-Eudicots | Malpighiales | Ricinus communis | F | FSK3 | Rco_30170.t000735 | 230 | 26018 | 5,25 | -0,330 | -1,565 | 4,8 |
| Angiosperms-Eudicots | Malpighiales | Ricinus communis | H | HKS | Rco_30072.m000963 | 96 | 10915 | 6,64 | -0,409 | -2,000 | 13,5 |
| Angiosperms-Eudicots | Malpighiales | Ricinus communis | Y | Y2SK2 | Rco_29634.t000016 | 149 | 16722 | 7,25 | -0,256 | -1,608 | 14,1 |
| Angiosperms-Eudicots | Malpighiales | Ricinus communis | Y | YSK2 | Rco_29634.t000017 | 146 | 16028 | 8,74 | -0,200 | -1,384 | 16,4 |
| Angiosperms-Eudicots | Malpighiales | Ricinus communis | Y | Y3SK2 | Rco_30131.t000233 | 189 | 20330 | 5,88 | -0,155 | -1,161 | 9,5 |
| Angiosperms-Eudicots | Malpighiales | Linum usitatissimum | F | FSK2 | Lus_Lus10005652.g | 201 | 22112 | 5,85 | -0,198 | -1,309 | 6,5 |
| Angiosperms-Eudicots | Malpighiales | Linum usitatissimum | F | FSK2 | Lus_Lus10021240.g | 215 | 23562 | 6,03 | -0,204 | -1,351 | 8,8 |
| Angiosperms-Eudicots | Malpighiales | Linum usitatissimum | F | FSK2 | Lus_Lus10021827.g | 225 | 24978 | 5,21 | -0,257 | -1,381 | 6,2 |
| Angiosperms-Eudicots | Malpighiales | Linum usitatissimum | F | FSK2 | Lus_Lus10034568.g | 229 | 25306 | 5,48 | -0,238 | -1,365 | 7,4 |
| Angiosperms-Eudicots | Malpighiales | Linum usitatissimum | F* | SK3 | Lus_Lus10022643.g | 179 | 19861 | 5,47 | -0,188 | -1,226 | 11,2 |
| Angiosperms-Eudicots | Malpighiales | Linum usitatissimum | H | HKS | Lus_Lus10020271.g | 93 | 10422 | 6,61 | -0,412 | -2,009 | 15,1 |
| Angiosperms-Eudicots | Malpighiales | Linum usitatissimum | Y | YSK2 | Lus_Lus10014280.g | 154 | 16240 | 9,07 | -0,152 | -1,207 | 14,9 |
| Angiosperms-Eudicots | Malpighiales | Linum usitatissimum | Y | Y3SK2 | Lus_Lus10017977.g | 218 | 23017 | 5,94 | -0,038 | -0,750 | 9,2 |
| Angiosperms-Eudicots | Malpighiales | Linum usitatissimum | Y | YSK2 | Lus_Lus10025983.g | 146 | 15485 | 9,05 | -0,157 | -1,222 | 15,1 |
| Angiosperms-Eudicots | Malpighiales | Linum usitatissimum | Y | Y3SK2 | Lus_Lus10041969.g | 218 | 23017 | 5,94 | -0,023 | -0,750 | 9,2 |
| Angiosperms-Eudicots | Malpighiales | Manihot esculenta | F | FSK3 | Mes_Manes.05G140400 | 218 | 24687 | 5,35 | -0,306 | -1,548 | 6,4 |
| Angiosperms-Eudicots | Malpighiales | Manihot esculenta | H | HKS | Mes_Manes.08G154500 | 102 | 11470 | 7,94 | -0,433 | -1,883 | 13,7 |
| Angiosperms-Eudicots | Malpighiales | Manihot esculenta | H | HKS | Mes_Manes.09G138100 | 102 | 11515 | 6,69 | -0,351 | -2,083 | 14,7 |
| Angiosperms-Eudicots | Malpighiales | Manihot esculenta | Y | Y2SK2 | Mes_Manes.02G098200 | 159 | 17137 | 7,00 | -0,178 | -1,335 | 17,0 |
| Angiosperms-Eudicots | Malpighiales | Manihot esculenta | Y | Y3SK2 | Mes_Manes.04G074135 | 164 | 17697 | 6,13 | -0,234 | -1,437 | 14,6 |
| Angiosperms-Eudicots | Malpighiales | Manihot esculenta | Y | Y3SK2 | Mes_Manes.04G087233 | 164 | 17678 | 6,06 | -0,237 | -1,429 | 14,6 |
| Angiosperms-Eudicots | Malpighiales | Manihot esculenta | Y | Y3SK2 | Mes_Manes.11G091500 | 183 | 19926 | 6,10 | -0,240 | -1,431 | 12,0 |
| Angiosperms-Eudicots | Malpighiales | Populus trichocarpa | F | FSK2 | Ptr_Potri.005G248100 | 225 | 25623 | 5,13 | -0,344 | -1,620 | 6,2 |
| Angiosperms-Eudicots | Malpighiales | Populus trichocarpa | F* | K11 | Ptr_Potri.002G013200 | 542 | 60489 | 6,32 | -0,154 | -1,241 | 4,1 |
| Angiosperms-Eudicots | Malpighiales | Populus trichocarpa | H | HK4 | Ptr_Potri.013G062200 | 238 | 23816 | 9,90 | -0,132 | -1,032 | 33,6 |
| Angiosperms-Eudicots | Malpighiales | Populus trichocarpa | H | HK8 | Ptr_Potri.013G062301 | 389 | 40176 | 8,32 | -0,180 | -1,346 | 31,6 |

| | | | | | | | | | | | |
|---|---|---|---|---|---|---|---|---|---|---|---|
| Angiosperms-Eudicots | Malpighiales | Populus trichocarpa | H | HS | Ptr_Potri.013G062401 | 96 | 10679 | 8,66 | -0,397 | -1,995 | 17,7 |
| Angiosperms-Eudicots | Malpighiales | Populus trichocarpa | Y | Y3K | Ptr_Potri.004G158500 | 133 | 14013 | 5,15 | -0,119 | -0,995 | 14,3 |
| Angiosperms-Eudicots | Malpighiales | Populus trichocarpa | Y | Y3K2 | Ptr_Potri.009G120100 | 183 | 19465 | 6,45 | -0,121 | -1,099 | 14,2 |
| Angiosperms-Eudicots | Malpighiales | Salix purpurea | F | FSK3 | Spu_SapurV1A.0573s0050 | 225 | 25618 | 5,56 | -0,322 | -1,650 | 6,2 |
| Angiosperms-Eudicots | Malpighiales | Salix purpurea | F | FSK3 | Spu_SapurV1A.5414s0010 | 229 | 26132 | 5,57 | -0,332 | -1,686 | 6,1 |
| Angiosperms-Eudicots | Malpighiales | Salix purpurea | F* | K12 | Spu_SapurV1A.0016s0910 | 557 | 61600 | 5,31 | -0,177 | -1,235 | 5,4 |
| Angiosperms-Eudicots | Malpighiales | Salix purpurea | H | HK | Spu_SapurV1A.0564s0010 | 115 | 11889 | 8,47 | -0,206 | -1,391 | 29,6 |
| Angiosperms-Eudicots | Malpighiales | Salix purpurea | H | HKS | Spu_SapurV1A.0564s0030 | 143 | 14726 | 9,05 | -0,292 | -1,657 | 28,0 |
| Angiosperms-Eudicots | Malpighiales | Salix purpurea | H | HK6 | Spu_SapurV1A.0732s0100 | 334 | 34310 | 7,18 | -0,132 | -1,206 | 30,5 |
| Angiosperms-Eudicots | Malpighiales | Salix purpurea | H | HK9 | Spu_SapurV1A.2014s0020 | 492 | 51065 | 6,45 | -0,161 | -1,268 | 29,1 |
| Angiosperms-Eudicots | Malpighiales | Salix purpurea | H | HK3 | Spu_SapurV1A.2014s0030 | 214 | 22340 | 9,48 | -0,186 | -1,229 | 27,6 |
| Angiosperms-Eudicots | Malpighiales | Salix purpurea | H | HK10 | Spu_SapurV1A.2659s0010 | 556 | 57303 | 6,27 | -0,158 | -1,243 | 30,2 |
| Angiosperms-Eudicots | Malpighiales | Salix purpurea | Y | Y3K2 | Spu_SapurV1A.0432s0100 | 175 | 19113 | 6,14 | -0,151 | -1,157 | 13,1 |
| Angiosperms-Eudicots | Malpighiales | Salix purpurea | Y | Y3K2 | Spu_SapurV1A.5492s0010 | 175 | 19082 | 6,14 | -0,147 | -1,146 | 13,1 |
| Angiosperms-Eudicots | Rutaceae | Citrus clementina | F | FSK3 | Ccl_Ciclev10002349m | 234 | 26647 | 5,62 | -0,317 | -1,626 | 5,6 |
| Angiosperms-Eudicots | Rutaceae | Citrus clementina | F | FK3 | Ccl_Ciclev10029952m | 167 | 18708 | 5,44 | -0,241 | -1,405 | 7,8 |
| Angiosperms-Eudicots | Rutaceae | Citrus clementina | H | HK2S | Ccl_XP_006443476.2 | 135 | 14794 | 6,57 | -0,279 | -1,587 | 17,0 |
| Angiosperms-Eudicots | Rutaceae | Citrus clementina | H | HKS | Ccl_XP_024045887.1 | 99 | 11175 | 7,26 | -0,333 | -1,838 | 14,1 |
| Angiosperms-Eudicots | Rutaceae | Citrus clementina | Y | Y2SK2 | Ccl_Ciclev10026675m | 158 | 16564 | 9,40 | -0,143 | -1,141 | 18,4 |
| Angiosperms-Eudicots | Rutaceae | Citrus clementina | Y | Y3SK2 | Ccl_Ciclev10029285m | 208 | 21953 | 6,15 | -0,110 | -1,042 | 12,0 |
| Angiosperms-Eudicots | Malvaceae | Carica papaya | F | FSK3 | Cpa_evm.TU.supercontig_26.225 | 198 | 22635 | 5,10 | -0,330 | -1,585 | 4,5 |
| Angiosperms-Eudicots | Malvaceae | Carica papaya | H | HKS | Cpa_evm.model.supercontig_161.14 | 93 | 10502 | 6,62 | -0,405 | -1,984 | 16,1 |
| Angiosperms-Eudicots | Malvaceae | Carica papaya | Y | Y3SK2 | Cpa_evm.TU.supercontig_106.3 | 147 | 16018 | 6,12 | -0,219 | -1,378 | 8,8 |
| Angiosperms-Eudicots | Malvaceae | Carica papaya | Y | YSK2 | Cpa_evm.TU.supercontig_6.176 | 137 | 14760 | 9,45 | -0,173 | -1,222 | 11,7 |
| Angiosperms-Eudicots | Malvaceae | Gossypium raimondii | F | FSK2 | Gri_Gorai.002G119600 | 197 | 22221 | 5,57 | -0,304 | -1,582 | 6,6 |
| Angiosperms-Eudicots | Malvaceae | Gossypium raimondii | F | FSK3 | Gri_Gorai.008G038700 | 172 | 19275 | 6,12 | -0,321 | -1,742 | 11,0 |
| Angiosperms-Eudicots | Malvaceae | Gossypium raimondii | F | FSK3 | Gri_Gorai.009G189500 | 217 | 24121 | 5,21 | -0,271 | -1,447 | 7,4 |
| Angiosperms-Eudicots | Malvaceae | Gossypium raimondii | H | HKS | Gri_Gorai.007G257100 | 202 | 24003 | 6,61 | -0,569 | -2,572 | 7,9 |
| Angiosperms-Eudicots | Malvaceae | Gossypium raimondii | Y | Y3K | Gri_Gorai.005G245900 | 161 | 17949 | 4,79 | -0,146 | -1,052 | 5,6 |
| Angiosperms-Eudicots | Malvaceae | Gossypium raimondii | Y | Y2SK2 | Gri_Gorai.007G199900 | 135 | 14716 | 9,49 | -0,243 | -1,444 | 14,1 |
| Angiosperms-Eudicots | Malvaceae | Gossypium raimondii | Y | Y3SK2 | Gri_Gorai.008G030800 | 195 | 21975 | 6,26 | -0,2 | -1,343 | 8,2 |
| Angiosperms-Eudicots | Malvaceae | Gossypium raimondii | Y | Y3SK2 | Gri_Gorai.012G154800 | 199 | 21701 | 9,24 | -0,114 | -1,052 | 8,5 |
| Angiosperms-Eudicots | Brasicaceae | Arabidopsis thaliana | F | FSK3 | Ath_AT1G20440 | 265 | 29896 | 4,74 | -0,277 | -1,249 | 3,8 |
| Angiosperms-Eudicots | Brasicaceae | Arabidopsis thaliana | F | FSK3 | Ath_AT1G20450 | 260 | 29547 | 5,11 | -0,253 | -1,348 | 5,0 |
| Angiosperms-Eudicots | Brasicaceae | Arabidopsis thaliana | F | FSK2 | Ath_AT1G76180 | 185 | 20786 | 5,40 | -0,199 | -1,265 | 4,9 |
| Angiosperms-Eudicots | Brasicaceae | Arabidopsis thaliana | F | FK2 | Ath_AT4G38410 | 163 | 18255 | 5,88 | -0,293 | -1,629 | 8,0 |
| Angiosperms-Eudicots | Brasicaceae | Arabidopsis thaliana | H | HKS | Ath_AT1G54410 / XP_020868341 | 98 | 10795 | 6,67 | -0,357 | -1,868 | 15,3 |
| Angiosperms-Eudicots | Brasicaceae | Arabidopsis thaliana | Y | Y3SK2 | Ath_AT2G21490 | 185 | 19297 | 6,38 | -0,100 | -1,032 | 11,4 |
| Angiosperms-Eudicots | Brasicaceae | Arabidopsis thaliana | Y | YSK2 | Ath_AT3G50980 | 128 | 13434 | 9,19 | -0,108 | -1,053 | 16,4 |
| Angiosperms-Eudicots | Brasicaceae | Arabidopsis thaliana | Y | Y2K | Ath_AT4G39130 | 151 | 16259 | 6,09 | -0,031 | -0,774 | 9,9 |
| Angiosperms-Eudicots | Brasicaceae | Arabidopsis thaliana | Y | Y2SK2 | Ath_AT5G66400 | 186 | 18463 | 7,10 | -0,124 | -1,182 | 33,9 |
| Angiosperms-Eudicots | Brasicaceae | Arabidopsis thaliana | Y* | K6 | Ath_AT3G50970 | 193 | 20909 | 9,38 | -0,153 | -1,173 | 15,5 |
| Angiosperms-Eudicots | Brassicaceae | Arabidopsis lyrata | F | FSK3 | Aly_AL1G33370 | 264 | 29793 | 4,76 | -0,295 | -1,317 | 4,2 |
| Angiosperms-Eudicots | Brassicaceae | Arabidopsis lyrata | F | FSK3 | Aly_AL1G33380 | 256 | 29249 | 5,30 | -0,273 | -1,449 | 5,1 |
| Angiosperms-Eudicots | Brassicaceae | Arabidopsis lyrata | F | FSK2 | Aly_AL2G36240 | 170 | 19256 | 5,32 | -0,233 | -1,364 | 4,1 |
| Angiosperms-Eudicots | Brassicaceae | Arabidopsis lyrata | F | FSK2 | Aly_AL7G10280 | 164 | 18194 | 5,82 | -0,278 | -1,560 | 9,1 |
| Angiosperms-Eudicots | Brasicaceae | Arabidopsis lyrata | H | HKS | Aly_AL1G64750 | 98 | 10795 | 6,67 | -0,359 | -1,876 | 15,3 |
| Angiosperms-Eudicots | Brassicaceae | Arabidopsis lyrata | Y | Y3SK2 | Aly_AL4G10674 | 187 | 19662 | 6,29 | -0,097 | -1,007 | 10,2 |
| Angiosperms-Eudicots | Brassicaceae | Arabidopsis lyrata | Y | YSK2 | Aly_AL5G31400 | 127 | 13434 | 9,19 | -0,084 | -0,974 | 15,7 |
| Angiosperms-Eudicots | Brassicaceae | Arabidopsis lyrata | Y | Y3K | Aly_AL7G11070 | 161 | 17476 | 6,04 | -0,065 | -0,891 | 10,6 |
| Angiosperms-Eudicots | Brassicaceae | Arabidopsis lyrata | Y | Y3SK2 | Aly_AL8G45130 | 170 | 17545 | 7,96 | -0,157 | -1,269 | 28,2 |
| Angiosperms-Eudicots | Brassicaceae | Arabidopsis lyrata | Y* | K5 | Aly_AL5G31370 | 158 | 17103 | 6,78 | -0,148 | -1,219 | 15,8 |
| Angiosperms-Eudicots | Brassicaceae | Arabidopsis lyrata | Y* | K6 | Aly_AL5G31390 | 213 | 23053 | 9,34 | -0,178 | -1,264 | 17,8 |
| Angiosperms-Eudicots | Brassicaceae | Boechera stricta | F | FSK2 | Bst_Bostr.20129s0054 | 199 | 22446 | 5,34 | -0,232 | -1,368 | 4,0 |
| Angiosperms-Eudicots | Brassicaceae | Boechera stricta | F | FSK3 | Bst_Bostr.25542s0027 | 177 | 19694 | 5,54 | -0,296 | 1,555 | 7,9 |
| Angiosperms-Eudicots | Brassicaceae | Boechera stricta | F | FSK3 | Bst_Bostr.7128s0640 | 243 | 27518 | 5,55 | -0,265 | -1,437 | 3,7 |
| Angiosperms-Eudicots | Brassicaceae | Boechera stricta | F | FSK3 | Bst_Bostr.7128s0641 | 257 | 29493 | 5,67 | -0,300 | -1,575 | 4,3 |
| Angiosperms-Eudicots | Brasicaceae | Boechera stricta | H | HKS | Bst_Bostr.13404s0009 | 99 | 11003 | 6,70 | -0,371 | -1,913 | 14,1 |
| Angiosperms-Eudicots | Brassicaceae | Boechera stricta | Y | Y2SK2 | Bst_Bostr.0568s0050 | 190 | 18587 | 6,70 | -0,080 | -1,022 | 33,2 |
| Angiosperms-Eudicots | Brassicaceae | Boechera stricta | Y | Y3K | Bst_Bostr.25542s0101 | 149 | 16335 | 6,09 | -0,086 | -0,949 | 9,4 |
| Angiosperms-Eudicots | Brassicaceae | Boechera stricta | Y | Y3SK2 | Bst_Bostr.5022s0083 | 191 | 20250 | 6,56 | -0,108 | -1,079 | 9,9 |
| Angiosperms-Eudicots | Brassicaceae | Boechera stricta | Y | YSK2 | Bst_Bostr.6864s0146 | 127 | 13457 | 9,22 | -0,117 | -1,083 | 16,5 |
| Angiosperms-Eudicots | Brassicaceae | Boechera stricta | Y* | K5 | Bst_Bostr.6864s0147 | 216 | 23044 | 9,16 | -0,127 | -1,132 | 16,2 |

| | | | | | | | | | | | |
|---|---|---|---|---|---|---|---|---|---|---|---|
| Angiosperms-Eudicots | Brassicaceae | Capsella grandiflora | F | FSK2 | Cgr_Cagra.0799s0088 | 197 | 22120 | 5,41 | -0,226 | -1,364 | 4,6 |
| Angiosperms-Eudicots | Brassicaceae | Capsella grandiflora | F | FSK3 | Cgr_Cagra.1383s0028 | 175 | 19484 | 5,36 | -0,268 | -1,425 | 8,6 |
| Angiosperms-Eudicots | Brassicaceae | Capsella grandiflora | F | FSK3 | Cgr_Cagra.25489s0001 | 262 | 29373 | 4,95 | -0,256 | -1,289 | 5,0 |
| Angiosperms-Eudicots | Brassicaceae | Capsella grandiflora | F | FSK3 | Cgr_Cagra.25489s0002 | 264 | 30105 | 5,46 | -0,274 | -1,472 | 4,5 |
| Angiosperms-Eudicots | Brasicaceae | Capsella grandiflora | H | HKS | Cgr_Cagra.27207s0001 | 100 | 11011 | 6,95 | -0,345 | -1,864 | 16,0 |
| Angiosperms-Eudicots | Brassicaceae | Capsella grandiflora | Y | YSK2 | Cgr_Cagra.0926s0063 | 120 | 12833 | 8,90 | -0,119 | -1,112 | 13,3 |
| Angiosperms-Eudicots | Brassicaceae | Capsella grandiflora | Y | Y2K | Cgr_Cagra.2374s0005 | 145 | 16277 | 5,97 | -0,155 | -1,146 | 6,9 |
| Angiosperms-Eudicots | Brassicaceae | Capsella grandiflora | Y* | K6 | Cgr_Cagra.0926s0062 | 204 | 21755 | 9,21 | -0,119 | -1,086 | 17,6 |
| Angiosperms-Eudicots | Brassicaceae | Capsella grandiflora | Y* | SK2 | Cgr_Cagra.2007s0059 | 143 | 14354 | 9,52 | -0,152 | -1,157 | 30,1 |
| Angiosperms-Eudicots | Brassicaceae | Eutrema salsugineum | F | FSK3 | Esa_Thhalv10008313m | 301 | 34146 | 5,55 | -0,184 | -1,224 | 4,3 |
| Angiosperms-Eudicots | Brassicaceae | Eutrema salsugineum | F | FSK3 | Esa_Thhalv10008706m | 224 | 25231 | 5,18 | -0,256 | -1,361 | 5,8 |
| Angiosperms-Eudicots | Brassicaceae | Eutrema salsugineum | F | FSK2 | Esa_Thhalv10019152m | 201 | 22323 | 5,42 | -0,218 | -1,341 | 6,0 |
| Angiosperms-Eudicots | Brassicaceae | Eutrema salsugineum | F | FSK3 | Esa_Thhalv10026303m | 195 | 21681 | 5,71 | -0,307 | -1,622 | 7,7 |
| Angiosperms-Eudicots | Brasicaceae | Eutrema salsugineum | H | HKS | Esa_Thhalv10023775m | 99 | 10939 | 6,73 | -0,334 | -1,793 | 15,2 |
| Angiosperms-Eudicots | Brassicaceae | Eutrema salsugineum | Y | Y3SK2 | Esa_Thhalv10000340m | 181 | 18973 | 6,56 | -0,077 | -0,975 | 11,6 |
| Angiosperms-Eudicots | Brassicaceae | Eutrema salsugineum | Y | Y2SK3 | Esa_Thhalv10004906m | 215 | 21416 | 7,14 | -0,089 | -1,068 | 30,2 |
| Angiosperms-Eudicots | Brassicaceae | Eutrema salsugineum | Y | YSK2 | Esa_Thhalv10010821m | 125 | 13132 | 8,69 | -0,041 | -0,862 | 15,2 |
| Angiosperms-Eudicots | Brassicaceae | Eutrema salsugineum | Y | Y2K | Esa_Thhalv10026916m | 152 | 16405 | 6,28 | -0,032 | -0,799 | 9,2 |
| Angiosperms-Eudicots | Fabidae | Cucumis sativus | F | FSK3 | Csat_Cucsa.077690 | 236 | 26625 | 5,17 | -0,324 | -1,525 | 6,8 |
| Angiosperms-Eudicots | Fabidae | Cucumis sativus | H | HKS | Csat_Cucsa.338040 | 101 | 11368 | 6,43 | -0,434 | -2,023 | 16,8 |
| Angiosperms-Eudicots | Fabidae | Cucumis sativus | Y | Y4SK2 | Csat_Cucsa.106380 | 177 | 19181 | 7,26 | -0,186 | -1,383 | 20,3 |
| Angiosperms-Eudicots | Fabidae | Glycine max | F | FSK3 | Gma_Glyma.04G009400 | 214 | 24164 | 5,53 | -0,291 | -1,509 | 7,0 |
| Angiosperms-Eudicots | Fabidae | Glycine max | H | HKS | Gma_Glyma.16G037900 | 112 | 12592 | 6,24 | -0,401 | -1,874 | 16,1 |
| Angiosperms-Eudicots | Fabidae | Glycine max | H | HKS | Gma_Glyma.16G038000 | 84 | 9442 | 6,21 | -0,419 | -1,942 | 16,7 |
| Angiosperms-Eudicots | Fabidae | Glycine max | H | HKS | Gma_Glyma.17G187600 | 90 | 10192 | 7,36 | -0,377 | -1,998 | 15,6 |
| Angiosperms-Eudicots | Fabidae | Glycine max | H | HKS | Gma_Glyma.19G114700 | 113 | 12606 | 6,36 | -0,381 | -1,842 | 16,8 |
| Angiosperms-Eudicots | Fabidae | Glycine max | Y | Y3SK2 | Gma_Glyma.04G009900 | 166 | 17319 | 9,22 | -0,077 | -0,971 | 14,5 |
| Angiosperms-Eudicots | Fabidae | Glycine max | Y | Y2K | Gma_Glyma.07G090400 | 243 | 25658 | 6,02 | -0,191 | -1,279 | 22,2 |
| Angiosperms-Eudicots | Fabidae | Glycine max | Y | Y2K | Gma_Glyma.09G185500 | 253 | 26630 | 6,29 | -0,167 | -1,244 | 19,8 |
| Angiosperms-Eudicots | Fabidae | Glycine max | Y | Y3K | Gma_Glyma.12G235800 | 135 | 14870 | 5,54 | -0,148 | -1,066 | 5,9 |
| Angiosperms-Eudicots | Fabidae | Glycine max | Y | Y3SK | Gma_Glyma.13G201300 | 139 | 15133 | 5,52 | -0,190 | -1,186 | 8,6 |
| Angiosperms-Eudicots | Fabidae | Glycine max | Y* | K | Gma_Glyma.08G048900 | 91 | 9917 | 6,64 | -0,098 | -1,062 | 9,9 |
| Angiosperms-Eudicots | Fabidae | Medicago truncatula | F | FSK2 | Mtr_Medtr3g117290 | 209 | 23528 | 5,52 | -0,272 | -1,475 | 5,3 |
| Angiosperms-Eudicots | Fabidae | Medicago truncatula | H | HKS | Mtr_Medtr6g027810 / XP_013451547 | 100 | 11104 | 7,28 | -0,337 | -2,010 | 15,0 |
| Angiosperms-Eudicots | Fabidae | Medicago truncatula | H | HKS | Mtr_Medtr7g086340 / XP_003624683 | 121 | 13071 | 6,26 | -0,38 | -1,682 | 24,0 |
| Angiosperms-Eudicots | Fabidae | Medicago truncatula | Y | Y3SK2 | Mtr_Medtr3g117190 | 196 | 20314 | 8,81 | -0,048 | -0,904 | 13,8 |
| Angiosperms-Eudicots | Fabidae | Medicago truncatula | Y | Y2K4 | Mtr_Medtr6g084640 | 312 | 31219 | 6,45 | -0,097 | -1,043 | 29,8 |
| Angiosperms-Eudicots | Fabidae | Medicago truncatula | Y* | K | Mtr_Medtr8g106140 | 85 | 9200 | 5,75 | -0,075 | -0,869 | 10,6 |
| Angiosperms-Eudicots | Fabidae | Phaseolus vulgaris | F | FSK2 | Phvul.009G004400 | 202 | 22973 | 5,32 | -0,300 | -1,558 | 6,4 |
| Angiosperms-Eudicots | Fabidae | Phaseolus vulgaris | H | HKS | Phvul.001G114100 | 107 | 12134 | 6,43 | -0,424 | -2,001 | 15,9 |
| Angiosperms-Eudicots | Fabidae | Phaseolus vulgaris | H | HKS | Phvul.004G051100 | 119 | 13831 | 6,88 | -0,480 | -2,277 | 14,3 |
| Angiosperms-Eudicots | Fabidae | Phaseolus vulgaris | Y | Y2K | Phvul.004G158800 | 259 | 26379 | 6,34 | -0,122 | -1,125 | 25,9 |
| Angiosperms-Eudicots | Fabidae | Phaseolus vulgaris | Y | Y3SK2 | Phvul.009G005300 | 177 | 18839 | 9,64 | -0,098 | -1,007 | 8,5 |
| Angiosperms-Eudicots | Fabidae | Prunus persica | F | FSK2 | Pper_Prupe.1G356400 | 254 | 28937 | 5,27 | -0,317 | -1,577 | 7,9 |
| Angiosperms-Eudicots | Fabidae | Prunus persica | F | FK3 | Pper_Prupe.8G173200 | 182 | 20409 | 6,35 | -0,294 | -1,642 | 10,4 |
| Angiosperms-Eudicots | Fabidae | Prunus persica | H | HS | Pper_Prupe.7G009800 | 85 | 9758 | 8,62 | -0,539 | -2,446 | 10,6 |
| Angiosperms-Eudicots | Fabidae | Prunus persica | Y | Y2K3 | Pper_Prupe.7G161000 | 268 | 28990 | 6,25 | -0,157 | -1,226 | 13,4 |
| Angiosperms-Eudicots | Fabidae | Prunus persica | Y | Y2K7 | Pper_Prupe.7G161100 | 380 | 39942 | 6,39 | -0,156 | -1,215 | 20,8 |
| Angiosperms-Eudicots | Fabidae | Prunus persica | Y | Y2SK3 | Pper_Prupe.7G161200 | 202 | 21447 | 7,95 | -0,146 | -1,235 | 16,8 |
| Angiosperms-Eudicots | Fabidae | Prunus persica | Y | Y3SK2 | Pper_Prupe.8G091000 | 206 | 21597 | 5,88 | -0,128 | -1,067 | 18,0 |
| Angiosperms-Eudicots | Fabidae | Trifolium pratense | F | FSK2 | Tpr_Tp57577_TGAC_v2_gene32200 | 214 | 24082 | 5,33 | -0,281 | -1,491 | 5,6 |
| Angiosperms-Eudicots | Fabidae | Trifolium pratense | H | HKS | Tpr_PNX72597.1 | 150 | 16090 | 6,07 | -0,305 | -1,573 | 26,0 |
| Angiosperms-Eudicots | Fabidae | Trifolium pratense | H | HKS | Tpr_Tp57577_TGAC_v2_mRNA30898 | 100 | 11176 | 7,95 | -0,410 | -2,072 | 14,0 |
| Angiosperms-Eudicots | Fabidae | Trifolium pratense | Y | Y3SK2 | Tpr_Tp57577_TGAC_v2_gene32191 | 184 | 18996 | 9,27 | -0,058 | -0,897 | 13,6 |
| Angiosperms-Eudicots | Fabidae | Trifolium pratense | Y | Y2K4 | Tpr_Tp57577_TGAC_v2_gene35228 | 314 | 31025 | 6,61 | -0,086 | -1,017 | 31,5 |
| Angiosperms-Eudicots | Fabidae | Trifolium pratense | Y* | K | Tpr_Tp57577_TGAC_v2_gene12814 | 97 | 10454 | 5,16 | -0,048 | -0,768 | 10,3 |

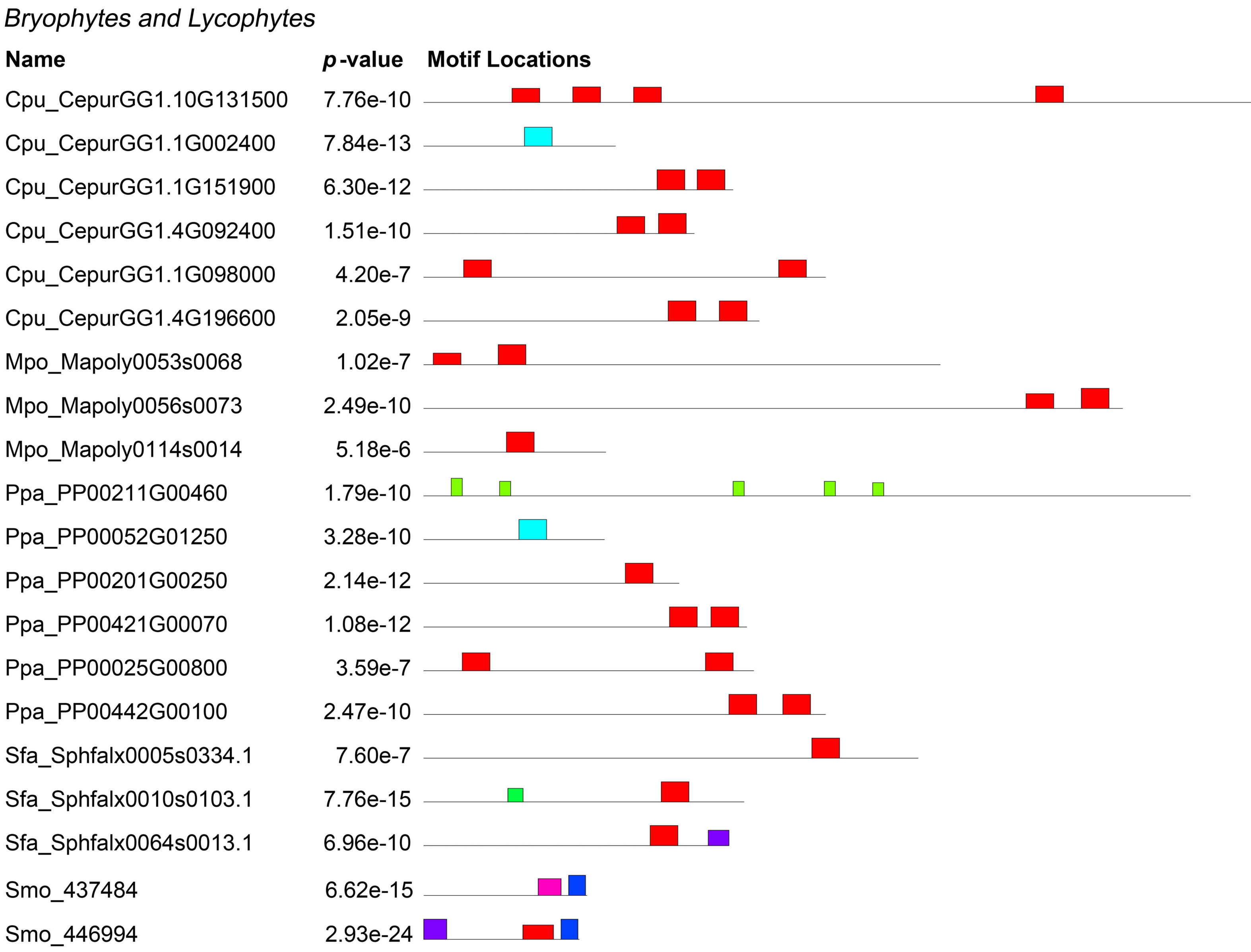

Bryophytes and Lycophytes
Name
p-value
Motif Locations
Cpu_CepurGG1.10G131500 7.76e-10
Cpu_CepurGG1.1G002400 7.84e-13
Cpu_CepurGG1.1G151900 6.30e-12
Cpu_CepurGG1.4G092400 1.51e-10
Cpu_CepurGG1.1G098000 4.20e-7
Cpu_CepurGG1.4G196600 2.05e-9
Mpo_Mapoly0053s0068 1.02e-7
Mpo_Mapoly0056s0073 2.49e-10
Mpo_Mapoly0114s0014 5.18e-6
Ppa_PP00211G00460 1.79e-10
Ppa_PP00052G01250 3.28e-10
Ppa_PP00201G00250 2.14e-12
Ppa_PP00421G00070 1.08e-12
Ppa_PP00025G00800 3.59e-7
Ppa_PP00442G00100 2.47e-10
Sfa_Sphfalx0005s0334.1 7.60e-7
Sfa_Sphfalx0010s0103.1 7.76e-15
Sfa_Sphfalx0064s0013.1 6.96e-10
Smo_437484 6.62e-15
Smo_446994 2.93e-24

*Gymnosperms*

| Name | *p*-value |
|---|---|
| Gbi_GBI00024041 | 2.44e-39 |
| Gbi_GBI00029052 | 1.29e-19 |
| Gbi_GBI00018450 | 3.77e-22 |
| Pab_MA_120345g0010 | 7.05e-31 |
| Pab_MA_130855g0010 | 7.26e-38 |
| Pab_MA_151075g0010 | 6.31e-30 |
| Pab_MA_187114g0010 | 9.58e-35 |
| Pab_MA_19049g0010 | 1.35e-4 |
| Pab_MA_205576g0010 | 3.38e-14 |
| Pab_MA_448618g0010 | 2.92e-5 |
| Pab_MA_474985g0010 | 3.85e-38 |
| Pab_MA_95995g0010 | 2.27e-17 |
| Pab_MA_104214g0010 | 8.77e-35 |
| Pab_MA_10430236g0010 | 1.21e-7 |
| Pab_MA_107783g0020 | 6.11e-12 |
| Pab_MA_144878g0010 | 1.00e-9 |
| Pab_MA_17462g0010 | 1.00e-9 |
| Pab_MA_2408574g0010 | 1.87e-13 |
| Pab_MA_43017g0010 | 1.76e-11 |
| Pab_MA_45025g0010 | 2.01e-11 |
| Pab_MA_4978040g0010 | 2.22e-11 |
| Pab_MA_59157g0010 | 2.62e-7 |
| Pab_MA_65580g0010 | 3.34e-29 |
| Pab_MA_816406g0020 | 7.15e-12 |

**Motif Locations**

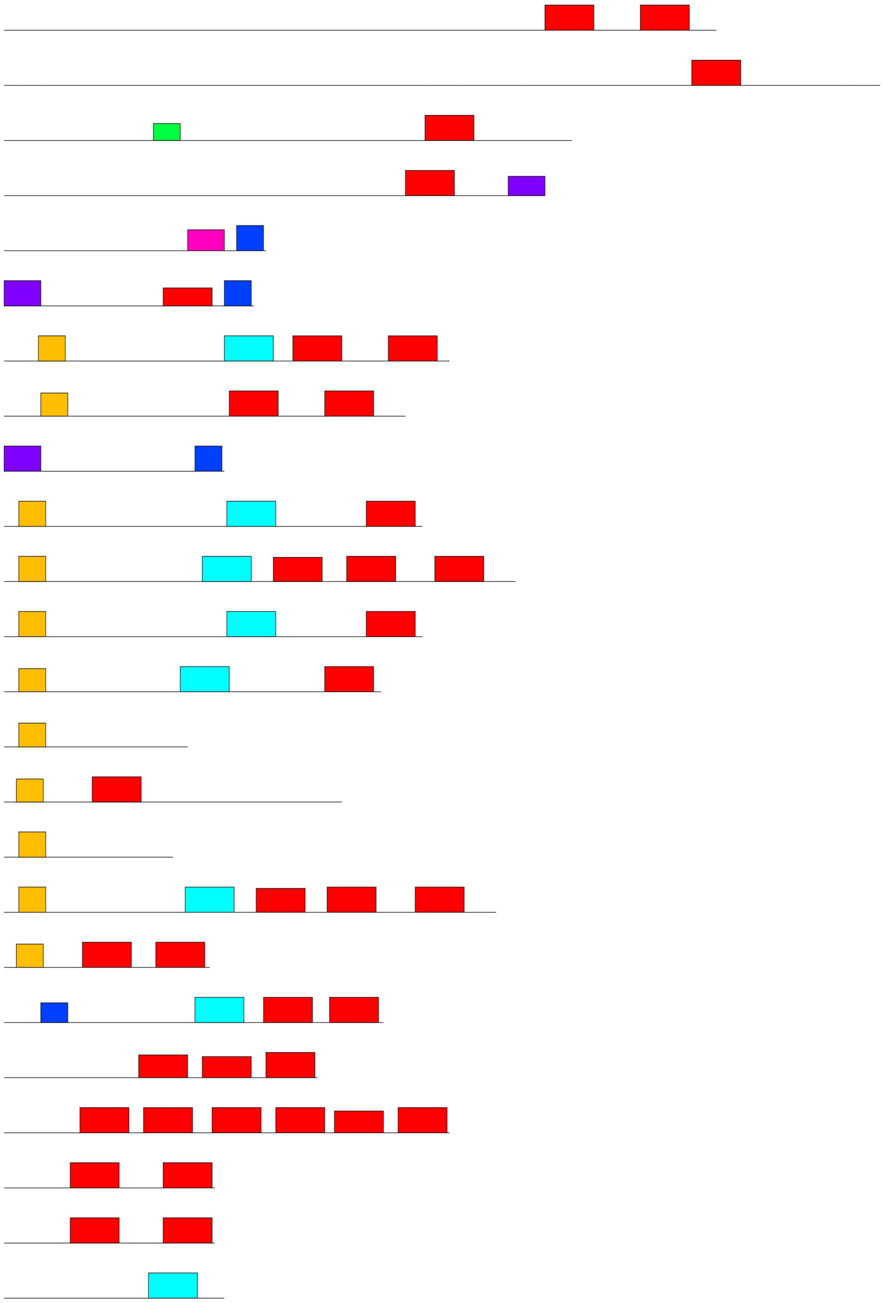

*Angiosperms-Basal*

| Name | *p*-value | Motif Locations |
|---|---|---|
| Atr_evm_27.TU.AmTr_v1.0_scaffold00001.295 | 2.78e-38 | |
| Atr_evm_27.model.AmTr_v1.0_scaffold00004.97 | 3.02e-35 | |
| Atr_evm_27.TU.AmTr_v1.0_scaffold00082.27 | 2.64e-32 | |
| Nco_Nycol.J01522 | 4.08e-28 | |
| Nco_Nycol.I01776 | 3.16e-39 | |
| Nco_Nycol.B01145/ | 3.16e-24 | |
| Nco_Nycol.B00068 | 1.63e-35 | |

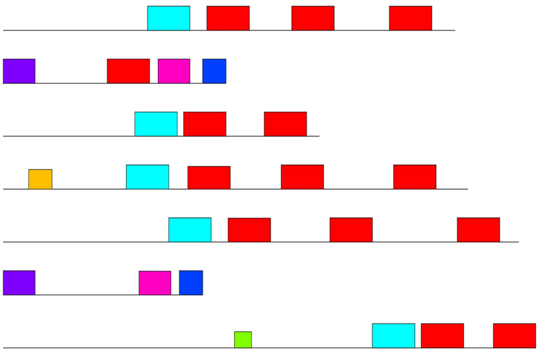

*Angiosperms-Monocots*

| Name | *p*-value | Motif Locations |
|---|---|---|
| Macu_GSMUA_Achr11G16760_001 | 2.96e-42 | |
| Macu_GSMUA_Achr4T21460_001 | 1.74e-33 | |
| Macu_GSMUA_Achr4G11310_001 | 2.88e-31 | |
| Aco_Aco011968 | 4.70e-42 | |
| Aco_OAY73247.1 | 5.86e-33 | |
| Aco_Aco016515 | 6.73e-36 | |
| Aco_Aco016518 | 1.85e-35 | |
| Aco_Aco016516 | 1.04e-29 | |
| Aco_Aco021122 | 6.74e-18 | |
| Aco_Aco021124 | 5.16e-28 | |
| Zmar_Zosma103g00290 | 2.36e-32 | |
| Zmar_Zosma440g00040 | 3.57e-40 | |
| Bdi_Bradi3g51200 | 3.59e-43 | |
| Bdi_Bradi5g10860 | 5.29e-43 | |
| Bdi_Bradi1g13330 | 4.29e-35 | |
| Bdi_Bradi1g37410 | 1.91e-43 | |
| Bdi_Bradi2g47575 | 3.14e-37 | |
| Bdi_Bradi3g43855 | 1.91e-43 | |
| Bdi_Bradi3g43870 | 3.14e-37 | |
| Bdi_Bradi4g19525 | 2.39e-42 | |
| Bdi_Bradi4g22280 | 8.75e-43 | |
| Bdi_Bradi4g22290 | 3.44e-39 | |
| Osa_LOC_Os02g44870 | 7.67e-31 | |
| Osa_LOC_Os03g45280 | 2.41e-32 | |
| Osa_LOC_Os01g50700 | 3.67e-43 | |
| Osa_LOC_Os11g26570 | 2.06e-37 | |
| Osa_LOC_Os11g26750 | 3.73e-31 | |
| Osa_LOC_Os11g26760 | 4.48e-39 | |
| Osa_LOC_Os11g26780 | 4.76e-40 | |
| Osa_LOC_Os11g26790 | 1.41e-44 | |

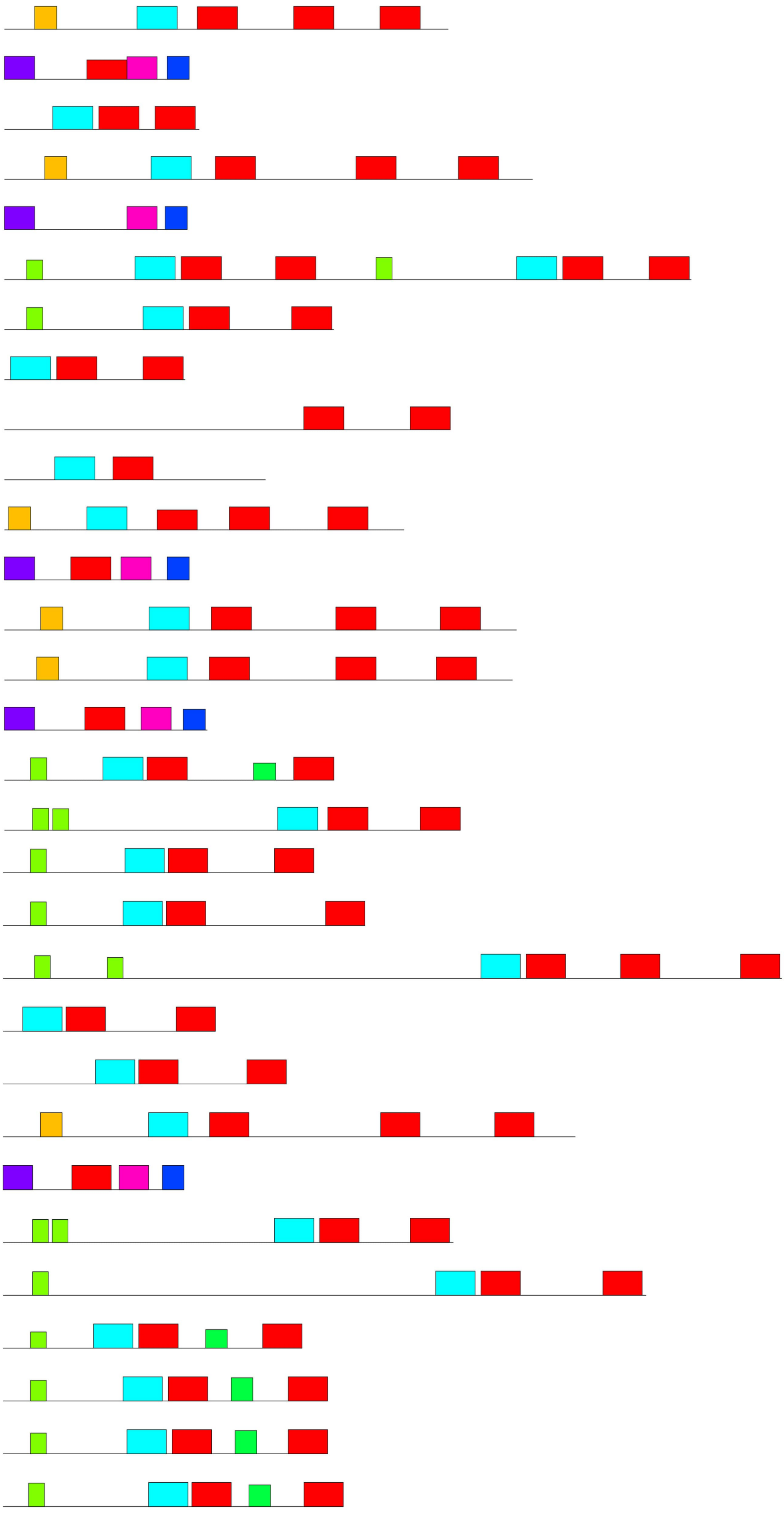

*(Continue to next page)*

*Angiosperms-Monocots*

| Name | *p*-value |
|---|---|
| Sbi_Sobic.004G286600 | 3.64e-43 |
| Sbi_Sobic.001G149500 | 1.79e-37 |
| Sbi_Sobic.003G270200 | 1.61e-40 |
| Sbi_Sobic.009G116700 | 3.88e-43 |
| Sbi_Sobic.010G041900 | 5.24e-38 |
| Zma_GRMZM2G147014 | 5.66e-41 |
| Zma_GRMZM2G373522 | 1.95e-40 |
| Zma_GRMZM2G169372 | 2.76e-35 |
| Zma_GRMZM2G448511 | 7.41e-37 |
| Zma_GRMZM2G052364 | 1.03e-35 |
| Zma_GRMZM2G079440 | 4.21e-46 |
| Zma_GRMZM2G098750 | 1.62e-37 |
| Pha_Pahal.A02866 | 8.18e-40 |
| Pha_Pahal.I02369 | 7.41e-40 |
| Pha_Pahal.E01980 | 1.04e-37 |
| Pha_Pahal.J00522 | 9.09e-46 |
| Pha_Pahal.J00526 | 1.17e-36 |
| Sit_Seita.1G267200 | 6.21e-40 |
| Sit_Seita.9G151800 | 2.34e-38 |
| Sit_Seita.5G166200 | 7.08e-41 |
| Sit_Seita.5G166300 | 2.10e-40 |
| Sit_Seita.5G286000 | 3.89e-35 |
| Sit_Seita.8G115200 | 2.56e-40 |
| Sit_Seita.8G115400 | 4.73e-44 |
| Sit_Seita.8G115100 | 1.23e-29 |

**Motif Locations**

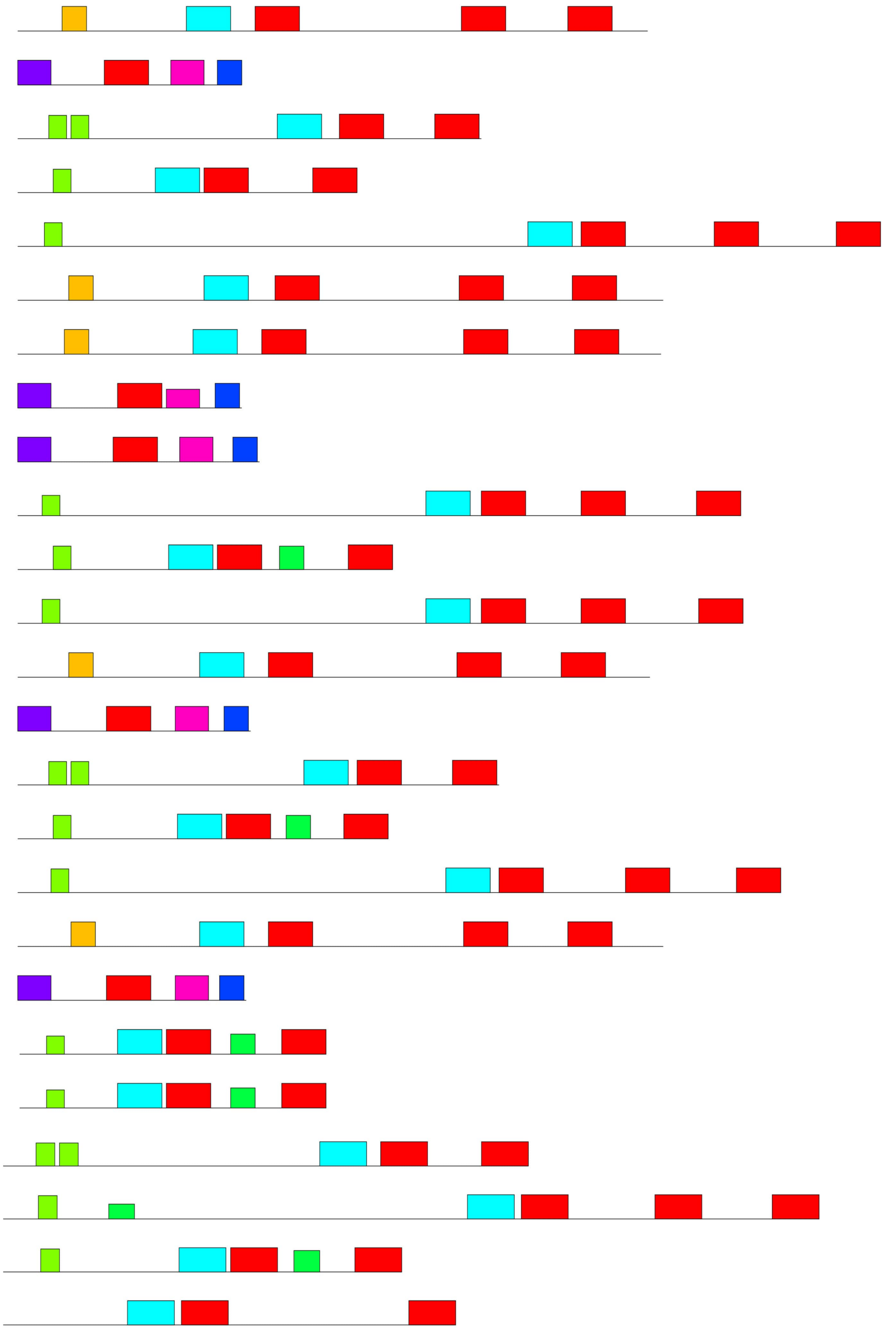

*Angiosperms-Eudicots*

| **Name** | ***p*-value** |
|---|---|
| Nnu_XP_010266960 | 7.30e-46 |
| Nnu_XP_019054435 | 1.02e-27 |
| Nnu_XP_010254713 | 9.78e-38 |
| Aqcoe5G125000 | 4.51e-34 |
| Aqcoe6G197600 | 6.00e-46 |
| Aqcoe5G125200 | 1.25e-15 |
| Aqcoe2G065100 | 2.06e-35 |
| Aqcoe5G186200 | 1.14e-14 |
| Aqcoe6G128700 | 5.21e-36 |
| Dca_DCAR_009522 | 1.92e-50 |
| Dca_XP_017246606.1 | 1.85e-25 |
| Dca_XP_017249516.1 | 3.69e-25 |
| Dca_DCAR_017310 | 1.92e-37 |
| Dca_DCAR_017311 | 2.77e-36 |
| Dca_DCAR_028967 | 8.44e-34 |
| Dca_DCAR_021479 | 3.99e-30 |
| Dca_DCAR_024077 | 6.02e-13 |
| Mgu_Migut.C00923 | 5.36e-42 |
| Mgu_Migut.J00768 | 1.52e-31 |
| Mgu_Migut.L01868 | 7.16e-30 |
| Mgu_Migut.B00209 | 3.54e-37 |
| Mgu_Migut.H02513 | 1.08e-35 |
| Stu_PGSC0003DMG400009968 | 7.40e-39 |
| Stu_XP_006355701.1 | 3.39e-37 |
| Stu_XP_006364877.1 | 2.43e-31 |
| Stu_XP_006365372.1 | 9.44e-23 |
| Stu_PGSC0003DMG400003530 | 3.43e-31 |
| Stu_PGSC0003DMG400003531 | 2.78e-44 |
| Stu_PGSC0003DMG400030949 | 1.48e-31 |
| Stu_PGSC0003DMG400034095 | 1.99e-34 |
| Solyc04g082200 | 3.90e-42 |
| Solyc_Solyc05g053070 | 1.41e-35 |
| Solyc01g109920 | 1.37e-33 |
| Solyc02g062390 | 6.22e-30 |
| Solyc02g084840 | 2.78e-44 |
| Solyc02g084850 | 1.39e-31 |
| Solyc01g065820 | 2.53e-14 |
| Solyc02g093255 | 1.22e-15 |
| Solyc09g074765 | 5.28e-14 |

**Motif Locations**

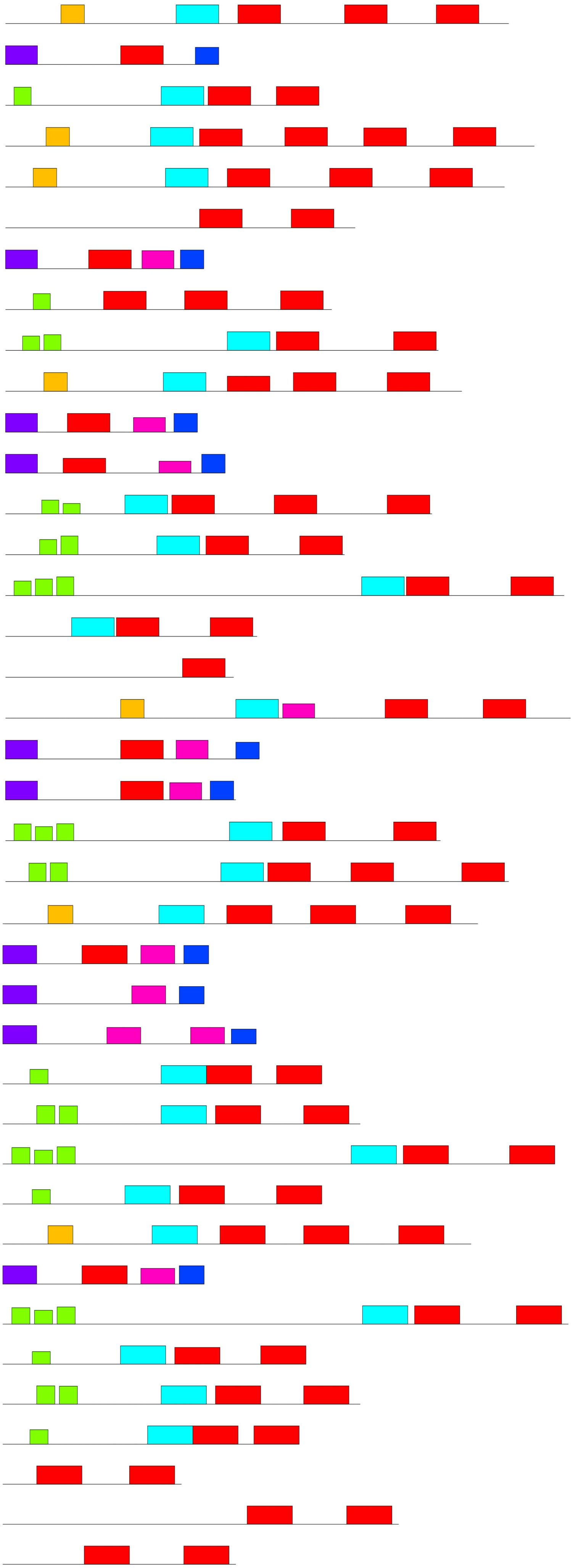

*(Continue to next page)*

*Angiosperms-Eudicots*

| Name | *p*-value |
|---|---|
| Kfe_Kaladp0040s0609 | 1.11e-38 |
| Kfe_Kaladp0477s0002 | 2.03e-44 |
| Kfe_Kaladp0098s0091 | 3.81e-34 |
| Kfe_Kaladp0911s0009 | 2.02e-36 |
| Kfe_Kaladp0018s0236 | 2.67e-33 |
| Kfe_Kaladp0040s0543 | 2.95e-42 |
| Kfe_Kaladp0047s0007 | 1.39e-31 |
| Egr_Eucgr.F01726 | 1.71e-25 |
| Egr_XP_010036245.1 | 1.55e-31 |
| Egr_Eucgr.I01292 | 1.13e-15 |
| Egr_Eucgr.I02392 | 1.58e-32 |
| Egr_Eucgr.I02395 | 4.42e-29 |
| Egr_Eucgr.J02380 | 2.69e-42 |
| Vvi_VIT_218s0001g00360 | 2.95e-51 |
| Vvi_NP_001268149.1 | 6.63e-30 |
| Vvi_VIT_203s0038g04390 | 7.38e-28 |
| Vvi_VIT_204s0023g02480 | 5.06e-26 |
| Rco_30170.t000735 | 9.50e-48 |
| Rco_30072.m000963 | 1.45e-34 |
| Rco_29634.t000016 | 4.86e-36 |
| Rco_29634.t000017 | 3.15e-37 |
| Rco_30131.t000233 | 1.80e-32 |
| Lus_Lus10005652.g | 4.68e-47 |
| Lus_Lus10021240.g | 1.47e-45 |
| Lus_Lus10021827.g | 3.48e-50 |
| Lus_Lus10034568.g | 8.83e-49 |
| Lus_Lus10022643.g | 1.34e-23 |
| Lus_Lus10020271.g | 6.91e-35 |
| Lus_Lus10014280.g | 1.11e-38 |
| Lus_Lus10017977.g | 2.80e-22 |
| Lus_Lus10025983.g | 6.67e-40 |
| Lus_Lus10041969.g | 2.37e-22 |
| Mes_Manes.05G140400 | 4.40e-45 |
| Mes_Manes.08G154500 | 2.51e-30 |
| Mes_Manes.09G138100 | 1.75e-38 |
| Mes_Manes.02G098200 | 5.11e-38 |
| Mes_Manes.04G074135 | 8.07e-32 |
| Mes_Manes.04G087233 | 4.31e-32 |
| Mes_Manes.11G091500 | 2.52e-30 |

**Motif Locations**

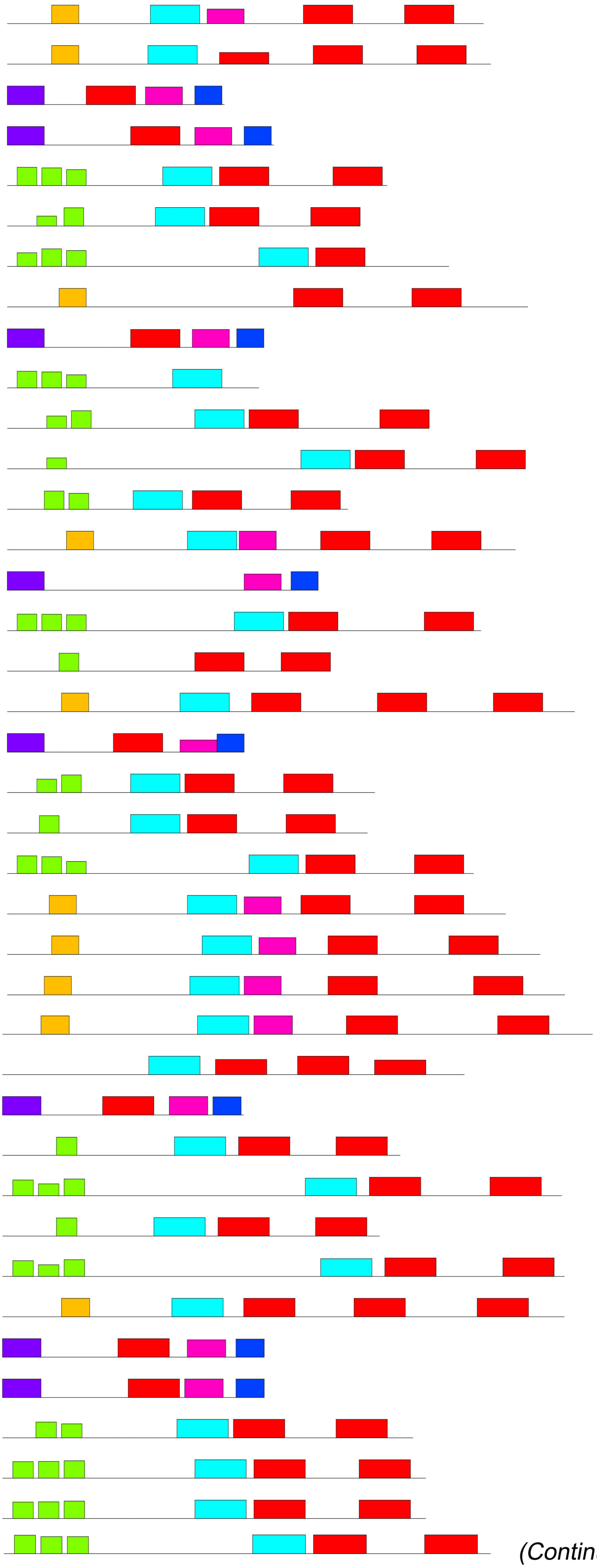

(Continue to next page)

*Angiosperms-Eudicots*

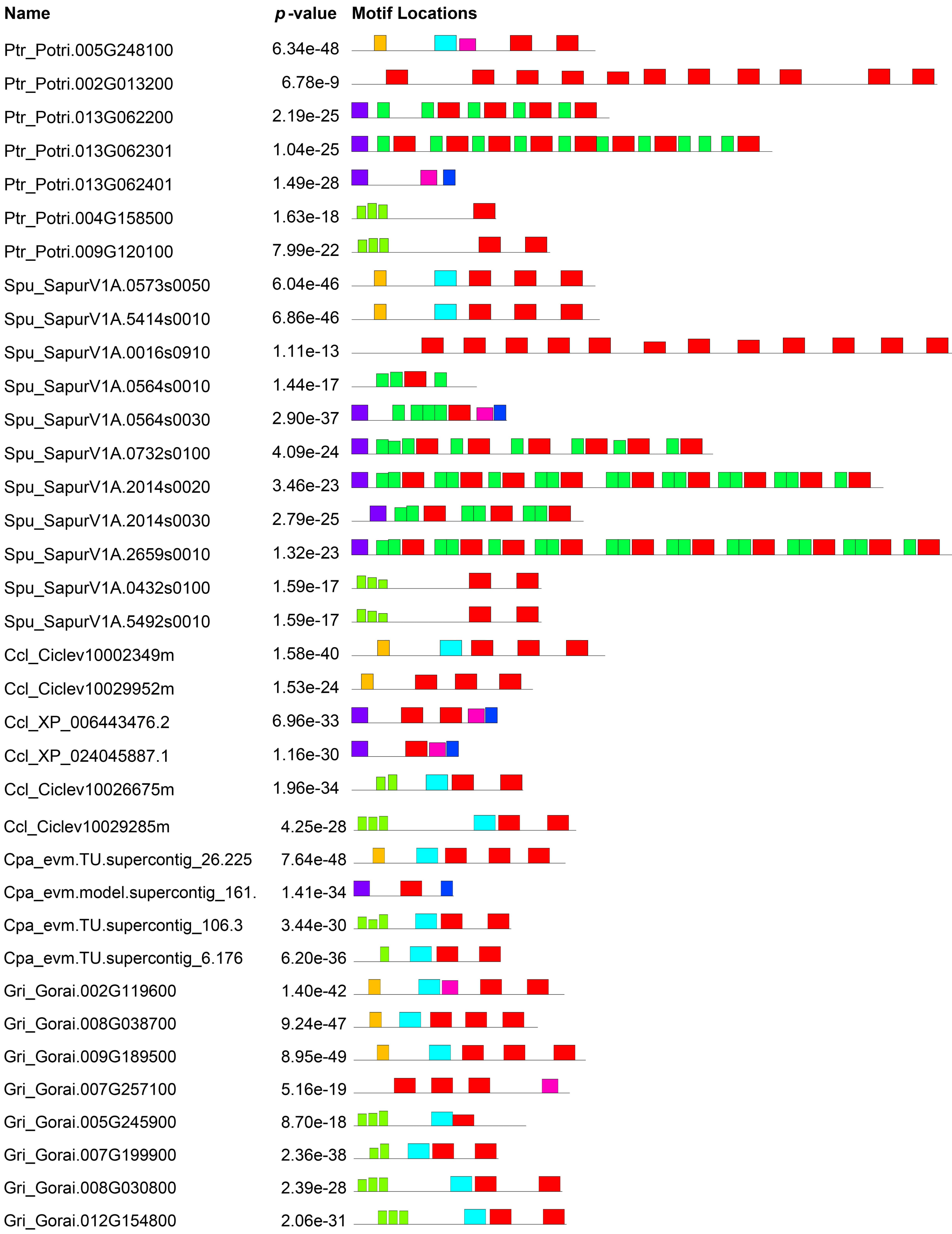


*(Continue to next page)*

*Angiosperms-Eudicots*

| **Name** | ***p*-value** |
|---|---|
| Ath_AT1G20440 | 9.04e-44 |
| Ath_AT1G20450 | 5.07e-50 |
| Ath_AT1G76180 | 1.22e-50 |
| Ath_AT4G38410 | 5.92e-34 |
| Ath_AT1G54410 | 4.42e-38 |
| Ath_AT2G21490 | 1.26e-38 |
| Ath_AT3G50980 | 4.94e-41 |
| Ath_AT4G39130 | 1.57e-14 |
| Ath_AT5G66400 | 1.09e-37 |
| Ath_AT3G50970 | 2.13e-9 |
| Aly_AL1G33370 | 8.21e-40 |
| Aly_AL1G33380 | 4.92e-52 |
| Aly_AL2G36240 | 2.90e-50 |
| Aly_AL7G10280 | 4.29e-30 |
| Aly_AL1G64750 | 4.49e-37 |
| Aly_AL4G10674 | 2.26e-38 |
| Aly_AL5G31400 | 1.97e-39 |
| Aly_AL7G11070 | 7.48e-15 |
| Aly_AL8G45130 | 1.80e-38 |
| Aly_AL5G31370 | 2.31e-9 |
| Aly_AL5G31390 | 4.00e-10 |
| Bst_Bostr.20129s0054 | 1.64e-51 |
| Bst_Bostr.25542s0027 | 6.59e-29 |
| Bst_Bostr.7128s0640 | 3.67e-41 |
| Bst_Bostr.7128s0641 | 3.91e-50 |
| Bst_Bostr.13404s0009 | 4.24e-37 |
| Bst_Bostr.0568s0050 | 2.55e-34 |
| Bst_Bostr.25542s0101 | 2.66e-13 |
| Bst_Bostr.5022s0083 | 7.50e-37 |
| Bst_Bostr.6864s0146 | 1.90e-33 |
| Bst_Bostr.6864s0147 | 1.27e-8 |
| Cgr_Cagra.0799s0088 | 1.30e-51 |
| Cgr_Cagra.1383s0028 | 1.46e-26 |
| Cgr_Cagra.25489s0001 | 7.90e-43 |
| Cgr_Cagra.25489s0002 | 3.31e-50 |
| Cgr_Cagra.27207s0001 | 2.23e-38 |
| Cgr_Cagra.0926s0063 | 5.58e-38 |

**Motif Locations**

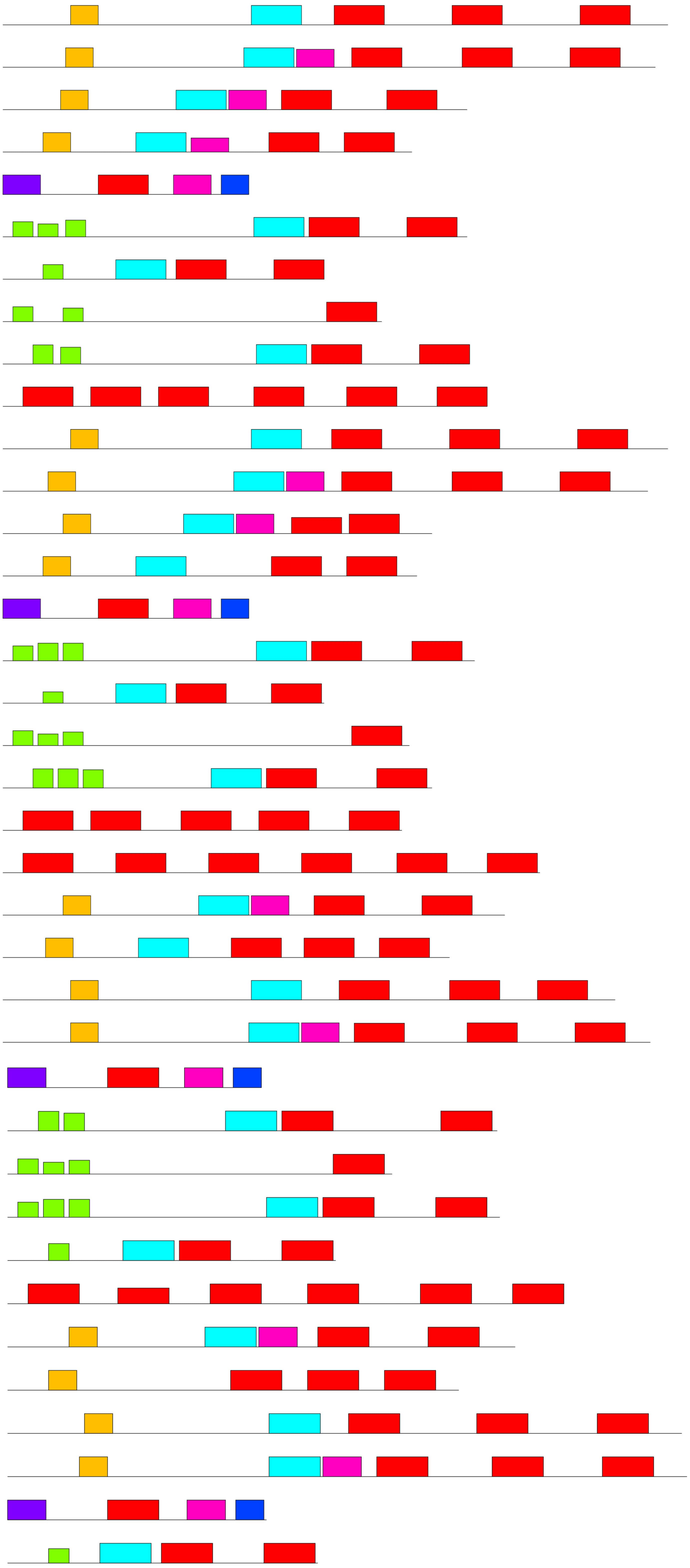

*(Continue to next page)*

*Angiosperms-Eudicots*

| **Name** | ***p*-value** |
|---|---|
| Cgr_Cagra.2374s0005 | 6.40e-15 |
| Cgr_Cagra.0926s0062 | 2.94e-10 |
| Cgr_Cagra.2007s0059 | 1.66e-32 |
| Esa_Thhalv10008313m | 7.52e-47 |
| Esa_Thhalv10008706m | 3.55e-36 |
| Esa_Thhalv10019152m | 2.42e-51 |
| Esa_Thhalv10026303m | 7.39e-33 |
| Esa_Thhalv10023775m | 3.35e-37 |
| Esa_Thhalv10000340m | 1.55e-39 |
| Esa_Thhalv10004906m | 1.36e-36 |
| Esa_Thhalv10010821m | 2.67e-36 |
| Esa_Thhalv10026916m | 6.79e-16 |
| Csat_Cucsa.077690 | 1.01e-44 |
| Csat_Cucsa.338040 | 9.69e-32 |
| Csat_Cucsa.106380 | 1.15e-39 |
| Gma_Glyma.04G009400 | 3.03e-49 |
| Gma_Glyma.16G037900 | 1.66e-37 |
| Gma_Glyma.16G038000 | 9.15e-32 |
| Gma_Glyma.17G187600 | 2.49e-35 |
| Gma_Glyma.19G114700 | 1.36e-34 |
| Gma_Glyma.04G009900 | 1.94e-36 |
| Gma_Glyma.07G090400 | 5.94e-20 |
| Gma_Glyma.09G185500 | 8.39e-22 |
| Gma_Glyma.12G235800 | 2.11e-18 |
| Gma_Glyma.13G201300 | 1.06e-25 |
| Gma_Glyma.08G048900 | 1.88e-11 |
| Mtr_Medtr3g117290 | 6.30e-48 |
| Mtr_Medtr6g027810 | 4.21e-37 |
| Mtr_Medtr7g086340 | 4.26e-32 |
| Mtr_Medtr3g117190 | 1.90e-35 |
| Mtr_Medtr6g084640 | 3.44e-19 |
| Mtr_Medtr8g106140 | 1.92e-13 |
| Phvul.009G004400 | 1.16e-49 |
| Phvul.001G114100 | 5.32e-35 |
| Phvul.004G051100 | 1.09e-30 |
| Phvul.004G158800 | 3.17e-17 |
| Phvul.009G005300 | 2.74e-36 |

**Motif Locations**

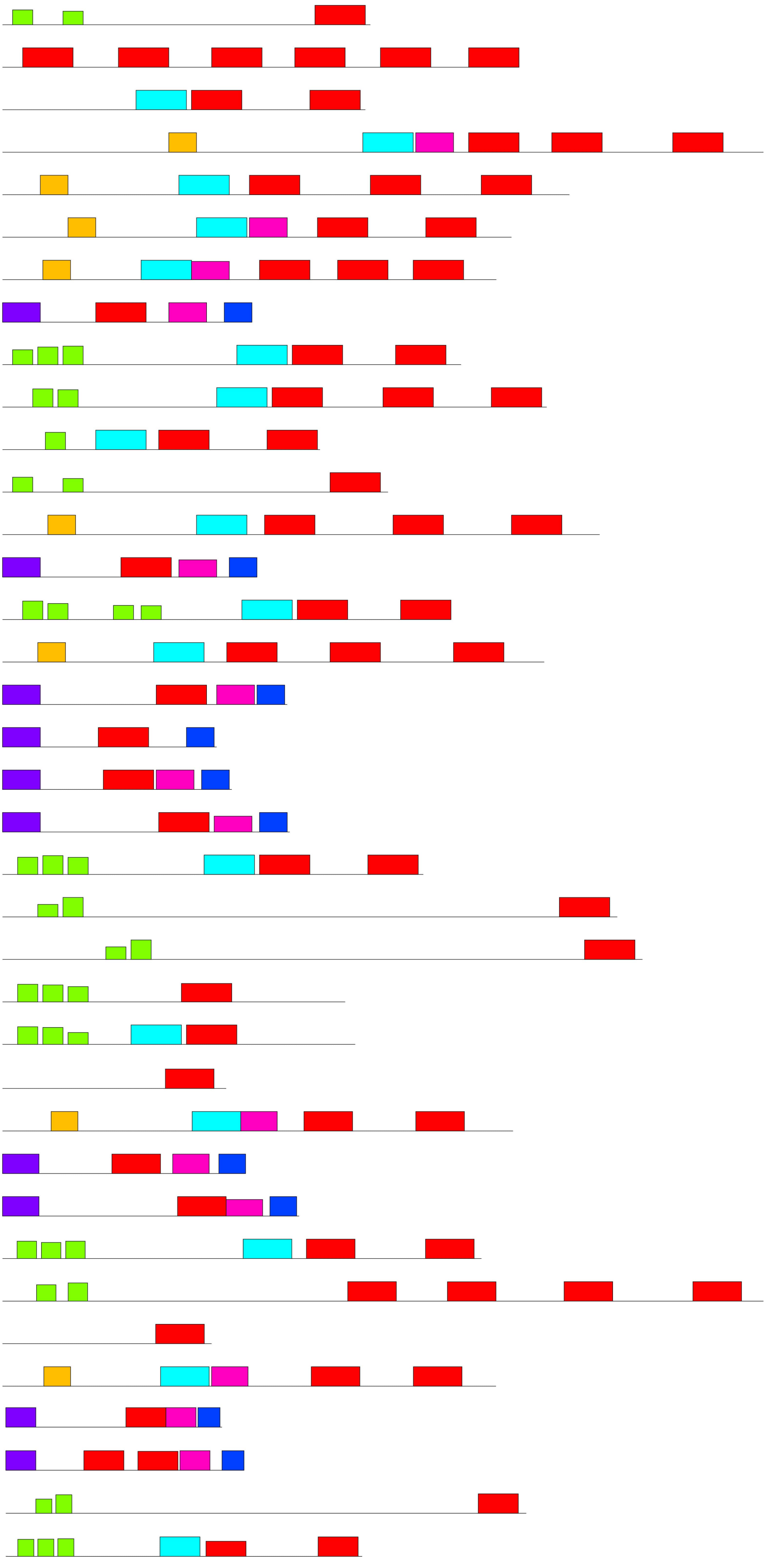

*(Continue to next page)*

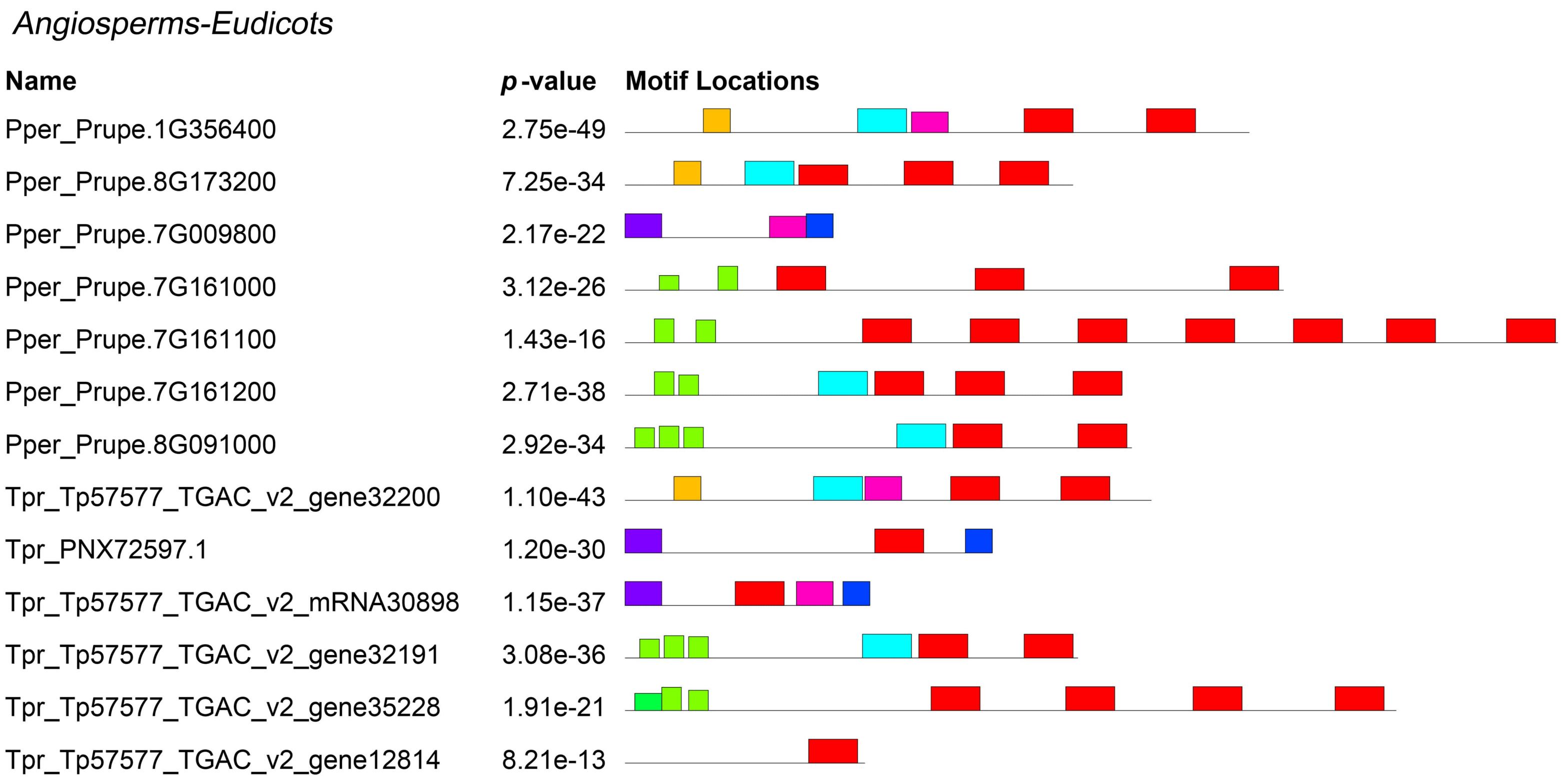
Angiosperms-Eudicots
Name
p-value
Motif Locations
Pper_Prupe.1G356400 2.75e-49
Pper_Prupe.8G173200 7.25e-34
Pper_Prupe.7G009800 2.17e-22
Pper_Prupe.7G161000 3.12e-26
Pper_Prupe.7G161100 1.43e-16
Pper_Prupe.7G161200 2.71e-38
Pper_Prupe.8G091000 2.92e-34
Tpr_Tp57577_TGAC_v2_gene32200 1.10e-43
Tpr_PNX72597.1 1.20e-30
Tpr_Tp57577_TGAC_v2_mRNA30898 1.15e-37
Tpr_Tp57577_TGAC_v2_gene32191 3.08e-36
Tpr_Tp57577_TGAC_v2_gene35228 1.91e-21
Tpr_Tp57577_TGAC_v2_gene12814 8.21e-13

**Supplementary Figures S2-S8 for Melgar and Zelada (2021)**

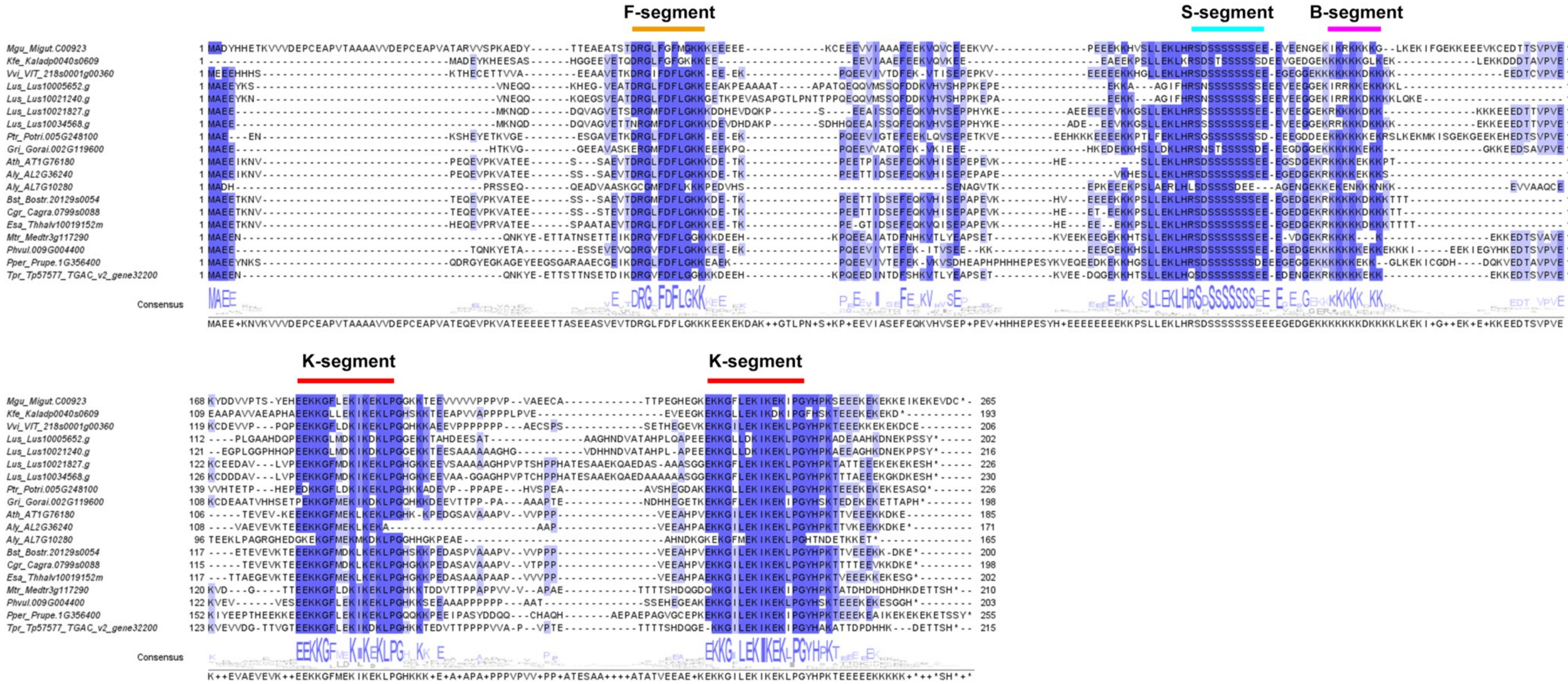


**Figure S2. Multiple sequence alignment of FSK2 dehydrins.** Protein sequences of FSK2 from eudicot species were aligned with Clustal Omega and visualised with Jalview. Structural segments are indicated and a consensus sequence is shown below the alignment.

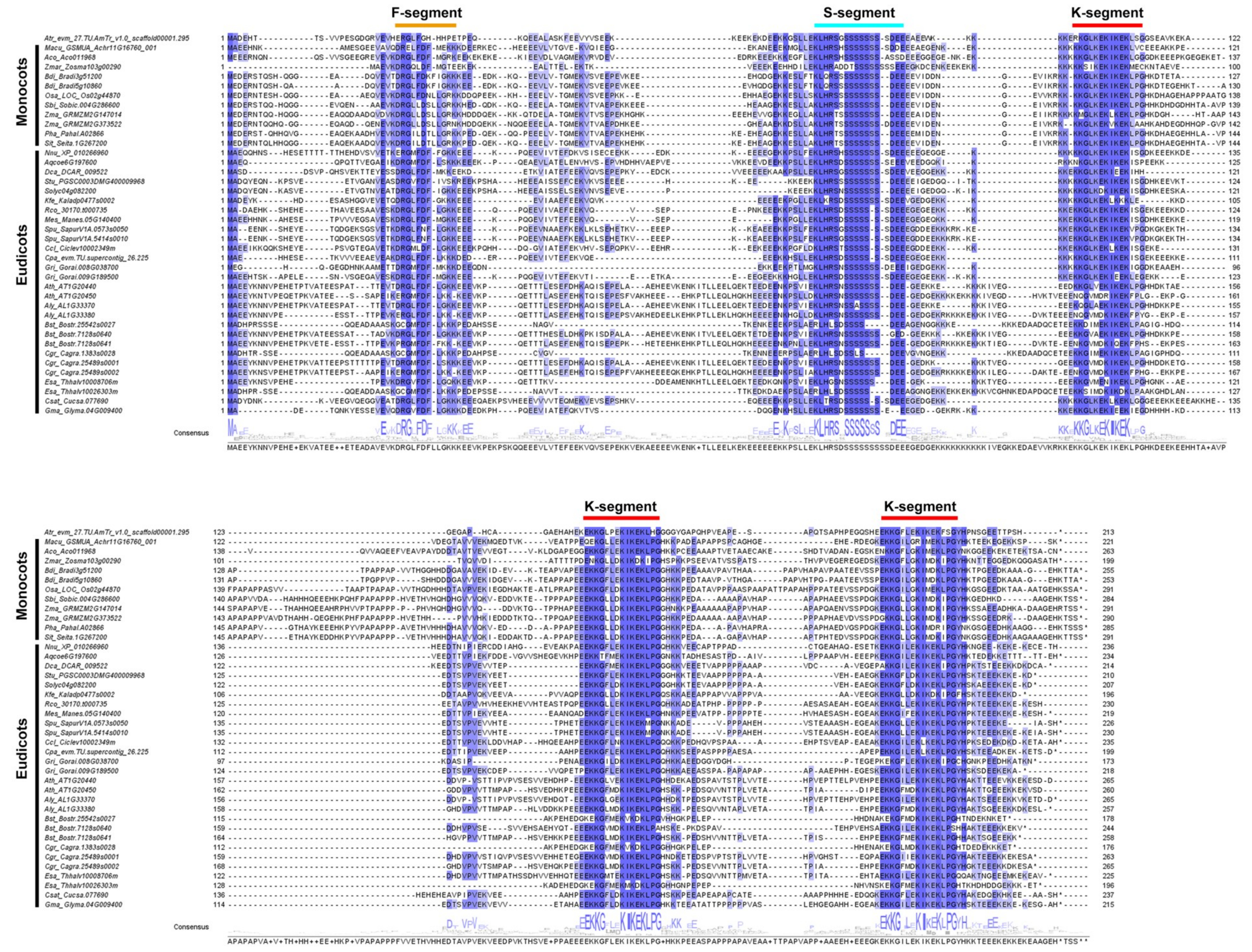


**Figure S3. Multiple sequences alignment of FSK3 dehydrins.** Protein sequences of FSK3-DHNs from angiosperms were aligned with Clustal Omega and visualised with Jalview. Structural segments are indicated and a consensus sequence is shown below the alignment. Note that there is a lysine-rich region adjacent to the S-segment but it is not as conserved as the B-segment found in FSK2-DHNs (compare to Fig. S2).

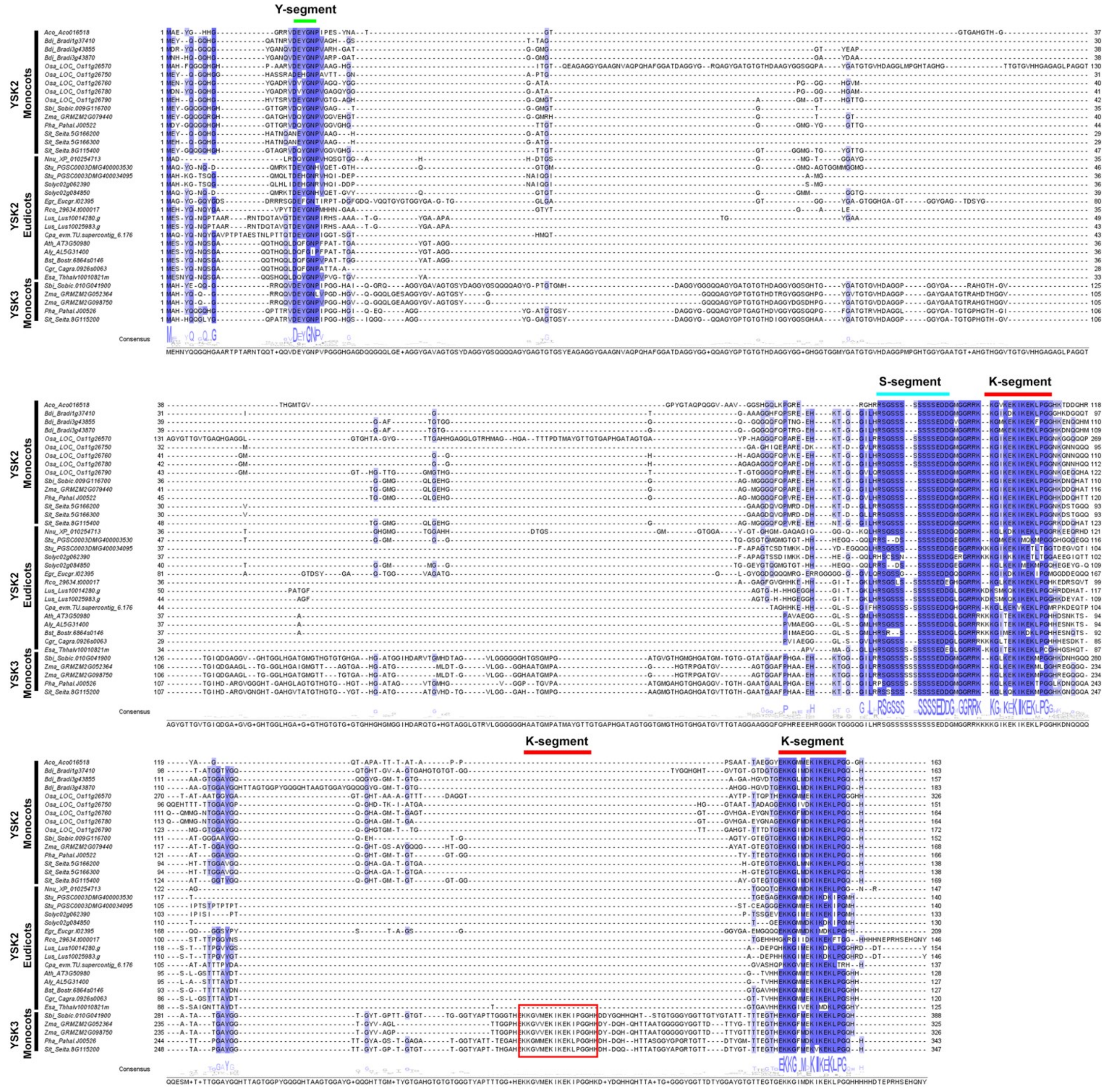

**Figure S4. Multiple sequences alignment of YSKn dehydrins.** Protein sequences of YSKn-DHNs from angiosperms were aligned with T-Coffee and visualised with Jalview. Structural segments are indicated and a consensus sequence is shown below the alignment.

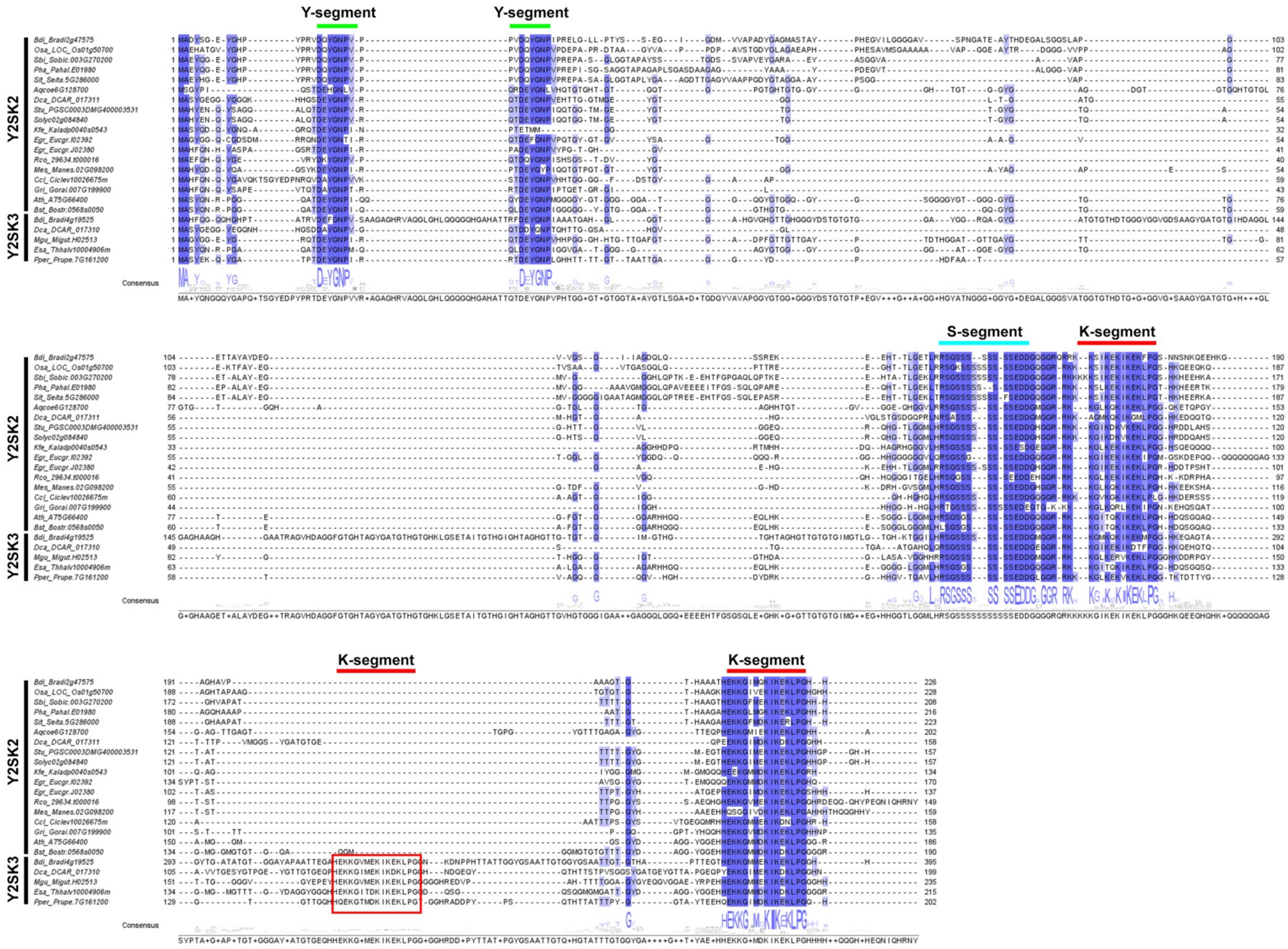


**Figure S5. Multiple sequence alignment of dehydrins Y2SKn.** Protein sequences of Y2SKn-DHNs from angiosperms were aligned with T-Coffee and visualised with Jalview. Structural segments are indicated and a consensus sequence is shown below the alignment. Y2SK2 and Y2SK3-DHNs are present in eudicots and the grass *Brachypodium distachyon*, while other Poaceae only have Y2SK2-DHNs.

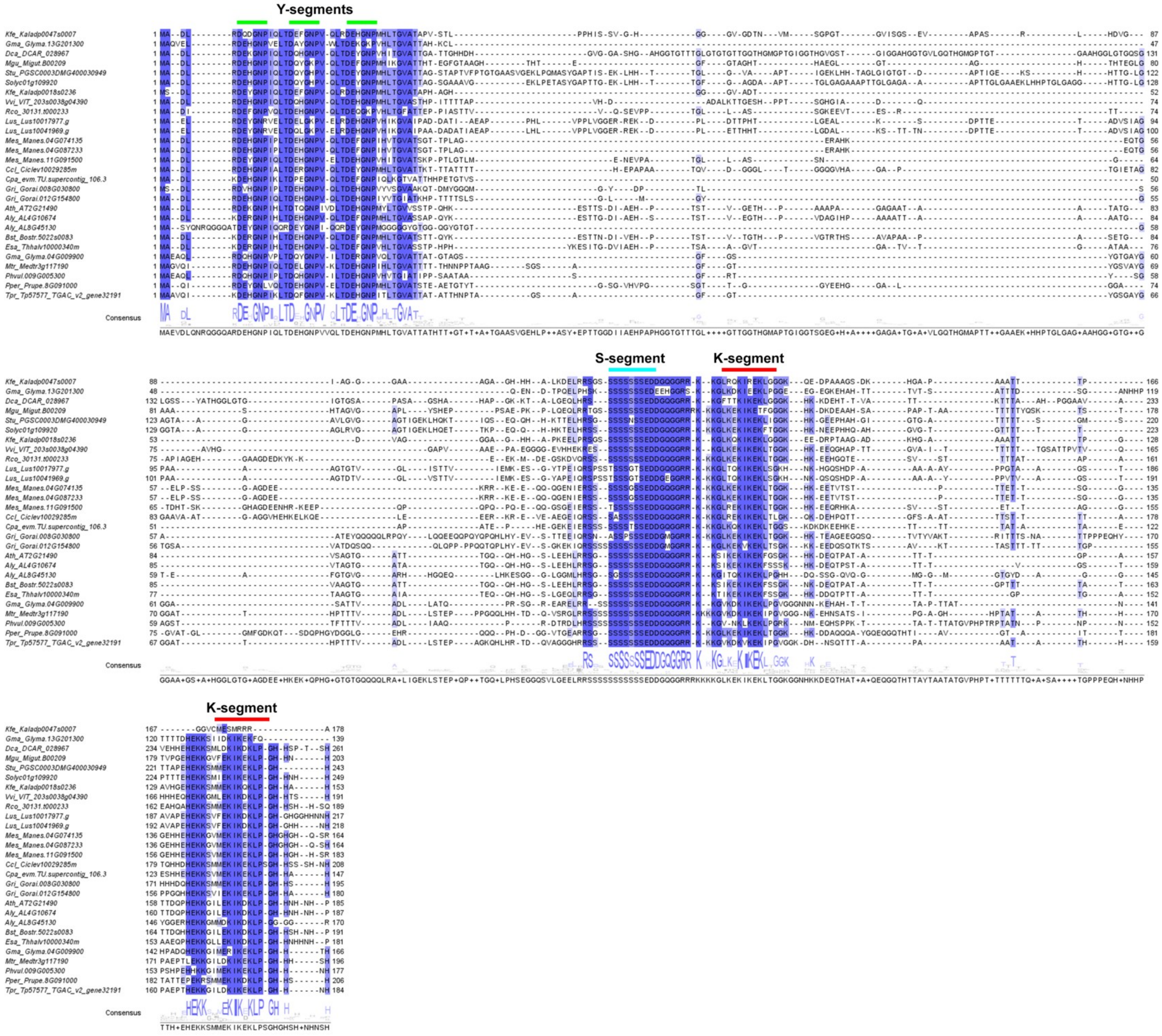


**Figure S6. Multiple sequences alignment of Y3SKn dehydrins.** Protein sequences of Y3SKn-DHNs from angiosperms were aligned with T-Coffee and visualised with Jalview. Structural segments are indicated and a consensus sequence is shown below the alignment.

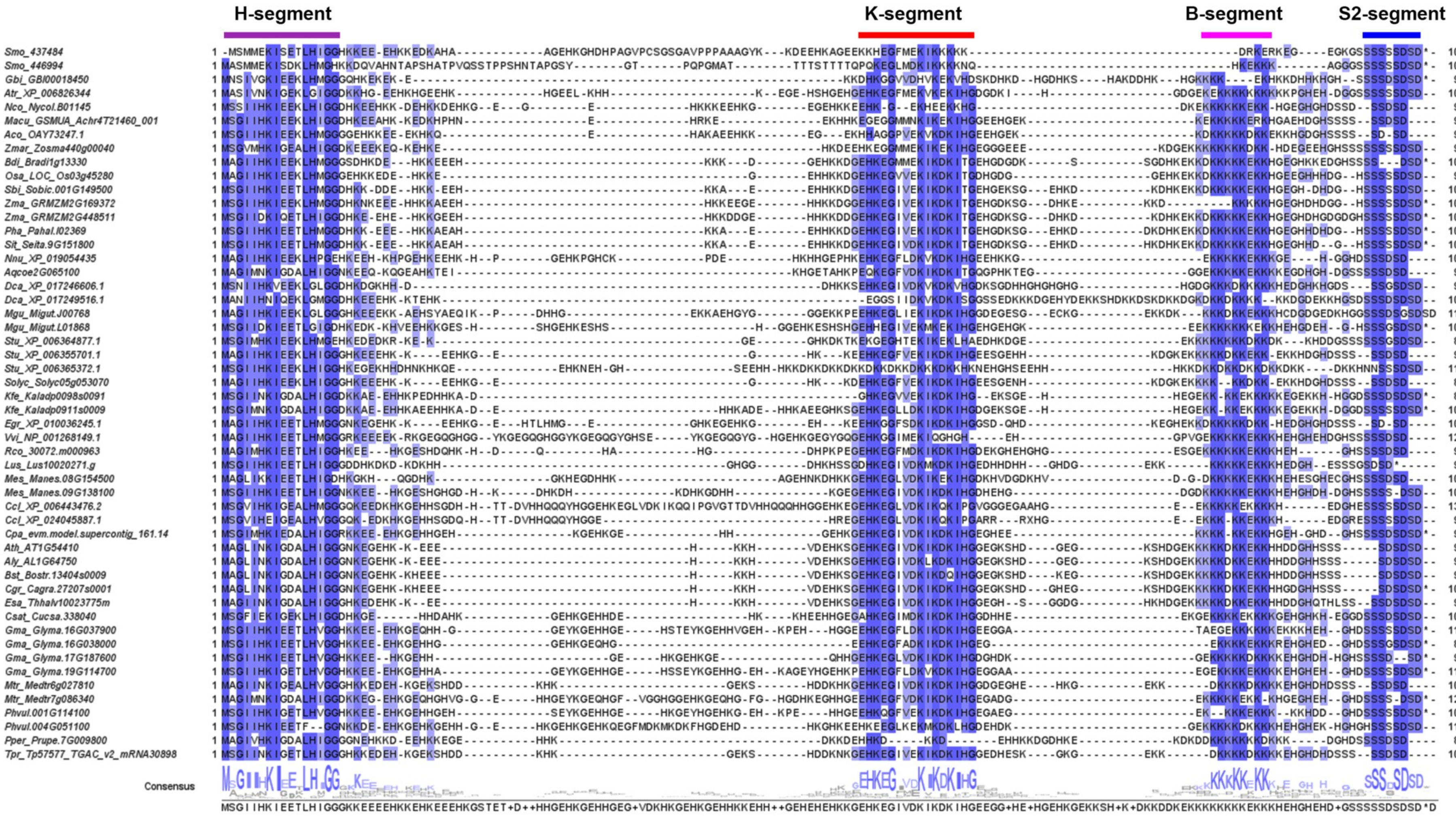


**Figure S7. Multiple sequence alignment of HSK-dehydrins.** Protein sequences of H-DHNs from vascular plants were aligned with Clustal Omega and visualised with Jalview. Structural segments are indicated and a consensus sequence is shown below the alignment. DHNs come from angiosperms except for proteins from *Selaginella moellendorffii* (Smo) and *Ginkgo biloba* (Gbi).

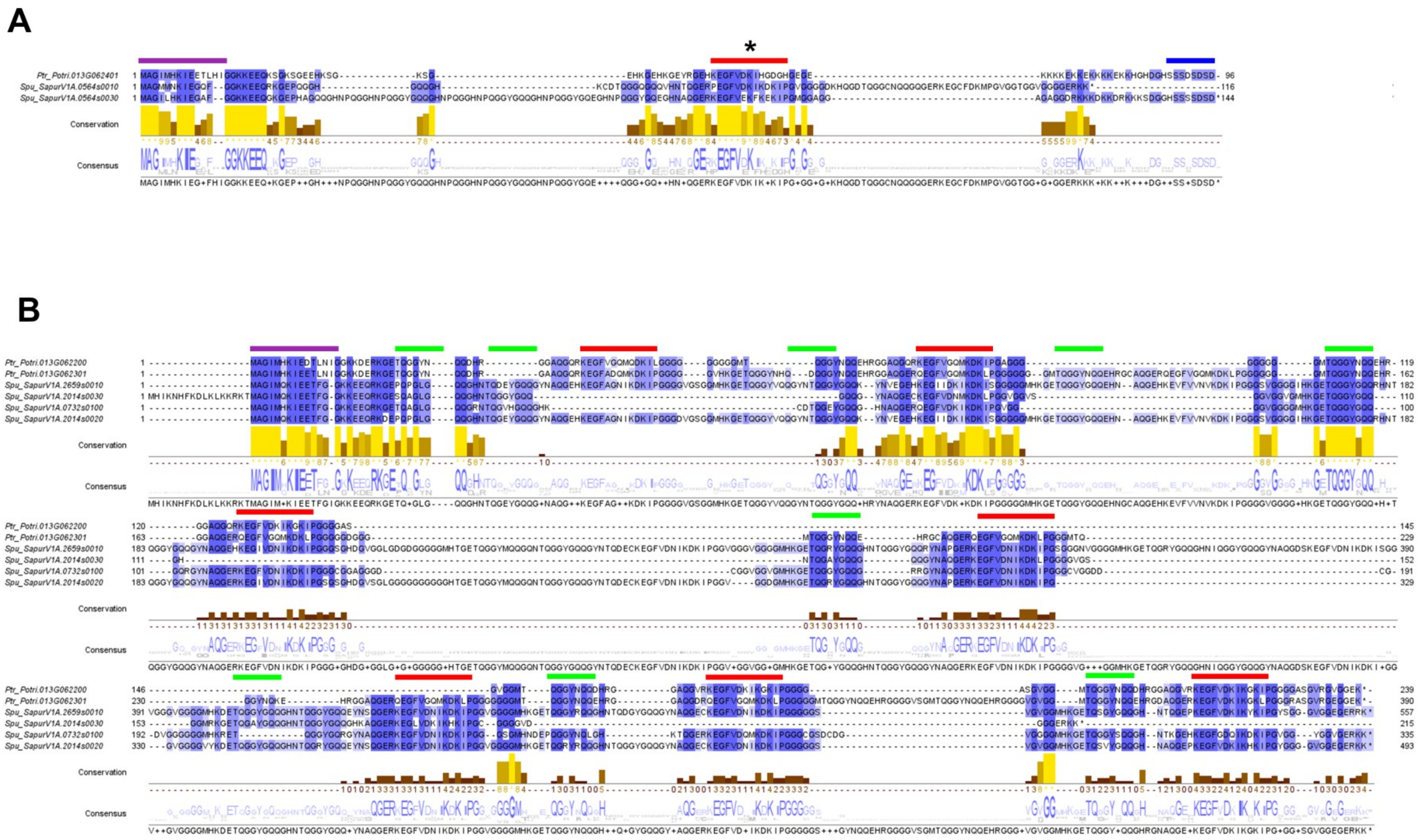


**Figure S8. Multiple sequence alignment of atypical H-DHNs from Malpighiales.** (**A**) Alignment of HKS-DHNs from *P. trichocarpa* and *S. purpurea* and a HS-DHN from *P. trichocarpa*. (**B**) Atypical H-DHNs with multiple K segments interspersed with Phi-segments. Segments are indicated by a colour code: H (purple), K (red), S2 (blue) and Phi (green). Sequences were aligned with Clustal Omega and visualised with Jalview.

**A**

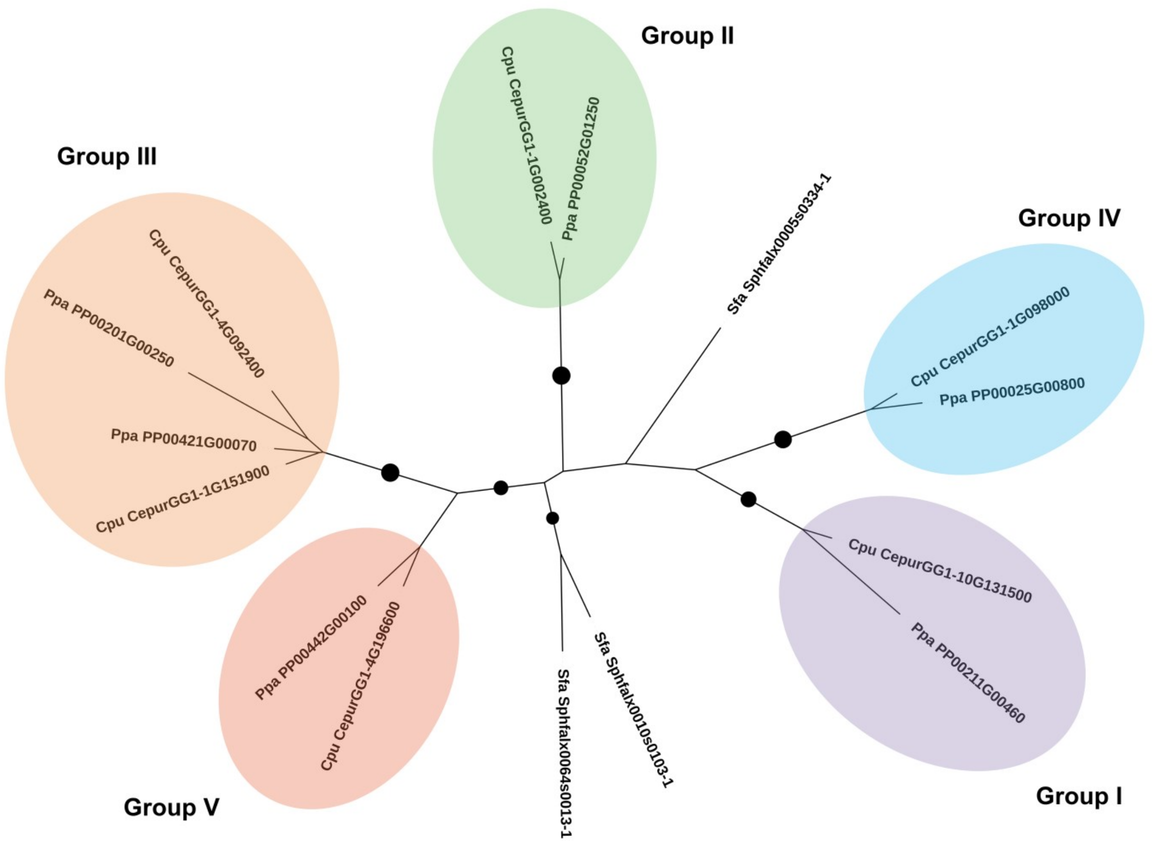


**B**

| | DHN orthologous groups | |
|---|---|---|
| | ***Physcomitrella patens*** | ***Ceratodon purpureus*** |
| **Group I** | PP00211G00460 (PpDHNA) | CepurGG1.10G131500 |
| **Group II** | PP00052G01250 (PpDHNB) | CepurGG1.1G002400 |
| **Group III** | PP00421G00070 (PpDHNC) | CepurGG1.1G151900 |
| | PP00201G00250 (PpDHND) | CepurGG1.4G092400 |
| **Group IV** | PP00025G00800 | CepurGG1.1G098000 |
| **Group V** | PP00442G00100 | CepurGG1.4G196600 |

**Figure S9. Evolutionary relationships of bryophyte dehydrins.** (**A**) Maximum-likelihood phylogenetic tree constructed with PhyML 3.0. Branches with bootstrap values over 90 are indicated with a circle. Note that DHN sequences from *P. patens* and *C. purpureus* form five homologous groups, while *S. fallax* DHNs are not grouped with the other sequences. (**B**) DHN sequences and homologous groups of *P. patens* and *C. purpureus*.

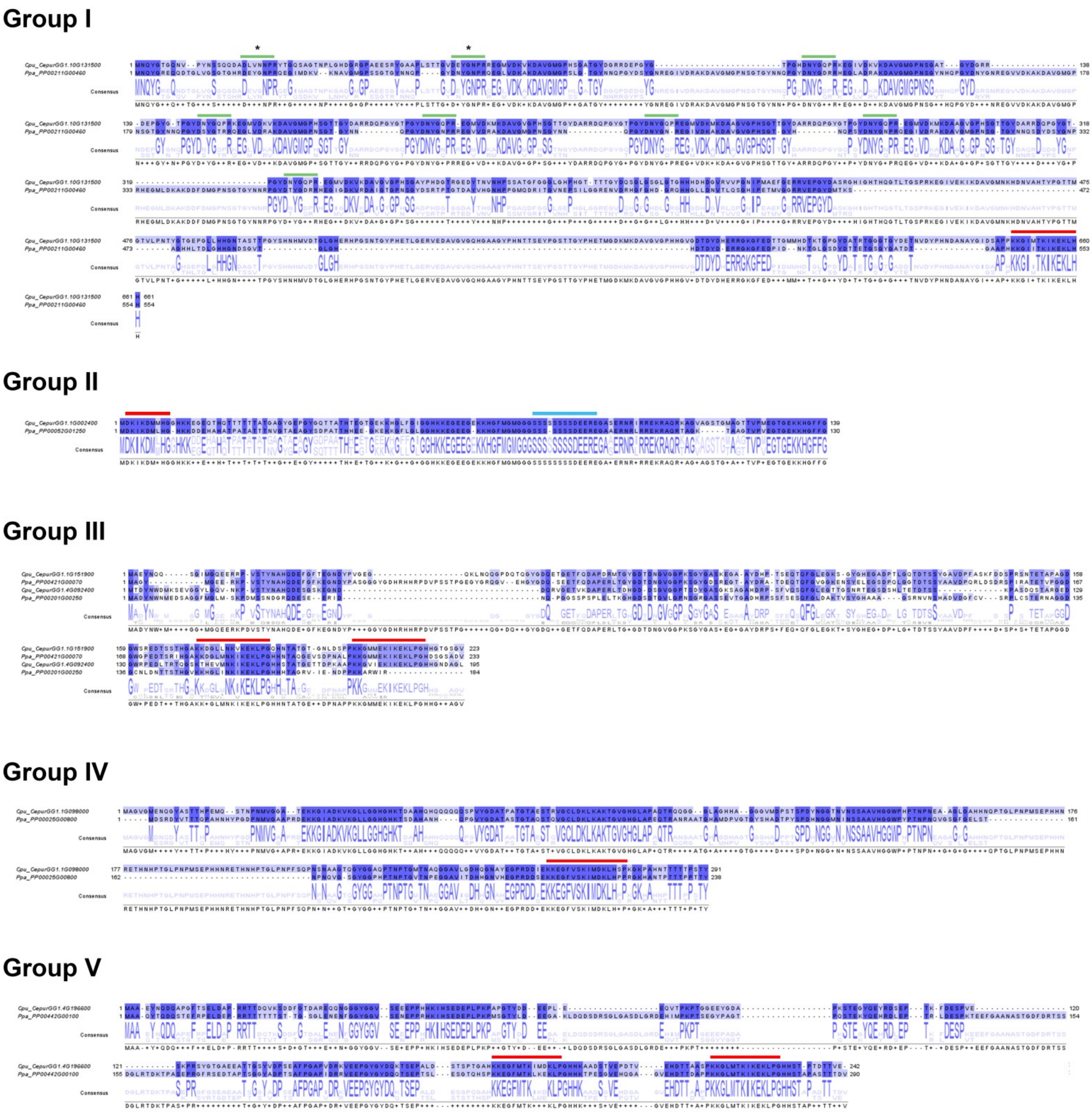


**Figure S10. Multiple sequence aligment of bryophyte DHNs.** Segments are indicated by a colour code: K (red), Y (green) and S (blue). Note that Group I has a Y8K structure; the Y-segments with an asterisk (*) have a sequence identical to the Y-segments of angiosperms (DEYGNP), while the others have a modified Y-segment (DNYGN/QP). Group II has a KS-structure, Group III and V have a K2-structure and Group IV a K-structure. Sequences were aligned with T-Coffee and visualised with Jalview.

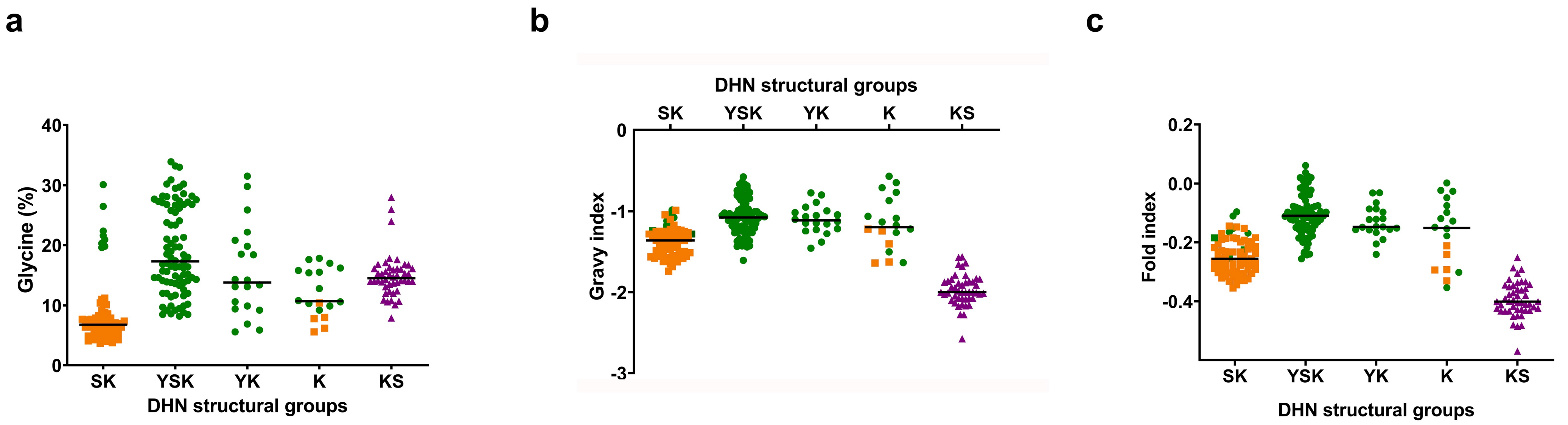

a
b
c
Glycine (%)
Gravy index
Fold index
DHN structural groups
SK
YSK
YK
K
KS
